# New mechanisms of radioiodide uptake revealed via a novel high throughput drug screening approach in thyroid cancer

**DOI:** 10.1101/2020.07.21.213967

**Authors:** Martin L. Read, Katie Brookes, Caitlin E.M. Thornton, Alice Fletcher, Mohammed Alshahrani, Rashida Khan, Hannah R. Nieto, Patricia Borges de Souza, Jamie R.M. Webster, Luke J. Alderwick, Kristien Boelaert, Vicki E. Smith, Christopher J. McCabe

## Abstract

New combinatorial drug strategies are urgently needed to improve radioiodide (RAI) uptake and efficiently ablate thyroid cancer cells, thereby addressing recurrent and metastatic disease. Cellular iodide uptake is accomplished solely by the sodium iodide symporter (NIS), but the complexity of NIS functional regulation and a lack of amenable high-throughput screening assays has impeded progress. We utilised mutated yellow fluorescent protein (YFP) as a surrogate biosensor of intracellular iodide for ∼1200 FDA-approved drugs, allowing us to appraise the impact of 73 leading compounds at 10 doses on ^125^I uptake in thyroid cancer cell lines. Subsequent mechanistic analysis suggests three predominant modes of drug action: Firstly, a number of drugs inhibited specific regulation of NIS function by the protein VCP. Secondly, some drugs enhanced transcriptional or post-transcriptional regulation of NIS expression. Thirdly, several drugs strongly implicated proteasomal degradation and the unfolded protein response in the cellular processing of NIS. Exploiting these mechanistic insights, multiple compounds gave striking increases in radioiodide uptake when combined with the drug SAHA. Importantly, our new drug combination strategies were also effective in human primary thyrocytes, suggesting they target endogenous NIS physiology. In patients with papillary thyroid cancer, genes involved in proteostasis were remarkably altered and predicted significantly worse outcome, but only in those patients who received RAI therapy. Collectively, we therefore propose a new model of intracellular NIS processing, and identify key nodes which may now be druggable in patients with aggressive thyroid cancer.

**SUMMARY:** Our data identify FDA-approved drugs that enhance radioiodide uptake outside of the canonical pathways of NIS processing, leading to a new mechanistic understanding of endogenous NIS function which is subverted in cancer.

## INTRODUCTION

For ∼80 years, radioiodide (RAI) has been the central post-surgical treatment for patients with differentiated thyroid cancer (DTC). However, around 25 percent of DTC patients have radioiodide-refractory thyroid cancer, being unable to uptake adequate RAI for effective therapeutic ablation (1). This is especially problematic for metastatic disease, which is associated with a life expectancy of 3–5 years, and hence defines a clear unmet medical need (1).

NIS is the sole transporter responsible for specific cellular iodide uptake (2); exploitation of its function remains the first – and most specifically targeted – internal radiation therapy: high-energy β-emitting ^131^I destroys remaining thyroid cells post-surgery, and targets metastases. The central mechanisms which underlie radioiodide-refractoriness are decreased levels of NIS expression and/or its diminished targeting to the plasma membrane (3). The regulation of NIS is principally accomplished via transcription factors, histone acetylation, post-translational modifications, hormonal signalling, and by iodide itself (4-10). These levels of control are generally disrupted in cancer by a plethora of mechanisms including altered histone acetylation and methylation of the NIS promoter (5, 6), distorted miRNA expression (11-13), increased oxidative stress (14), and changed growth factor signalling (15, 16), predominantly driven by the activation of oncogenes which directly stimulate the MAPK pathway. Thus in addressing repressed NIS function in thyroid cancer, this inherent multiplicity of regulation adds significantly to the complexity of potential therapeutic strategies.

Extensive pre-clinical and clinical studies have attempted to enhance NIS expression and function, focussing mainly on ‘re-differentiation agents’, which stimulate the expression of thyroid-specific genes including NIS. Because constitutive activation of the MAPK pathway in thyroid cancer frequently represses NIS expression and function, and is associated with decreased RAI uptake and poor patient prognosis, the majority of studies have focused on MAPK inhibitors (17-20). Other valuable approaches have addressed transcriptional and epigenetic alterations (21, 22) via the use of retinoids (23-25), PPARγ agonists (26, 27) and histone deacetylase (HDAC) inhibitors (28-31). However, while some drugs have convincingly been shown to enhance NIS expression in a subset of patients, fresh approaches are required to address the intricacy of altered NIS regulation in thyroid neoplasia.

For the present study we explored the hypothesis that we might identify new combinatorial drug approaches for improving NIS function by utilising NIS as a drug target in high throughput screening (HTS). HTS of NIS function has been reported before, but primarily to identify pollutants which disrupt NIS activity (32-34). Our study is therefore – to our knowledge – the first attempt to directly screen on a large scale novel pharmacological strategies capable of enhancing radioiodide uptake. By utilising a panel of predominantly FDA-approved drugs we sought to define new therapeutic possibilities which might ultimately systemically enhance thyroidal RAI uptake in patients. Thus we report a number of drugs which have never been identified before as critical enhancers of radioiodide uptake, and propose combinatorial approaches which markedly increase the efficacy of NIS function. Mechanistically, our findings provide new insight into the cellular processes which govern NIS function in normal and cancerous thyroid cells.

## MATERIALS AND METHODS

### Cell culture and stable cell line generation

Human thyroid carcinoma cell lines TPC-1 and 8505C, and human breast lines MDA-MB-231 and MCF7, were cultured as before (35), at low passage, authenticated by short tandem repeat analysis (NorthGene) and tested for Mycoplasma contamination (EZ-PCR; Geneflow). TPC-1 cells were kindly provided by Dr Rebecca Schweppe (University of Colorado, Denver). Other lines were obtained from DSMZ and ECACC.

Full-length human NIS cDNA (36) and the triple mutant enhanced YFP-H148Q/I152L/F46L [kindly provided by Prof Alan Verkman and Dr Peter Haggie (UCSF, San Francisco)] were cloned into pcDNA3.1. Stable NIS and/or YFP cells were generated using FACS single cell sorting (Supplementary Table S1). RAI uptake, Western blotting and NIS mRNA expression assays were used to monitor NIS activity as previously (35, 36). The expression of specific mRNAs was determined with a 7500 Real-time PCR system (Applied Biosystems). Opera Phenix™ live cell imaging (EvoTec, Germany) was used to validate YFP cell lines. Further details on nucleic acids and antibodies are provided (Supplementary Table S1).

### Human primary thyrocyte culture

The collection of normal human thyroid tissue was approved by the Local Research Ethics Committee, and subjects gave informed written consent. Human primary thyrocytes were isolated as previously (35, 37). Only cultures with positive TSH dose responses were progressed to cell experiments after 6 days.

### YFP-iodide assay based drug screening

High-throughput drug screening was performed via an automated biochemical screening platform using the Microlab STAR Liquid Handling Robotic System (Hamilton) integrated with a PHERAstar FS plate reader (BMG Labtech; 510 nm-550 nm optic module). Further details on YFP-iodide high-throughput assays and data analysis are provided (Supplementary Materials and Methods). 1200 approved compounds from the Prestwick Chemical Library (Birmingham Drug Discovery Facility; University of Birmingham) were screened (controls outlined in Supplementary Table S2).

### Gene expression data analyses

Normalized gene expression data and clinical information for papillary thyroid cancer (PTC) were downloaded from TCGA via cBioPortal and FireBrowse (THCA). Gene expression values were transformed as X=log_2_(X+1) where X represents the normalized fragments per kilobase transcript per million mapped reads (FPKM) values. In total, RNA-seq data for 59 normal thyroid and 501 PTC TCGA samples were analyzed. Further details on TCGA data analysis are provided (Supplementary Materials and Methods).

### Statistical analyses

Data were analysed using IBM SPSS, GraphPad Prism and Microsoft Excel (Supplementary Materials and Methods).

## RESULTS

### Drug screening identifies novel compounds which alter surrogate NIS activity

We first tested the known ability of mutated YFP to be fluorescently quenched by intracellular iodide (38, 39) in TPC-1 and 8505C thyroid cancer cells stably transfected with NIS and/or YFP. YFP fluorescence was quenched in a time and dose dependent manner (Fig. 1A and B), and we further confirmed time and dose dependent quenching via Opera Phenix™ system live cell imaging (Fig. 1C). We therefore adapted YFP-iodide sensing to a high-throughput screen of the Prestwick Chemical Library (∼1200 drugs; 95% FDA-approved), structured as indicated (Fig. 1A; Supplementary Fig. S1 and 2). Initial screening (*n*=2 entire screens in TPC-1-NIS-YFP cells; 10 μM) revealed that most drugs either repressed or had no impact on YFP (Fig. 1D and E). For 73 drugs which did reduce YFP fluorescence, we performed secondary screening at 10 different drug doses (Fig. 1F). Some drugs (e.g. clotrimazole (35)) showed increased activity over the whole 0.1 to 50 μM range of testing; others (e.g. terfenadine) were bi-phasic, whilst several (e.g. niflumic acid) showed a negative relationship at higher doses (Fig. 1F). Overall, adjusting for cell viability and subtracting YFP-only effects, we ranked the top drugs which specifically dimmed YFP fluorescence as a marker of cellular iodide uptake (Fig. 1G; Supplementary Fig. S3 and 4).

**Figure 1.**
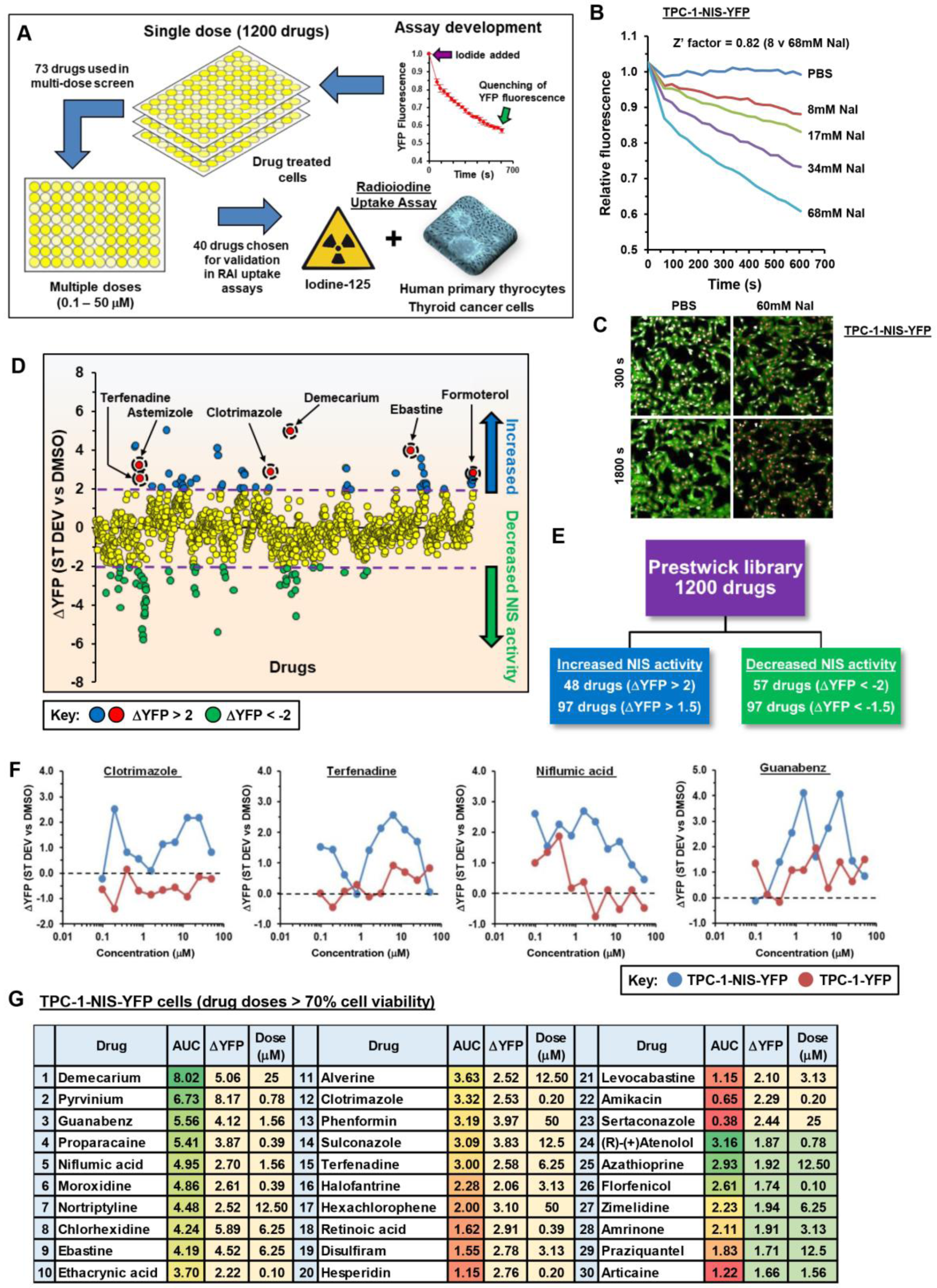
YFP-based biosensor strategy to screen drugs that increase intracellular iodide. (*A*) Overview of scheme used to identify novel compounds. (*B*) Validation of YFP-iodide assay in TPC-1-NIS-YFP cells using increasing NaI (0-68mM). (*C*) Opera Phenix™ live cell imaging of TPC-1-NIS-YFP cells treated with 60mM NaI. (*D*) Relative YFP fluorescence (ΔYFP) of TPC-1-NIS-YFP cells treated with the Prestwick Chemical Library (1200 drugs, 10 μM, 24 hours) versus DMSO. (*E*) Number of drugs showing maximal biological effect. (*F*) Representative drug dose-response YFP-iodide profiles in TPC-1-NIS-YFP vs TPC-1-YFP cells. (*G*) Top 30 drugs that increase intracellular iodide in TPC-1-NIS-YFP cells ranked on AUC and ΔYFP values (>70% cell viability).

### Testing the impact of HTS drugs on radioiodide uptake and NIS expression

We initially tested 38 of our top drugs in radioiodide uptake assays in TPC-1 cells stably expressing NIS; 16 of these gave significant dose-dependent increases in ^125^I uptake (Fig. 2A; Supplementary Fig. S5A and data not shown). To investigate potential mechanisms we firstly assessed their impact on NIS protein expression; 3 drugs significantly increased NIS protein levels and radioiodide uptake both in TPC-1 and 8505C cells stably expressing NIS (formoterol, pravastatin and disulfiram; Fig. 2B-E). Niflumic acid and zimelidine additionally increased radioiodide uptake in 8505Cs (Fig. 2F).

**Figure 2.**
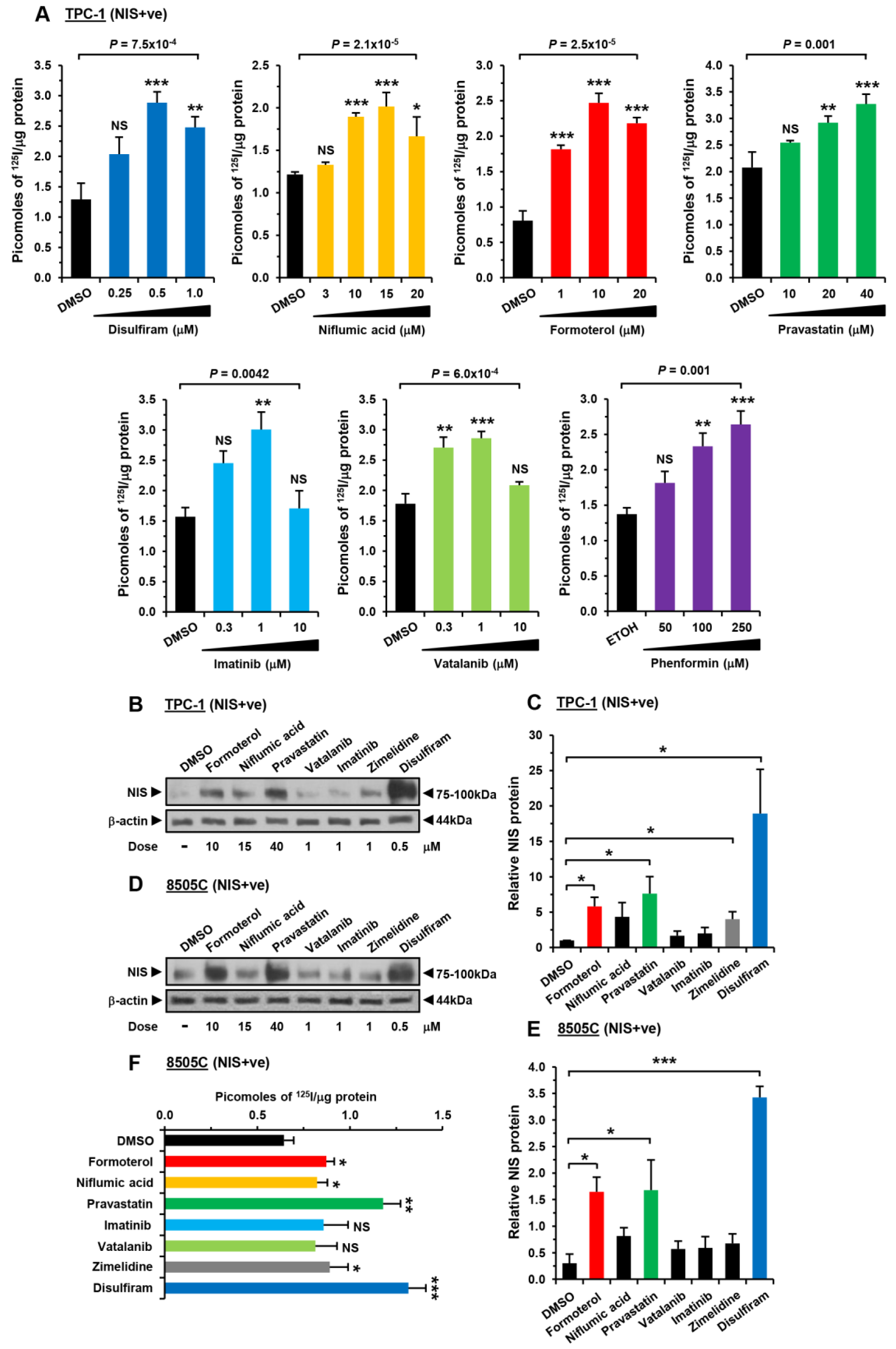
Validation of novel drugs enhancing RAI uptake and NIS expression. (*A*) RAI uptake of TPC-1 (NIS+ve) cells treated with drugs at indicated doses for 24 hours; one-way ANOVA followed by Dunnett’s post hoc test. (*B*) Western blot analysis of NIS expression levels in TPC-1 (NIS+ve) cell as described in (*A*). (*C*) Quantification of NIS from (*B*); *n*=3. (*D*-*F*) Same as (*A*-*C*) but RAI uptake and NIS expression in 8505C (NIS+ve) cells; unpaired t-test. NS, not significant; \**P*<0.05; \*\**P*<0.01; \*\*\**P*<0.001.

We next assessed the impact of our drugs on NIS mRNA expression. Although there were some cell-specific differences, only formoterol consistently induced NIS mRNA expression in both thyroid cancer cell lines (Supplementary Fig. S5B). Thus whereas formoterol acts transcriptionally, pravastatin and disulfiram are likely to exert posttranscriptional effects on NIS expression.

### Drug effects on endogenous NIS function

TPC-1 and 8505C thyroid cancer cells are likely to have inherently dysregulated cellular signalling (40). We therefore examined whether drugs identified from our HTS approaches would modulate endogenous NIS function in non-transformed human primary thyroid cells. Significant inductions of radioiodide uptake were apparent for – amongst others – niflumic acid, zimelidine, disulfiram, pravastatin and chloroquine (Fig. 3A; Supplementary Fig. S5C). Calculated EC50 values ranged from 65 nM (astemizole) to 66.8 μM (phenformin) (Fig. 3B). Thus disulfiram, pravastatin, formoterol, imatinib, niflumic acid, vatalanib, and chloroquine, which all induced radioiodide uptake in thyroid cancer cells stably expressing NIS, also induced significant ^125^I uptake in human primary cells, implicating them in the pathways which control normal physiological NIS function.

**Figure 3.**
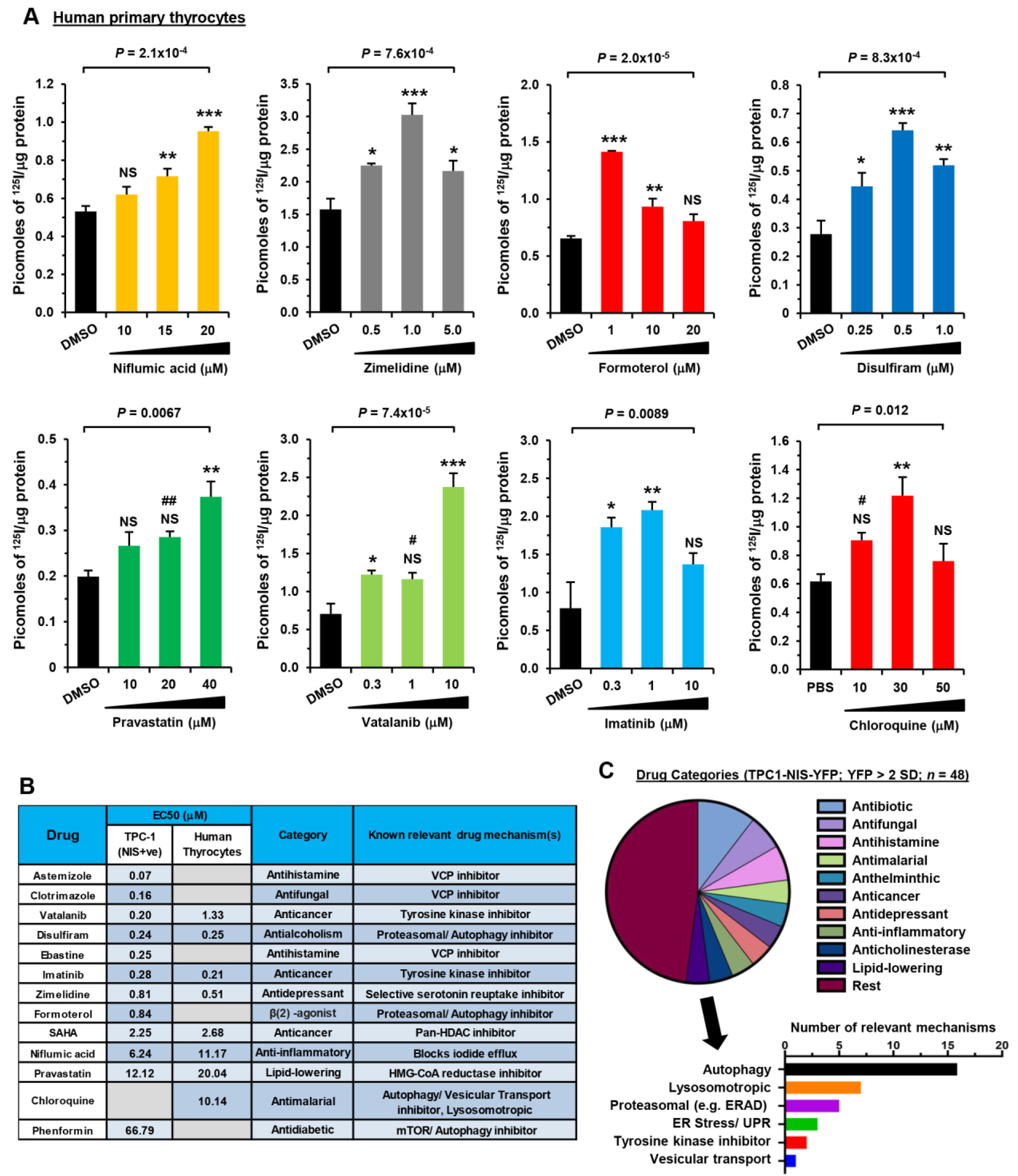
Significant drug-induced RAI uptake in human primary thyrocytes. (*A*) RAI uptake of primary human thyrocytes cells treated with drugs at indicated doses for 24 hours; one-way ANOVA followed by Dunnett’s post hoc test (NS, not significant; \**P*<0.05; \*\**P*<0.01; \*\*\**P*<0.001) or unpaired t-test (^#^*P*<0.05; ^##^*P*<0.01). (*B*) EC50 values for 13 leading candidate drugs in TPC-1 (NIS+ve) cells and/or primary human thyrocytes. (*C*) Categorisation of 48 drugs (pie chart) identified in primary screen in TPC-1-NIS-YFP cells with ΔYFP>2. Graph (below) shows number of relevant drug mechanisms.

Preliminary categorisation of YFP-screened data (ΔYFP>2; Fig. 1D) revealed a high proportion of drugs that modulate the proteostasis network (∼44%, 21/48 drugs; Supplementary Table S3), including key processes in protein homeostasis such as endoplasmic reticulum-associated protein degradation (ERAD) and autophagy (Fig. 3C). To predict in more detail which pathways of NIS processing might be targeted by our drug approaches, we next used Connectivity Map (CMAP) hierarchical clustering, which identified a major cluster strongly associated with genes implicated in the proteasome pathway and unfolded-protein response (Cluster 2; Supplementary Fig. S6). Most strikingly, 11/34 drugs in hierarchical cluster 2 were associated with loss of function of VCP, a critical component of multiple cellular functions including ERAD and autophagy (Supplementary Fig. S6). In support of this, 8/20 drugs (40%) validated for enhancing ^125^I uptake were associated with VCP loss of function, including ebastine and its metabolite carebastine (Supplementary Fig. S7).

### The mechanistic relationship between ebastine, carebastine and SAHA

As our most prominent drugs in terms of fold induction of NIS function were ebastine, carebastine and the histone deacetylase SAHA (Supplementary Table S4), we investigated their potential for mechanistic interaction. Thyroid cancers frequently show repression of NIS mRNA expression (11), and VCP is critical to the post-translational processing of NIS (35). We therefore addressed the hypothesis that SAHA treatment of thyroid cancer cells would induce NIS mRNA expression, as reported before (29, 41, 42), and that subsequently enhanced NIS protein levels might then ‘benefit’ from inhibition of VCP via repressed ER-associated degradation. However, a second hypothetical possibility also exists: SAHA inhibits histone deacetylase 6 (HDAC6), known to be critical to autophagosome–lysosome fusion. VCP is a functional binding partner of HDAC6, with VCP and HDAC6 able to cooperatively modulate the autophagic clearance of ubiquitylated proteins (43), which might include NIS. We therefore challenged these two potential hypotheses.

As expected, SAHA induced large dose-dependent increases in NIS protein in stable TPC-1 cells, accompanied by marked increases in radioiodide uptake (Supplementary Fig. S8). We previously identified ebastine as enhancing the localisation of NIS to the PM (35). Larger fold changes in radioiodide uptake were apparent in cells treated with ebastine (EBT) and SAHA than with SAHA alone (Fig. 4A; Supplementary Fig. S9A), predominantly reflecting increases in NIS protein expression. Combined SAHA treatment with VCP inhibition also revealed marked induction of ^125^I uptake for the putative VCP inhibitors astemizole and clotrimazole (Fig. 4A and B; Supplementary Fig. S9A) (44), which had demonstrated prominent increases in radioiodide uptake alone (Supplementary Table S4). Importantly, the specific VCP inhibitors (VCPi) ES-1 and NMS-873 confirmed potent combinatorial effects with SAHA, which reflected increases in NIS protein mediated via SAHA but also by the VCPi themselves (Supplementary Fig. S9B and C).

**Figure 4.**
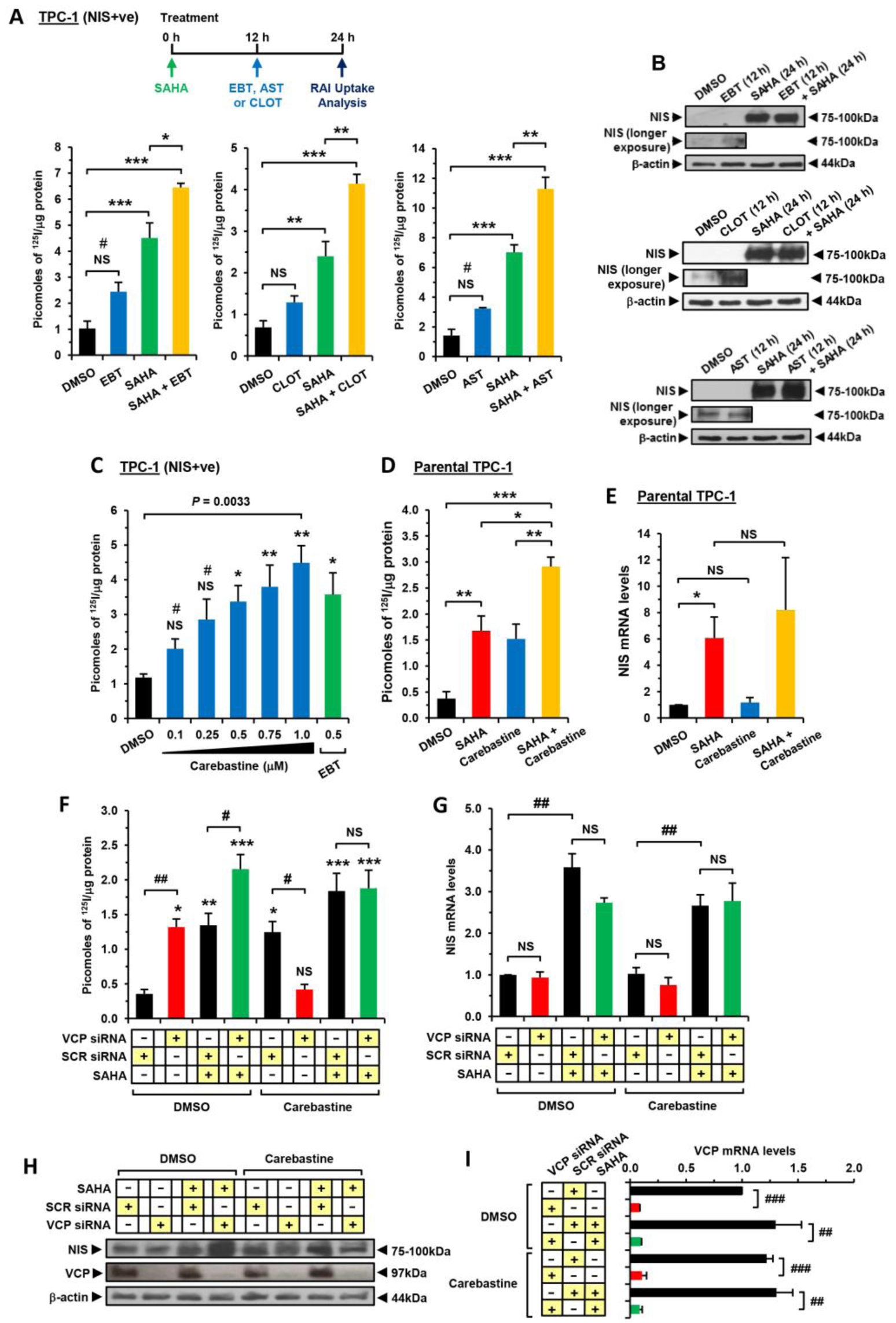
Enhanced RAI uptake by combining VCP inhibitors with SAHA. (*A*) RAI uptake and (*B*) NIS protein levels in TPC-1 (NIS+ve) cells treated with VCP inhibitors [ebastine (EBT), clotrimazole (CLOT) and astemizole (AST)], either alone or in combination with SAHA; one-way ANOVA followed by Tukey’s post hoc test. (*C*) RAI uptake of TPC-1 (NIS+ve) cells treated with carebastine at indicated doses versus EBT; one-way ANOVA followed by Dunnett’s post hoc test or unpaired t-test. (*D*) Same as (*A*) but parental TPC-1 cells treated with carebastine and/or SAHA. (*E*) NIS expression levels in parental TPC-1 cells as described in (*D*). (*F*) RAI uptake of TPC-1 (NIS+ve) cells following VCP-siRNA depletion and treatment with carebastine and/or SAHA; one-way ANOVA followed by Dunnett’s post hoc test or unpaired t-test. (*G*) NIS mRNA and (H) NIS protein levels in TPC-1 (NIS+ve) cells as described in (*F*). (*I*) VCP expression in VCP-siRNA ablated TPC-1 (NIS+ve) cells treated with carebastine and/or SAHA. NS, not significant; One-way ANOVAs: \**P*<0.05; \*\**P*<0.01; \*\*\**P*<0.001; Unpaired t-tests: ^#^*P*<0.05; ^##^*P*<0.01; ^###^*P*<0.001.

Whilst ebastine and SAHA were the most potent individual drugs in NIS-positive TPC-1 cells (Supplementary Table S4), ebastine is almost entirely converted to its active metabolite carebastine in vivo (45), and hence we also examined the functional relationship between carebastine and SAHA. Carebastine showed a dose dependent effect on radioiodide uptake in TPC-1 stable cells (Fig. 4C), with similar efficacy to its parent compound. To test whether the impact of carebastine is simply a thyroid-specific phenomenon, we utilised breast cancer MDA-MB-231 cells lentivirally expressing NIS. Carebastine was similarly potent in stimulating radioiodide uptake, confirming that it targets a generic mechanism of NIS processing (Supplementary Fig. S10A).

In combination with SAHA, carebastine worked additively in inducing radioiodide uptake in parental TPC-1s (Fig. 4D). Detectable radioiodide uptake in parental TPC-1 cells reflected SAHA induction of NIS mRNA expression, unlike carebastine, which did not impact NIS mRNA expression (Fig. 4E). Changes in NIS protein levels were difficult to detect due to the low NIS expression of thyroid cancer cells (40). However, VCP protein levels, which is a highly abundant cellular protein (46), were unaltered by either treatment (Supplementary Fig. S10B). In support of this, carebastine worked additively with SAHA to induce radioiodide uptake in breast cancer parental MDA-MB-231 and MCF7 cells without increases in NIS mRNA or VCP protein (Supplementary Fig. S10C to H). In control experiments, combining carebastine with the MEK inhibitor selumetinib did not enhance RAI uptake (Supplementary Fig. S10I).

Together, these combinatorial data suggest that rather than SAHA inhibition of HDAC6 acting in the same pathway as VCPi in blocking ERAD/autophagy, it is more likely that the additive effects of the 2 drugs arise from SAHA’s induction of NIS expression, and VCPi blockage of VCP-dependent pathways which control subsequent NIS protein processing (35). In further support, SAHA failed to alter the protein expression of the standard autophagy marker LC3B-II in TPC-1 cells (Supplementary Fig. 11A), and hence was not associated with an obvious modulation of autophagy.

To determine to what extent SAHA is dependent upon VCP, we depleted VCP, as previously (35). Both VCP siRNA and SAHA treatment increased radioiodide uptake, with a maximal effect when both were combined (Fig. 4F). However, carebastine was unable to induce ^125^I uptake when VCP was depleted, whereas SAHA retained its induction of NIS activity, confirming that carebastine’s impact is via VCP, but SAHA’s effect is not directly via the same pathway (Fig. 4F). NIS mRNA levels were not altered by VCP siRNA or carebastine treatment, but were altered by SAHA (Fig. 4G). Similarly, there was no induction of radioiodide uptake after carebastine treatment of VCP-ablated MDA-MB-231 (NIS+ve) cells (Supplementary Fig. S11B to E). In TPC-1s, NIS protein levels were increased by SAHA, which was potentiated in the presence of VCP knockdown (Fig. 4H). Meanwhile significant siRNA knockdown of VCP mRNA was unaffected by SAHA or carebastine (Fig. 4I). Thus, the drug combination of the HDACi SAHA and the VCPi carebastine appears to work principally by the induction of NIS expression via SAHA and the subsequent inhibition of VCP activity.

### Combinatorial use of chloroquine enhances RAI uptake

Given that VCP has well-established roles in protein processing and degradation, including via autophagy, we next investigated the role of the autophagy inhibitor chloroquine. Chloroquine roughly doubled the impact of SAHA on radioiodide uptake in TPC-1 cells stably expressing NIS (Fig. 5A), but this was independent of SAHA’s induction of NIS mRNA expression (Fig. 5B). Virtually identical data were apparent in 8505C cells (Fig. 5C). As a positive control, we inhibited BRAF^V600E^ activity via vemurafenib (VEM), which approximately doubled radioiodide uptake in BRAF^V600E^-mutant 8505C cells (Fig. 5D). Chloroquine did not enhance this further, whereas co-treatment of chloroquine and SAHA resulted in a much more potent 5.6-fold effect. Again, the predominant mechanistic theme here was the induction of NIS protein expression (Fig. 5E). In control experiments, we abrogated NIS function with the competitive inhibitor sodium perchlorate in TPC-1 and human primary cells (Fig. 5F; Supplementary Fig. S11F), proving that drug-induced radioiodide uptake was NIS-specific.

**Figure 5.**
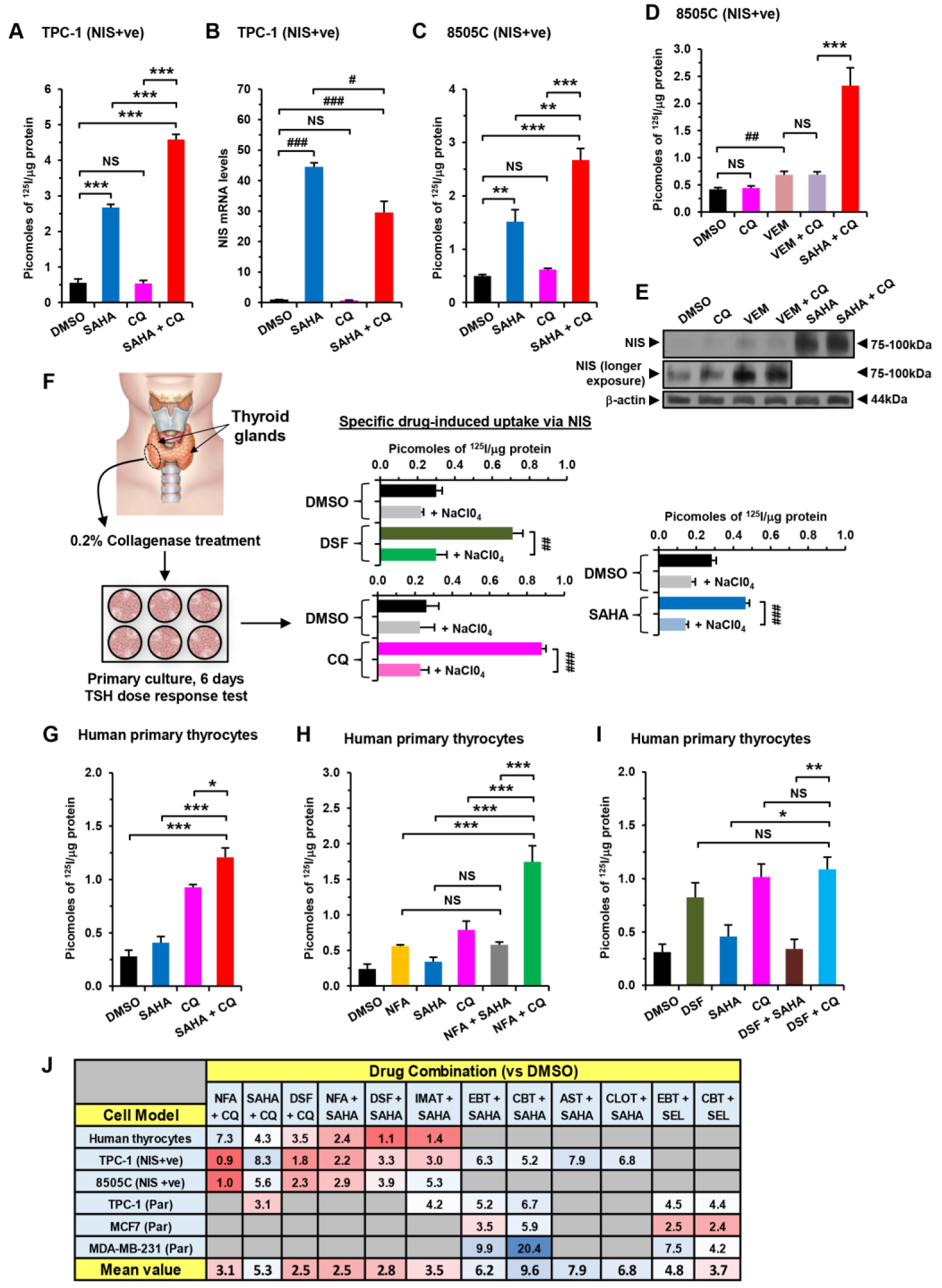
Chloroquine and SAHA robustly enhance radioiodide uptake. (*A*) RAI uptake and (*B*) NIS mRNA levels in TPC-1 (NIS+ve) cells treated with chloroquine (CQ) alone or in combination with SAHA; one-way ANOVA followed by Tukey’s post hoc test or unpaired t-test. (*C*) Same as (*A*) but RAI uptake in 8505C (NIS+ve) cells. (*D*) RAI uptake and (*E*) NIS protein levels in 8505C (NIS+ve) cells treated with vemurafenib (VEM)+CQ versus SAHA+CQ. (*F*) Schematic indicating protocol for obtaining human thyrocytes and validation of NIS specificity for drug-induced RAI uptake. Source: GoGraph. (*G-I*) RAI uptake in human primary thyrocytes treated with drugs alone [disulfiram (DSF), niflumic acid (NFA), SAHA and CQ] or in combination as indicated; one-way ANOVA followed by Tukey’s post hoc test. (*J*) RAI uptake (versus DMSO) with different drug combinations as indicated in thyroid and breast cells. NS, not significant; One-way ANOVAs: \**P*<0.05; \*\**P*<0.01; \*\*\**P*<0.001; Unpaired t-tests: ^#^*P*<0.05; ^##^*P*<0.01; ^###^*P*<0.001.

The impact of SAHA alone was relatively less effective in human primary thyrocytes (∼1.6-fold increase) than in TPC-1 cells (Fig. 5F and S5C; Supplementary Table S4), which likely reflected lessened epigenetic repression of NIS mRNA in non-transformed thyrocytes compared to transformed thyroid cells (5, 6). Importantly, our combination experiments did show that chloroquine and SAHA retained a robust and additive 4.3-fold effect on radioiodide uptake that was greater than chloroquine alone (Fig. 5G).

The highest fold induction of radioiodide uptake from combinatorial drug screening in non-transformed primary thyroid cells came from niflumic acid plus chloroquine (7.3-fold compared to DMSO; Fig. 5H). Interestingly, there was no additive effect from combining disulfiram with chloroquine (Fig. 5I). Full drug combination data (Fig. 5J) revealed that the best overall strategies which enhanced radioiodide uptake across at least 4 different cell types in multiple independent experiments were carebastine+SAHA (9.6-fold), ebastine+SAHA (6.2-fold), and chloroquine+SAHA (5.3-fold).

### Analysis of putative drug targets in human papillary thyroid cancer

Having identified drug combinations that robustly enhanced radioiodide uptake in thyroid cells, we next appraised the expression profiles and clinical relevance of gene drug targets in The Cancer Genome Atlas (TCGA) papillary thyroid cancer THCA dataset.

Firstly, we investigated a panel of 142 proteostasis genes in 4 functional categories (Supplementary Table S5), as many of our candidate drugs are known to target different arms of the proteostasis network, including proteasomal degradation, autophagy and trafficking, which might directly influence NIS function. Of significance, mRNA expression of a large proportion of proteasomal genes (Fig. 6A, 29/45) was higher in thyroid tumors versus normal tissue (Fig. 6B and Supplementary Fig. S12), whereas unfolded response protein (UPR) genes were generally down-regulated (5/6; Fig. 6A and Supplementary Fig. S13A and B). There was a striking correlation between proteasomal genes (Supplementary Fig. S14) that was significantly greater than in other categories (Fig. 6C and Supplementary Fig. S15A and B). VCP mRNA expression also positively correlated with ∼80% (35/44) proteasomal versus ∼34% (9/26) autophagy genes in THCA (*P*=0.001; Supplementary S15C to E).

**Figure 6.**
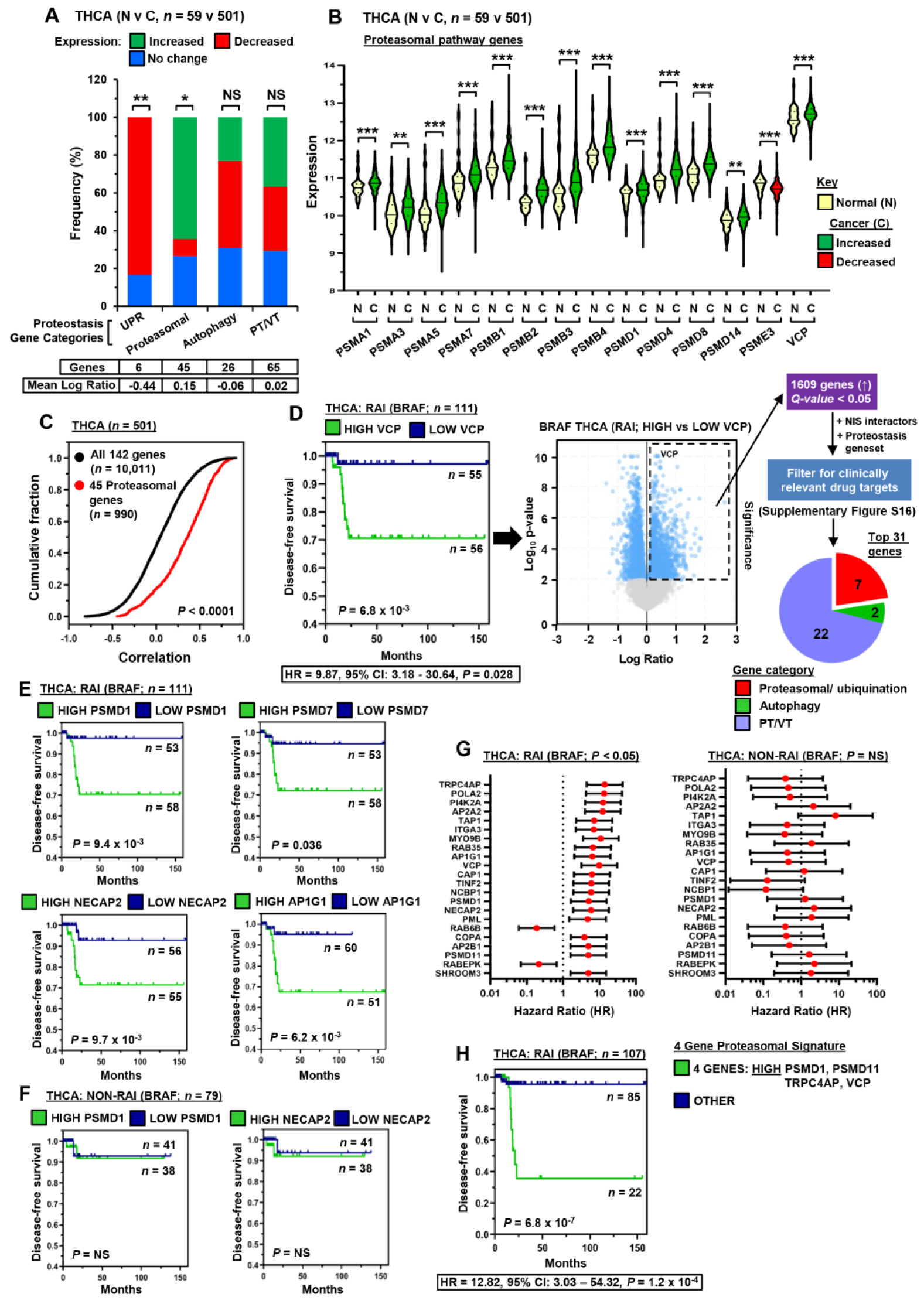
Dysregulated proteostasis gene expression linked to poorer patient survival. (*A*) 100% stacked bar chart comparing frequency of altered expression (N v C) in THCA TCGA for 142 proteostasis genes divided into unfolded protein response (UPR), proteasomal, autophagy, and protein transport/ vesicular trafficking (PT/VT) categories; Fisher’s exact test. (*B*) Violin plots showing representative gene expression profiles for 14 proteasomal genes as described in (*A*). (*C*) Cumulative frequency distribution plot comparing the correlation of 142 proteostasis versus 45 proteasomal genes as described in (*A*); Kolmogorov-Smirnov test. (*D*) Disease-free survival (DFS) for RAI-treated BRAF THCA patients with high (Q3Q4) versus low (Q1Q2) VCP expression (left), associated volcano plot (middle) and genes filtered for clinically relevant targets (right; *n*=31 proteostasis-related genes linked to reduced DFS), Examples shown: RAI-treated (*E*) and non-RAI treated patients (*F*). (*G*) Hazard ratios (HR)±95% CI for RAI-treated (left) versus non-RAI treated (right) patients stratified on median tumoral expression of 22 proteostasis genes. NS, not significant; \**P*<0.05. (*H*) Same as (*E*) but patients stratified on high tumoral expression of a 4-gene proteasomal signature.

Secondly, to identify the most clinically relevant proteostasis gene drug targets linked to NIS function and disease recurrence, we examined 347 genes (Supplementary Fig. S16) that included proteostasis-related genes from two additional sources: (i) NIS interactors recently detected by mass spectrometry (35) and (ii) VCP-associated genes in RAI-treated BRAF THCA patients with reduced disease-free survival (DFS) (Fig. 6D, Supplementary Fig. S17; Supplementary Table S5). Subsequent downstream analyses focussed on RAI-treated BRAF patients as this group represented a more uniform cohort with advanced disease staging characteristics (Supplementary Fig. S18).

In total the expression of 31 proteostasis genes was significantly associated with a higher rate of recurrence (i.e. lower DFS) in RAI-treated patients (Fig. 6E and Supplementary Table S6). Of these, 7 were proteasomal-related genes (e.g. *PSMD1*), 2 were autophagy-related genes (e.g. *ATG2A*), and the other 22 genes were associated with protein transport/vesicular trafficking (e.g. *NECAP2*) (Fig. 6E and Supplementary Fig. S19). Importantly, there was no significant difference in DFS of those patients who did not receive RAI treatment when stratified on median tumoral gene expression (Fig. 6F; Supplementary Fig. S20A and B; Supplementary Table S6). In addition, apart from *TAP1, CAP1* and *NCBP1*, clinical staging attributes for RAI treatment groups did not differ significantly when stratified for proteostasis gene expression (*P*=NS, 28 genes; Supplementary Table S6). Cox regression analysis further highlighted that higher tumoral expression of 22 proteostasis genes in RAI-treated THCA patients was linked with a significantly increased risk of recurrence (Hazard ratio (HR), *P*<0.05; Fig. 6G; Supplementary Table S6). In contrast, there was no corresponding difference in the risk of recurrence in THCA patients who were not treated with RAI (HR, *P*=NS; Fig. 6G; Supplementary Fig. S20C). Of particular significance, the risk of recurrence was even greater for RAI-treated patients using a gene signature based on 4 proteasomal-related genes, i.e. *VCP, PSMD1, PSMD11* and *TRPC4AP* (HR=12.82; *P*=1.2×10^−4^; Fig. 6H).

Collectively, we show that a panel of proteostasis genes are significantly dysregulated in PTC, with expression profiles which fit the repression of radioiodide uptake generally apparent in thyroid cancers. We hypothesise that dysregulation of these proteostasis genes results in reduced protein stability and trafficking of NIS to the PM. Further, we identify proteostasis genes associated with poorer survival characteristics in RAI-treated patients and which thus represent promising new drug targets in patients who are RAI-refractory. Based on our collective drug screening, experimental manipulations and clinical gene expression analyses, we propose a new model for the targetable steps of intracellular processing of NIS expression and function (Fig. 7).

**Figure 7.**
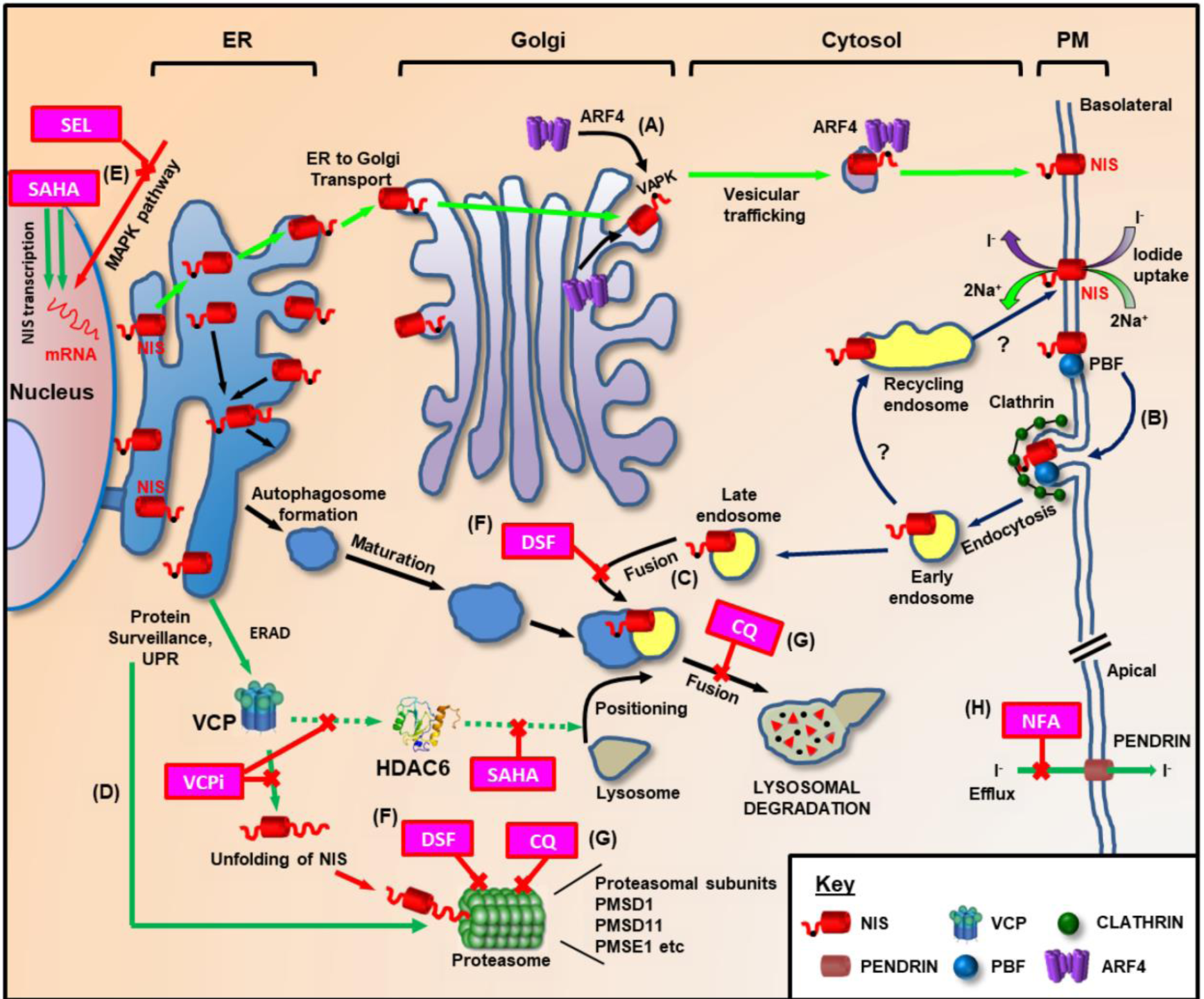
Hypothetical model of NIS intracellular processing revealed by drug screening. Given the consistently additive induction of ^125^I uptake when SAHA and VCPi are combined, we hypothesise that the predominant mechanism reflects enhanced NIS protein expression via NIS mRNA induction and reduced NIS degradation via ER-associated degradation (ERAD) and proteasomal degradation, rather than by targeting of dual VCP/HDAC6 function. These and previous data (35) suggest NIS maintains a delicate balance between protein synthesis, folding, assembly, trafficking and degradation. (*A*) ARF4 recognises the NIS C-terminus (black-spot) and promotes vesicular trafficking to the PM, where NIS is active. (*B*) NIS binds PBF, which has a YARF endocytosis motif and internalises NIS away from the PM in a clathrin-dependent process (36, 50). (*C*) Late endosomes merge with autophagosomes prior to fusing with lysosomes for degradation of protein cargoes. (*D*) Surveillance pathways target misfolded proteins for ERAD, as well as performing homeostatic regulation of correctly folded protein production (UPR). (*E*) SAHA stimulates transcriptional expression of NIS. Similarly, the MEK inhibitor Selumetinib (SEL) enhances NIS mRNA expression (13). (*F*) Disulfiram (DSF) inhibits autophagy (54) and proteasomal degradation (55, 57). (*G*) Chloroquine (CQ) acts ‘downstream’ of this, inhibiting the fusion of autophagosomes and lysosomes (52), and components of the proteasome (53). Dual inhibition of autophagy via DSF and CQ does not produce an additive impact on radioiodide uptake, suggesting they act in the same pathway. (*H*) Niflumic acid (NFA) reduces intracellular iodide efflux by inhibiting activity of the apical iodide transporter pendrin (58).

## DISCUSSION

HTS of NIS function has been reported before, but primarily for toxicology purposes. One previous study used HTS of the NIS promoter to detect drugs which increased NIS expression (47). However our study is the first attempt to directly ‘drug’ NIS function. Overall, we illustrate that pathways entirely separate to the canonical MAPK regulation of NIS exist, which can be drugged at multiple levels. In support of this, patients with PTC showed profound dysregulation of genes involved in proteasomal processing and the UPR, processes targeted by several of our drug strategies.

Clear repression of the UPR has not been reported before in thyroid cancer; we hypothesise that based on well-established mechanisms in other neoplasias, inhibition of the UPR is likely to reduce cellular apoptosis and may thus be permissive for tumor cell growth. In contrast, transcriptional activation of multiple components of the proteasome may enhance the degradation of NIS, leading to reduced radioiodide uptake and hence poorer clinical outcome.

Autophagy has been associated with NIS function in one previous report (48), though the precise mechanisms of how this might regulate NIS remain obscure. In thyroid cancers, markers of increased autophagy correlate with higher levels of plasma membranous NIS, and better clinical outcome (49). Our data clearly show that well established inhibitors of autophagy such as chloroquine and disulfiram induced dose-dependent increases in iodide uptake in untransformed primary thyroid cells, suggesting autophagy is critical to the central processing of NIS. SAHA and VCPi have been reported to interfere with autophagosome maturation, and hence it is possible that autophagosome maturation or processes downstream of it, rather than increased autophagy *per se*, is the more pertinent pathway to NIS drugability.

An alternative pathway could involve endosomes recycling NIS away from the plasma membrane, a process previously described by ourselves and others (35, 36, 50). Once endocytic vesicles become uncoated they fuse with early endosomes and mature into late endosomes before fusing with lysosomes. Endosomes are known to fuse with autophagosomes as part of lysosomal degradation. Autophagy selectively degrades targets and contributes to intracellular homeostasis (51). Drugs which interfere with autophagosome initiation (SAHA), maturation (disulfiram, VCPi) or fusion with lysosomes (chloroquine), could all potentially result in NIS-bearing endosomes not being degraded, and subsequently increased intracellular NIS protein. However, these NIS-containing endosomes would then have to return to the PM, and the notional mechanism for this is not clear.

SAHA is a pan-HDAC inhibitor, and amongst other HDACs inhibits HDAC6, required for autophagosome–lysosome fusion. We considered it possible that SAHA may work by inhibiting HDAC6, a functional binding partner of VCP. Against this possibility, SAHA was clearly associated with a rapid induction of NIS mRNA expression which closely mirrored its impact on radioiodide uptake, and also failed to induce canonical markers of autophagy.

New drug strategies, such as combining BRAF or MEK inhibitors with pan-PI3K inhibitors, are beginning to show pre-clinical promise in thyroid cancer (19). However, we propose that pathways outside those canonical signalling pathways may be capable of augmenting NIS expression, processing and trafficking to the PM to boost the efficacy of radioisotope treatment. In patients with PTC, perhaps the most striking observation was the dysregulation of multiple genes involved in proteasomal degradation, and the associated correlation with DFS, but only in those patients treated with RAI. Whilst this represents a circumstantial link between enhanced proteasomal degradation and reduced NIS function, it hints that brief therapeutic targeting of the proteasome at the time of RAI treatment might enhance RAI avidity. Of our drugs, chloroquine has a well described role in inhibiting the fusion of autophagosomes and lysosomes (52), but can also inhibit components of the proteasome (53). Disulfiram has recently been associated with the accumulation of ubiquitylated proteins in oesophageal squamous cell carcinoma (54), and also inhibits the proteasome (55). The relative brevity of treatment with drugs such as chloroquine and disulfiram would obviate some of the issues which have dogged longer term clinical trials into proteasome inhibitors, perhaps providing a therapeutic window in which NIS proteasomal degradation is inhibited, prior to ^131^I administration.

It is difficult to speculate which of our 3 proposed drug combinations (ebastine+SAHA, carebastine+SAHA or chloroquine+SAHA) might represent realistic human drug strategies in thyroid cancer. All the drugs of this study have favourable side effect profiles. Chloroquine’s standard dose is 250 mg/day and has radiosensitising properties (56), which might enhance the efficacy of radioiodide uptake independent of its effect on NIS. In cell experiments the maximal effective concentration of chloroquine was relatively high at ∼50 μM, but it is known to cross cell membranes easily and to accumulate within cell structures over time. SAHA can be taken at a dose of up to 400 mg/day giving a serum concentration of around 1.2 μM. Whilst ebastine and carebastine were consistently effective *in vitro*, at EC50s of around 250 nM, they are generally only taken at doses of ∼20 mg/day. Therefore we would propose that all of these combinations merit preclinical screening for definitive insight into time and dose dependent efficacies, as well as definitive mechanisms of drug action.

Taken together, we report the first large scale screen aimed at enhancing NIS activity. Our findings confirm that the cellular processing of NIS involves VCP, and suggest that given the marked dysregulation of proteasomal subunit genes in PTC patients treated with RAI who had poorer clinical outcomes, the proteasome is an integral regulatory pathway of NIS activity which is dysregulated in thyroid cancer. Combinatorial experiments reveal that >5-fold inductions of ^125^I uptake are routinely possible *in vitro*, opening the way for new *in vivo* strategies to enhance radioiodide uptake in human thyroid cancer.

## ACKNOWLEDGEMENTS

This work was supported by the MRC, Wellcome Trust, Get A-Head and The University of Birmingham. We acknowledge the contribution of the Human Biomaterials Resource Centre (UoB) and thank Evotec for help with the Opera Phenix™ system.

## REFERENCES

1 Schlumberger M, Brose M, Elisei R, Leboulleux S, Luster M, Pitoia F, et al. Definition and management of radioactive iodine-refractory differentiated thyroid cancer. Lancet Diabetes Endocrinol. 2014;2:356–8.

2 Dai G, Levy O, Carrasco N. Cloning and characterization of the thyroid iodide transporter. Nature. 1996;379:458–60.

3 Spitzweg C, Harrington KJ, Pinke LA, Vile RG, Morris JC. Clinical review 132: The sodium iodide symporter and its potential role in cancer therapy. J Clin Endocrinol Metab. 2001;86:3327–35.

4 Ravera S, Reyna-Neyra A, Ferrandino G, Amzel LM, Carrasco N. The Sodium/Iodide Symporter (NIS): Molecular Physiology and Preclinical and Clinical Applications. Annu Rev Physiol. 2017;79:261–89.

5 Zhang Z, Liu D, Murugan AK, Liu Z, Xing M. Histone deacetylation of NIS promoter underlies BRAF V600E-promoted NIS silencing in thyroid cancer. Endocr Relat Cancer. 2014;21:161–73.

6 Passon N, Puppin C, Lavarone E, Bregant E, Franzoni A, Hershman JM, et al. Cyclic AMP-response element modulator inhibits the promoter activity of the sodium iodide symporter gene in thyroid cancer cells. Thyroid. 2012;22:487–93.

7 Kogai T, Sajid-Crockett S, Newmarch LS, Liu YY, Brent GA. Phosphoinositide-3-kinase inhibition induces sodium/iodide symporter expression in rat thyroid cells and human papillary thyroid cancer cells. J Endocrinol. 2008;199:243–52.

8 Eskandari S, Loo DD, Dai G, Levy O, Wright EM, Carrasco N. Thyroid Na+/I-symporter. Mechanism, stoichiometry, and specificity. J Biol Chem. 1997;272:27230–8.

9 Dohan O, De l, V, Paroder V, Riedel C, Artani M, Reed M, et al. The sodium/iodide Symporter (NIS): characterization, regulation, and medical significance. Endocr Rev. 2003;24:48–77.

10 Arriagada AA, Albornoz E, Cecilia Opazo M, Becerra A, Vidal G, Fardella C, et al. Excess Iodide Induces an Acute Inhibition of the Sodium/Iodide Symporter in thyroid rat cells by Increasing Reactive Oxygen Species. Endocrinol. 2015;156:1540–51.

11 Cancer Genome Atlas Research N. Integrated genomic characterization of papillary thyroid carcinoma. Cell. 2014;159:676–90.

12 Riesco-Eizaguirre G, Wert-Lamas L, Perales-Paton J, Sastre-Perona A, Fernandez LP, Santisteban P. The miR-146b-3p/PAX8/NIS Regulatory Circuit Modulates the Differentiation Phenotype and Function of Thyroid Cells during Carcinogenesis. Cancer Res. 2015;75:4119–30.

13 Wachter S, Wunderlich A, Greene BH, Roth S, Elxnat M, Fellinger SA, et al. Selumetinib Activity in Thyroid Cancer Cells: Modulation of Sodium Iodide Symporter and Associated miRNAs. Int J Mol Sci. 2018;19:2077.

14 Azouzi N, Cailloux J, Cazarin JM, Knauf JA, Cracchiolo J, Al Ghuzlan A, et al. NADPH Oxidase NOX4 Is a Critical Mediator of BRAF(V600E)-Induced Downregulation of the Sodium/Iodide Symporter in Papillary Thyroid Carcinomas. Antioxid Redox Signal. 2017;26:864–77.

15 Garcia B, Santisteban P. PI3K is involved in the IGF-I inhibition of TSH-induced sodium/iodide symporter gene expression. Mol Endocrinol. 2002;16:342–52.

16 Riesco-Eizaguirre G, Rodriguez I, De la Vieja A, Costamagna E, Carrasco N, Nistal M, et al. The BRAFV600E oncogene induces transforming growth factor beta secretion leading to sodium iodide symporter repression and increased malignancy in thyroid cancer. Cancer Res. 2009;69:8317–25.

17 Buffet C, Wassermann J, Hecht F, Leenhardt L, Dupuy C, Groussin L, et al. Redifferentiation of radioiodine-refractory thyroid cancers. Endocr Relat Cancer. 2020; ERC-19-0491.R2.

18 Dunn LA, Sherman EJ, Baxi SS, Tchekmedyian V, Grewal RK, Larson SM, et al. Vemurafenib Redifferentiation of BRAF Mutant, RAI-Refractory Thyroid Cancers. J Clin Endocrinol Metab. 2019;104:1417–28.

19 Nagarajah J, Le M, Knauf JA, Ferrandino G, Montero-Conde C, Pillarsetty N, et al. Sustained ERK inhibition maximizes responses of BrafV600E thyroid cancers to radioiodine. J Clin Invest. 2016;126:4119–24.

20 Ho AL, Grewal RK, Leboeuf R, Sherman EJ, Pfister DG, Deandreis D, et al. Selumetinib-enhanced radioiodine uptake in advanced thyroid cancer. N Engl J Med. 2013;368:623–32.

21 Mancikova V, Buj R, Castelblanco E, Inglada-Perez L, Diez A, de Cubas AA, et al. DNA methylation profiling of well-differentiated thyroid cancer uncovers markers of recurrence free survival. Int J Cancer. 2014;135:598–610.

22 Kitazono M, Robey R, Zhan Z, Sarlis NJ, Skarulis MC, Aikou T, et al. Low concentrations of the histone deacetylase inhibitor, depsipeptide (FR901228), increase expression of the Na(+)/I(-) symporter and iodine accumulation in poorly differentiated thyroid carcinoma cells. J Clin Endocrinol Metab. 2001;86:3430–5.

23 Schmutzler C, Schmitt TL, Glaser F, Loos U, Kohrle J. The promoter of the human sodium/iodide-symporter gene responds to retinoic acid. Mol Cell Endocrinol. 2002;189:145–55.

24 Kogai T, Kanamoto Y, Che LH, Taki K, Moatamed F, Schultz JJ, et al. Systemic retinoic acid treatment induces sodium/iodide symporter expression and radioiodide uptake in mouse breast cancer models. Cancer Res. 2004;64:415–22.

25 Simon D, Kohrle J, Schmutzler C, Mainz K, Reiners C, Roher HD. Redifferentiation therapy of differentiated thyroid carcinoma with retinoic acid: basics and first clinical results. Exp Clin Endocrinol Diabetes. 1996;104 Suppl 4:13–5.

26 Kebebew E, Peng M, Reiff E, Treseler P, Woeber KA, Clark OH, et al. A phase II trial of rosiglitazone in patients with thyroglobulin-positive and radioiodine-negative differentiated thyroid cancer. Surgery. 2006;140:960-6; discussion 6-7.

27 Frohlich E, Brossart P, Wahl R. Induction of iodide uptake in transformed thyrocytes: a compound screening in cell lines. Eur J Nucl Med Mol Imaging. 2009;36:780–90.

28 Pugliese M, Fortunati N, Germano A, Asioli S, Marano F, Palestini N, et al. Histone deacetylase inhibition affects sodium iodide symporter expression and induces 131I cytotoxicity in anaplastic thyroid cancer cells. Thyroid. 2013;23:838–46.

29 Wachter S, Damanakis AI, Elxnat M, Roth S, Wunderlich A, Verburg FA, et al. Epigenetic Modifications in Thyroid Cancer Cells Restore NIS and Radio-Iodine Uptake and Promote Cell Death. J Clin Med. 2018;7:61.

30 Fu H, Cheng L, Jin Y, Cheng L, Liu M, Chen L. MAPK Inhibitors Enhance HDAC Inhibitor-Induced Redifferentiation in Papillary Thyroid Cancer Cells Harboring BRAF (V600E): An In Vitro Study. Mol Ther Oncolytics. 2019;12:235–45.

31 Massimino M, Tirro E, Stella S, Frasca F, Vella V, Sciacca L, et al. Effect of Combined Epigenetic Treatments and Ectopic NIS Expression on Undifferentiated Thyroid Cancer Cells. Anticancer Res. 2018;38:6653–62.

32 Agretti P, Dimida A, De Marco G, Ferrarini E, Rodriguez Gonzalez JC, Santini F, et al. Study of potential inhibitors of thyroid iodide uptake by using CHO cells stably expressing the human sodium/iodide symporter (hNIS) protein. J Endocrinol Invest. 2011;34:170–4.

33 Hallinger DR, Murr AS, Buckalew AR, Simmons SO, Stoker TE, Laws SC. Development of a screening approach to detect thyroid disrupting chemicals that inhibit the human sodium iodide symporter (NIS). Toxicol In Vitro. 2017;40:66–78.

34 Buckalew AR, Wang J, Murr AS, Deisenroth C, Stewart WM, Stoker TE, et al. Evaluation of potential sodium-iodide symporter (NIS) inhibitors using a secondary Fischer rat thyroid follicular cell (FRTL-5) radioactive iodide uptake (RAIU) assay. Arch Toxicol. 2020;94:873–85.

35 Fletcher A, Read ML, Thornton CEM, Larner DP, Poole VL, Brookes K, et al. Targeting Novel Sodium Iodide Symporter Interactors ADP-Ribosylation Factor 4 and Valosin-Containing Protein Enhances Radioiodine Uptake. Cancer Res. 2020;80:102–15.

36 Smith VE, Read ML, Turnell AS, Watkins RJ, Watkinson JC, lewy gd, et al. A novel mechanism of sodium iodide symporter repression in differentiated thyroid cancer. J Cell Sci. 2009;122:3393–402.

37 Read ML, Fong JC, Modasia B, Fletcher A, Imruetaicharoenchoke W, Thompson RJ, et al. Elevated PTTG and PBF predicts poor patient outcome and modulates DNA damage response genes in thyroid cancer. Oncogene. 2017;36:5296–308.

38 Rhoden KJ, Cianchetta S, Duchi S, Romeo G. Fluorescence quantitation of thyrocyte iodide accumulation with the yellow fluorescent protein variant YFP-H148Q/I152L. Anal Biochem. 2008;373:239–46.

39 Di Bernardo J, Iosco C, Rhoden KJ. Intracellular anion fluorescence assay for sodium/iodide symporter substrates. Anal Biochem. 2011;415:32–8.

40 Landa I, Pozdeyev N, Korch C, Marlow LA, Smallridge RC, Copland JA, et al. Comprehensive Genetic Characterization of Human Thyroid Cancer Cell Lines: A Validated Panel for Preclinical Studies. Clin Cancer Res. 2019;25:3141–51.

41 Puppin C, D’Aurizio F, D’Elia AV, Cesaratto L, Tell G, Russo D, et al. Effects of histone acetylation on sodium iodide symporter promoter and expression of thyroid-specific transcription factors. Endocrinol. 2005;146:3967–74.

42 Cheng W, Liu R, Zhu G, Wang H, Xing M. Robust Thyroid Gene Expression and Radioiodine Uptake Induced by Simultaneous Suppression of BRAF V600E and Histone Deacetylase in Thyroid Cancer Cells. J Clin Endocrinol Metab. 2016;101:962–71.

43 Boyault C, Gilquin B, Zhang Y, Rybin V, Garman E, Meyer-Klaucke W, et al. HDAC6-p97/VCP controlled polyubiquitin chain turnover. EMBO J. 2006;25:3357–66.

44 Segura-Cabrera A, Tripathi R, Zhang X, Gui L, Chou TF, Komurov K. A structure- and chemical genomics-based approach for repositioning of drugs against VCP/p97 ATPase. Sci Rep. 2017;7:44912.

45 Fujii T, Matsumoto S, Hatoyama T, Miyazaki H. Studies on the first-pass metabolism of ebastine in rats. Arzneimittelforschung. 1997;47:949–53.

46 Meyer H, Weihl CC. The VCP/p97 system at a glance: connecting cellular function to disease pathogenesis. J Cell Sci. 2014;127:3877–83.

47 Oh JM, Kalimuthu S, Gangadaran P, Baek SH, Zhu L, Lee HW, et al. Reverting iodine avidity of radioactive-iodine refractory thyroid cancer with a new tyrosine kinase inhibitor (K905-0266) excavated by high-throughput NIS (sodium iodide symporter) enhancer screening platform using dual reporter gene system. Oncotarget. 2018;9:7075–87.

48 Chai W, Ye F, Zeng L, Li Y, Yang L. HMGB1-mediated autophagy regulates sodium/iodide symporter protein degradation in thyroid cancer cells. J Exp Clin Cancer Res. 2019;38:325.

49 Plantinga TS, Tesselaar MH, Morreau H, Corssmit EP, Willemsen BK, Kusters B, et al. Autophagy activity is associated with membranous sodium iodide symporter expression and clinical response to radioiodine therapy in non-medullary thyroid cancer. Autophagy. 2016;12:1195–205.

50 Smith VE, Sharma N, Read ML, Ryan G, Martin A, Boelaert K, et al. Manipulation of PBF/PTTG1IP phosphorylation status; a new therapeutic strategy for improving radioiodine uptake in thyroid and other tumours. J Clin Endocrinol Metab. 2013;98:2876–86.

51 Kawabata T, Yoshimori T. Beyond starvation: An update on the autophagic machinery and its functions. J Mol Cell Cardiol. 2016;95:2–10.

52 Levy JMM, Towers CG, Thorburn A. Targeting autophagy in cancer. Nat Rev Cancer. 2017;17:528–42.

53 Ruschak AM, Slassi M, Kay LE, Schimmer AD. Novel proteasome inhibitors to overcome bortezomib resistance. J Natl Cancer Inst. 2011;103:1007–17.

54 Jivan R, Peres J, Damelin LH, Wadee R, Veale RB, Prince S, et al. Disulfiram with or without metformin inhibits oesophageal squamous cell carcinoma in vivo. Cancer Lett. 2018;417:1–10.

55 Lovborg H, Oberg F, Rickardson L, Gullbo J, Nygren P, Larsson R. Inhibition of proteasome activity, nuclear factor-KappaB translocation and cell survival by the antialcoholism drug disulfiram. Int J Cancer. 2006;118:1577–80.

56 Ye H, Chen M, Cao F, Huang H, Zhan R, Zheng X. Chloroquine, an autophagy inhibitor, potentiates the radiosensitivity of glioma initiating cells by inhibiting autophagy and activating apoptosis. BMC Neurol. 2016;16:178.

57 Cvek B, Dvorak Z. The value of proteasome inhibition in cancer. Can the old drug, disulfiram, have a bright new future as a novel proteasome inhibitor? Drug Discov Today. 2008;13:716–22.

58 Bernardinelli E, Costa R, Nofziger C, Paulmichl M, Dossena S. Effect of Known Inhibitors of Ion Transport on Pendrin (SLC26A4) Activity in a Human Kidney Cell Line. Cell Physiol Biochem. 2016;38:1984–98.

