## Supplementary Information for "New mechanisms of radioiodide uptake revealed via a novel high throughput drug screening approach in thyroid cancer"

**This PDF file includes:**

Supplementary Materials and Methods
Figures: S1 to S20
Tables: S1 to S6
References for SI

#### Supplementary Materials and Methods

##### Validation of the YFP-iodide assay for high throughput screening

The Z’ factor was used to determine the suitability of the YFP-iodide assay for high throughput drug screening. For this, the YFP fluorescence of TPC-1-NIS-YFP cells was quenched with different NaI concentrations (0-68 mM; Fig. 1B) and Z’ factor values calculated using the equation: **Estimated Z’-factor = 1- [3($SD_p$+$SD_n$)/($Mean_p$-$Mean_n$)];** SD = standard deviation; mean = mean YFP fluorescence values; p = positive control (i.e. higher NaI concentration); n = negative control (i.e. lower NaI concentration or PBS control). Z’factor values above 0.5 indicated that the assay was suitable for a full-scale high throughput drug screen.

##### Drug screening and cell viability

In brief, TPC-1 cells expressing NIS and halide-sensitive YFP were seeded into 96-well tissue culture plates at a density of 15,000 cells/well (black-walled, clear-bottom), using plates pre-loaded with test compounds. The Prestwick Chemical Library (~1200 approved compounds) was used for the primary screen at a single dose (10 μM; 1.25% DMSO). All plates contained positive (34 mM NaI) and negative (PBS) controls. The assay was performed using the PHERAstar FS plate reader (BMG Labtech) at 37°C with orbital averaging, 20 flashes/well and focal height kept constant throughout. The initial YFP fluorescence intensity was recorded prior to injection of NaI at a final concentration of 4 mM. YFP fluorescence measurements were taken by the PHERAstar FS plate reader every 3 minutes over 4 cycles and ΔYFP values calculated. In subsequent cell viability assays, cells were incubated with alamarBlue (0.02%; resazurin) solution for 1 hour and absorbance measured using the POLARstar Omega plate reader (BMG Labtech; 572-583 nm optic module).

##### Determination of ΔYFP values

ΔYFP values were calculated to quantify relative changes in YFP fluorescence in drug-treated YFP-cells compared to DMSO-treated controls. In particular, raw YFP fluorescence data was processed using the interquartile mean well-based normalization ($IQM_w$) method (1), according to the following equations:

**(1) %YFP = [($YFP_0$-$YFP_4$) / $YFP_0$] *100**;

%YFP = percentage change in YFP fluorescence; $YFP_0$ = raw YFP fluorescence value at time 0 prior to NaI injection; $YFP_4$= raw YFP fluorescence value after 4 cycles.

**(2)** $S_{IQM} = \%YFP / \mu_{iq}$;

$S_{IQM}$ = sample value after IQM normalization on a per plate basis; $\mu_{iq}$ = TRIMMEAN (values for all wells, 0.5), i.e. interquartile mean of the plate.

**(3)** $S_{IQMW} = S_{IQM} / \mu_{iqw}$;

$S_{IQMW}$ = sample value after IQM normalization for each well position; $\mu_{iqw}$ = TRIMMEAN (values for each well position, 0.5), i.e. mean ($\mu$) of the interquartile values (*iq*) of each well position (*w*) across the entire screen.

**(4)** $\Delta YFP = [S_{IQMW}(DRUG_x) - S_{IQMW}(DMSO_x)] / SD_{SIQMW\,(DMSOn)}$

ΔYFP = fold-change in YFP fluorescence for drug treatment wells relative to SD of YFP fluorescence in DMSO control wells. Candidate drugs were considered to be top hits in the primary screen at ΔYFP values > 1.5-2; $S_{IQMW}$ ($DRUG_x$) = sample value after IQM normalization for each drug treatment well; $S_{IQMW}$ ($DMSO_x$) = sample value after IQM normalization for plate-matched DMSO control wells; $SD_{SIQMW\,(DMSOn)}$ = SD of sample values after IQM normalization for DMSO control wells across the entire screen ($n$ = 120). Positive ΔYFP values denote increased intracellular iodide, whereas negative ΔYFP values indicate decreased intracellular iodide.

#### Drug ranking and statistics

ΔYFP, area under the curve (AUC), EC50 and RAI (iodine-125) uptake values were used to rank the potency of drugs to increase intracellular iodide levels. AUC values were determined ($log_{10}$ scale; GraphPad Prism) for positive peaks only (Fig. 1G; Supplementary Fig. S4C). AUC values for drug-treated TPC-1-YFP cells were subtracted from corresponding values in TPC-1-NIS-YFP cells to determine ΔAUC values (Supplementary Fig. S4B). EC50 values were calculated using the Quest Graph EC50 calculator [AAT Bioquest, Inc. (2020, April 08). Quest Graph™ EC50 Calculator. https://www.aatbio.com/tools/ec50-calculator].

All results were obtained from triplicate experiments unless otherwise indicated. For comparison between two groups, data were subjected to the Student’s t-test, and for multiple comparisons one-way ANOVA with either Dunnett’s or Tukey’s post-hoc test was used. $P<0.05$ was considered significant.

#### TCGA data analysis and survival

Normalized gene expression data and clinical information for papillary thyroid cancer (PTC) were downloaded from TCGA via cBioPortal (cbioportal.org/), FireBrowse (firebrowse.org) and NCI Genomic Data Commons (GDC; portal.gdc.cancer.gov/) (4-7). Non-parametric Kolmogorov-Smirnov, Kruskal-Wallis, and Spearman's correlation tests were performed as TCGA expression levels were not normally distributed. *P*-values were adjusted using the Benjamini-Hochberg FDR correction procedure (www.biostathandbook.com) to correct for multiple comparisons. Dunn's multiple comparison post-hoc testing was used after Kruskal-Wallis tests to determine significance between particular datasets in groups of 3 or more. Fisher's exact test was used to determine the significance of nonrandom associations between two categorical variables. For disease-free survival (DFS), the median value of gene expression (log2) in TCGA was used to stratify patients with high or low expression. Subsequent analyses were completed using Kaplan-Meier curves and cohort comparisons performed by the Log rank (P) test.

#### Connectivity Map (CMAP) and functional classification of candidate genes

The Connectivity Map (CMAP) resource (https://clue.io/cmap) was used to probe the transcriptional relationship between top drug hits (47 available in CMAP) and loss of function of a panel of 45 proteostasis genes (2, 3). Hierarchical cluster analysis was used to identify subsets of genes (e.g. VCP) associated with particular drugs.

Candidate proteostasis genes (Supplementary Fig. S16) to be studied for clinical relevance were pooled from three sources: (i) known NIS interactors from a recent mass spectrometry study (8), (ii) a well-characterized 142 proteostasis geneset, and (iii) genes associated with VCP overexpression in RAI-treated BRAF patients in TCGA (Fig. 6D). Proteostasis genes were filtered from this gene pool based on DAVID functional classification (9, 10) and/or entry in the human autophagy database (HADb; autophagy.lu/).

.

Figure S1

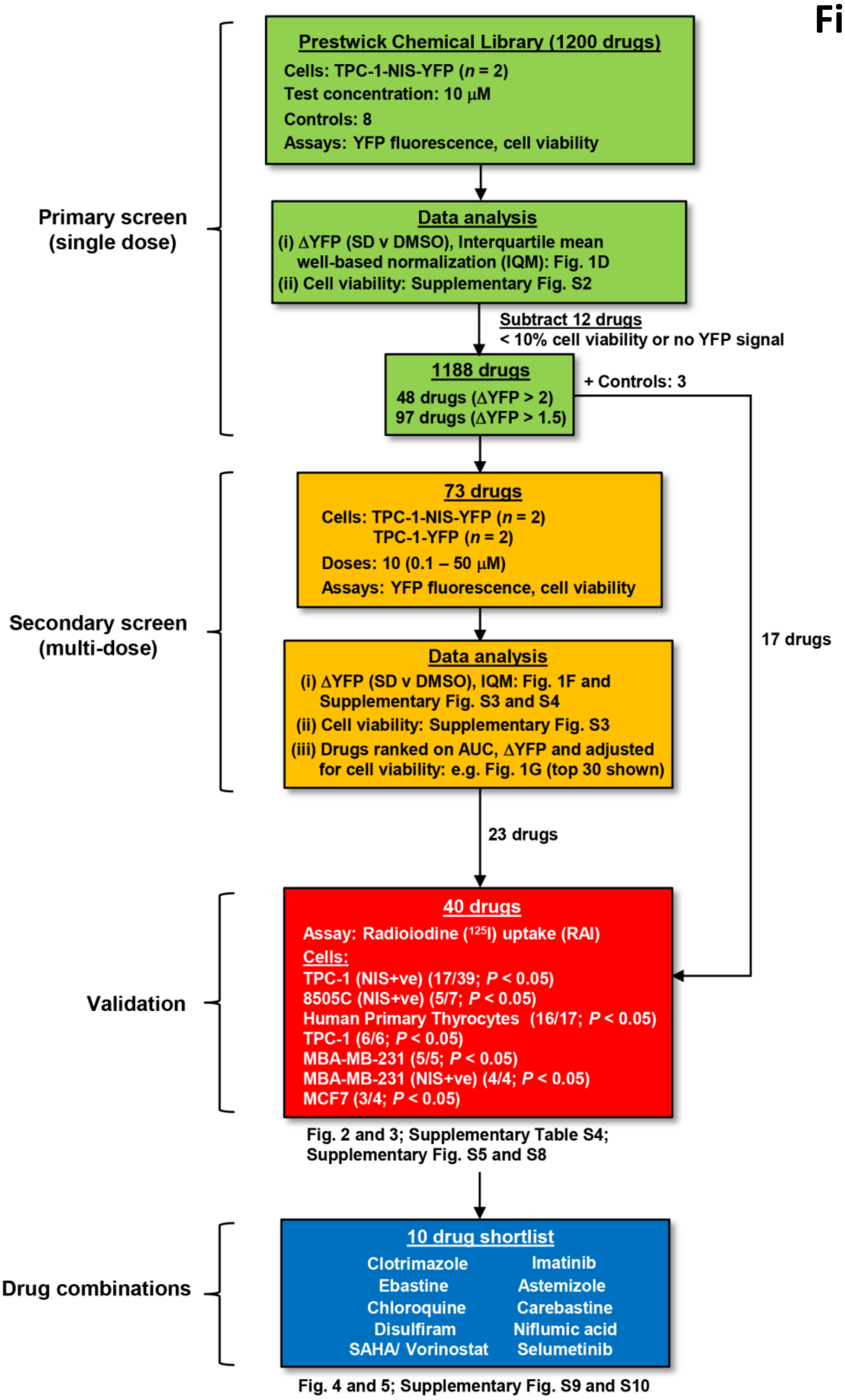


**Fig. S1. Identification of multiple novel compounds that enhance radioiodide (RAI) uptake.** Schematic showing details of the 4 different stages used to identify multiple novel drugs that enhance cellular uptake of RAI. A high throughput approach was initially adopted in the primary screen (green-colored boxes) using the Prestwick Chemical Library (1200 drugs, single dose) and a modified YFP biosensor for intracellular iodide. Ten different doses for 73 drugs were further examined using the YFP biosensor in the secondary screen (orange-colored boxes) which also compared two YFP-expressing cell lines (± NIS). From this, 40 drugs at multiple doses were validated in RAI-uptake assays (red-colored box) in a range of different cell types (e.g. human primary thyrocytes). Finally, a shortlist of 10 drugs was generated (blue-colored box) and different drug combinations evaluated in RAI uptake assays in multiple cell types.

#### Figure S2

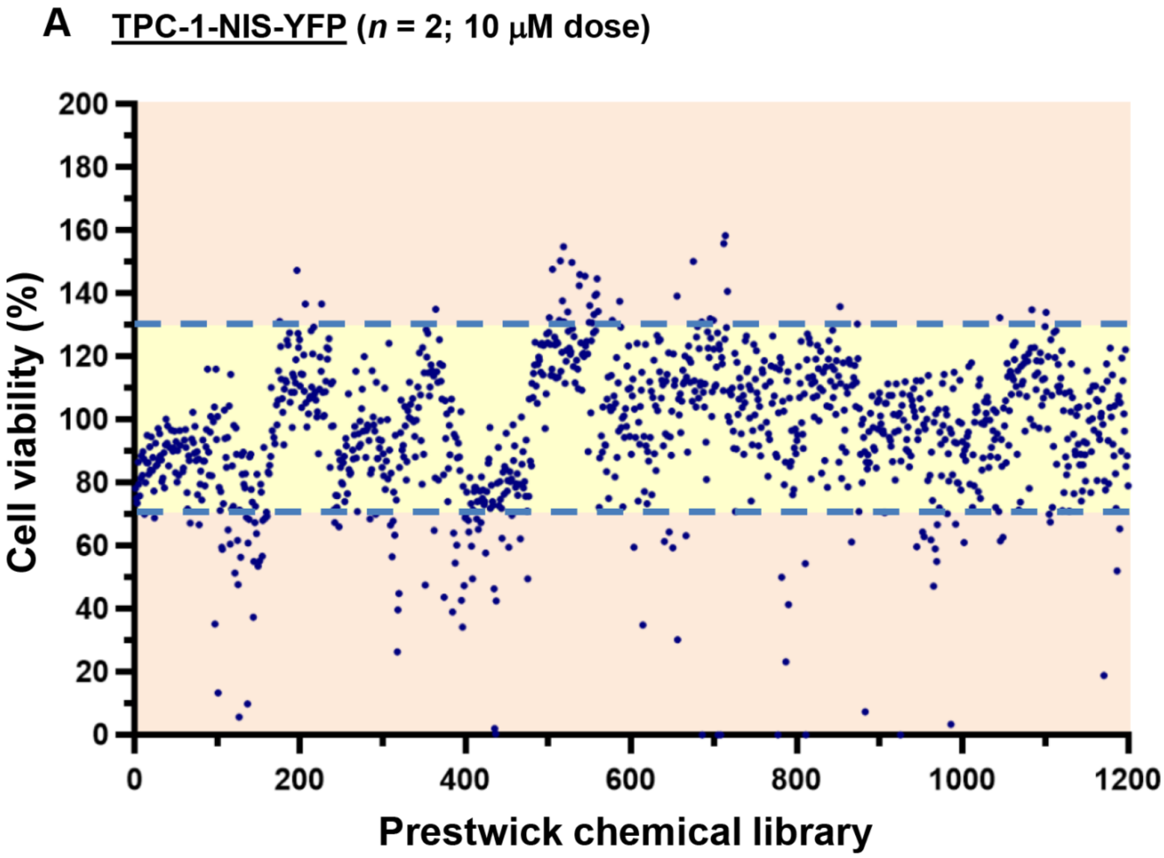


**B**

| Cell Viability | Number of drugs | % of drugs |
|---|---|---|
| ≥ 70% | 1095 | 91.3 |
| ≥ 80% | 971 | 80.9 |
| ≥ 90% | 777 | 64.8 |

**Fig. S2. Assessment of the Prestwick Chemical Library on TPC-1-NIS-YFP cell viability.** (*A*) TPC-1-NIS-YFP cell viability following treatment with a 10 µM dose of 1200 drugs in the Prestwick Chemical Library for 24 hours. Dashed blue line indicates cell viability at either 70 or 130%. Each circle represents a mean value from 2 independent experiments. (*B*) Table showing number of Prestwick drugs out of 1200 associated with TPC-1-NIS-YFP cellular viability equal or greater than 70, 80 or 90%. Cellular viability determined using the alamarBlue (resazurin) assay.

### Figure S3

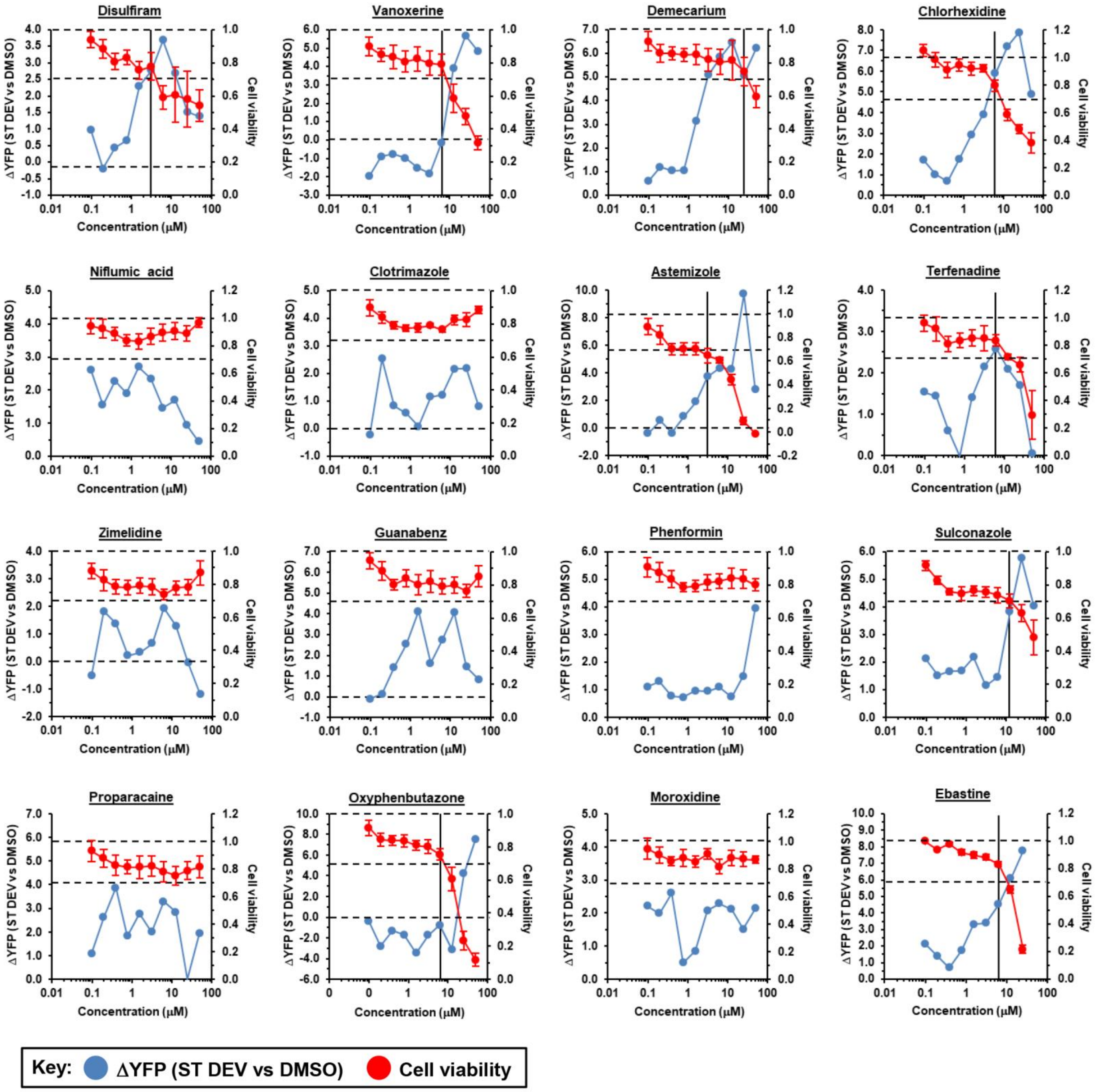


**Fig. S3. Dose-response curves and cell viability in drug-treated TPC-1-NIS-YFP cells.**
YFP-based biosensing of intracellular iodide of TPC-1-NIS-YFP cells treated with Prestwick Chemical library drugs as indicated for 24 hours at 10 different doses (0.1 – 50 μM). ΔYFP values signify changes in intracellular iodide (i.e. positive ΔYFP values denote increased intracellular iodide) and were derived relative to standard deviation (ST DEV) of YFP fluorescence of DMSO-treated TPC-1-NIS-YFP cells (*n*=120). An interquartile mean well-based method ($IQM_w$) was used to normalise YFP fluorescence measurements prior to calculation of ΔYFP values. 16 representative drug profiles are shown out of 73 performed in the secondary screen (multi-dose, Fig. S1). Blue circles represent mean ΔYFP values from 2 independent experiments. Cell viability (red circles) was determined using the alamarBlue (resazurin) assay (mean values ±SE, *n*=4). Significant toxicity was apparent for certain drugs at higher doses (e.g. vanoxerine, chlorhexidine). In these cases, specific parameters (e.g. AUC; Fig. 1G; Fig. S4B and C) were calculated only up to maximum drug doses associated with cell viability >70% (indicated by vertical black line).

**Figure S4**

A

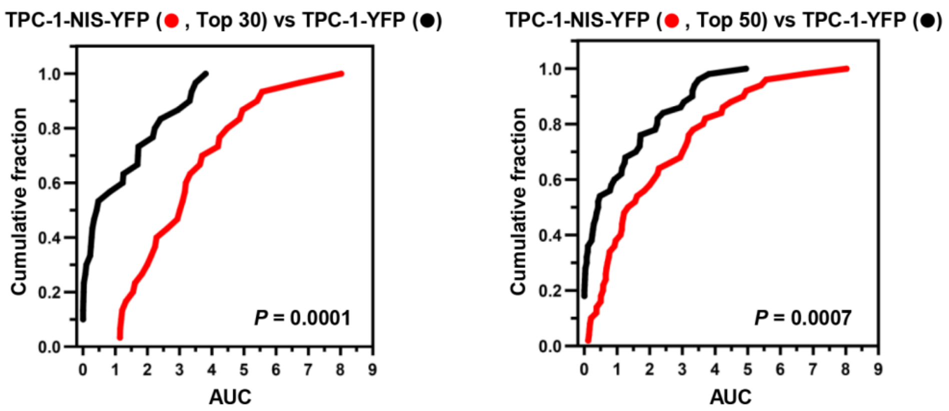


**B** TPC-1-NIS-YFP vs TPC-1-YFP cells (drug doses > 70% cell viability; i.e adjusted for YFP effects)

| | Drug | ΔAUC | | Drug | ΔAUC | | Drug | ΔAUC |
|---|---|---|---|---|---|---|---|---|
| 1 | Proparacaine | 5.31 | 11 | Sulconazole | 3.09 | 21 | Hesperidin | 1.14 |
| 2 | Moroxidine | 4.57 | 12 | Azathioprine | 2.86 | 22 | Levocabastine | 1.13 |
| 3 | Demecarium | 4.53 | 13 | Florfenicol | 2.61 | 23 | Halofantrine | 1.05 |
| 4 | Nortriptyline | 4.24 | 14 | Terfenadine | 2.53 | 24 | Articaine | 0.95 |
| 5 | Niflumic acid | 3.70 | 15 | Chlorhexidine | 2.07 | 25 | Khellin | 0.92 |
| 6 | Alverine | 3.41 | 16 | Hexachlorophene | 2.00 | 26 | Retinoic acid | 0.79 |
| 7 | Pyrvinium | 3.36 | 17 | Ethacrynic acid | 1.98 | 27 | Quinidine | 0.68 |
| 8 | Clotrimazole | 3.32 | 18 | Zimelidine | 1.81 | 28 | Molindone | 0.66 |
| 9 | (R)-(+)Atenolol | 3.16 | 19 | Amrinone | 1.77 | 29 | Astemizole | 0.60 |
| 10 | Guanabenz | 3.16 | 20 | Ebastine | 1.25 | 30 | Amikacin | 0.54 |

**C** TPC-1-YFP cells only (drug doses > 70% cell viability)

| | Drug | AUC | ΔYFP | Dose (μM) | | Drug | AUC | ΔYFP | Dose (μM) |
|---|---|---|---|---|---|---|---|---|---|
| 1 | Prednisolone | 4.95 | 3.27 | 0.78 | 15 | Guanabenz | 2.40 | 1.94 | 3.13 |
| 2 | Piperacetazine | 3.81 | 3.02 | 12.5 | 16 | Fenofibrate | 2.25 | 1.59 | 12.5 |
| 3 | Demecarium | 3.50 | 4.51 | 12.5 | 17 | Praziquantel | 2.22 | 1.91 | 0.39 |
| 4 | Pyrvinium | 3.37 | 4.99 | 0.78 | 18 | Piretanide | 1.41 | 1.86 | 0.1 |
| 5 | Phenformin | 3.31 | 3.90 | 50 | 19 | Niflumic acid | 1.26 | 1.87 | 0.39 |
| 6 | Mianserine | 3.30 | 4.34 | 50 | 20 | Ticonazole | 1.15 | 1.54 | 0.20 |
| 7 | Sertaconazole | 3.05 | 3.57 | 25 | 21 | Tridihexethyl | 0.92 | 1.92 | 50 |
| 8 | Ebastine | 2.94 | 4.56 | 6.25 | 22 | Retinoic acid | 0.83 | 1.89 | 0.39 |
| 9 | Dexamethasone | 2.24 | 2.03 | 0.78 | 23 | Clomiphene | 0.82 | 1.81 | 12.5 |
| 10 | Chlorhexidine | 2.17 | 2.49 | 6.25 | 24 | Clomipramine | 0.78 | 1.67 | 25 |
| 11 | Ethacrynic acid | 1.72 | 2.02 | 0.10 | 25 | Vanoxerine | 0.73 | 1.64 | 6.25 |
| 12 | Disulfiram | 1.69 | 2.32 | 0.39 | 26 | Bisacodyl | 0.42 | 1.86 | 50 |
| 13 | Fendiline | 1.58 | 2.41 | 12.5 | | | | | |
| 14 | Methotrimeprazine | 1.38 | 2.82 | 50 | | | | | |

**Fig. S4. Ranking of Prestwick Chemical Library drugs that increase intracellular iodide.** (*A*) Cumulative frequency distribution plot comparing the area under the curve (AUC) calculations for either the top 30 (left) or 50 (right) Prestwick Chemical Library drugs identified to increase intracellular iodide in TPC-1-NIS-YFP versus TPC-1-YFP cells; *P*-values determined by the Kolmogorov-Smirnov test. (*B*) Ranking of Prestwick Chemical Library drugs that increase intracellular iodide in thyroid cancer cells based on ΔAUC (i.e. difference in AUC values) as shown between TPC-1-NIS-YFP and TPC-1-YFP cells treated with 10 different doses (0.1 – 50 μM). AUC values calculated up to maximum drug doses associated with >70% cell viability. (*C*) Ranking of Prestwick Chemical Library drugs that increase intracellular iodide in drug-treated TPC-1-YFP cells based on AUC and ΔYFP values (pale yellow > 2; green > 1.5). AUC values calculated up to maximum drug doses associated with >70% cell viability. Maximal ΔYFP responses and associated drug doses (μM) are shown.

### Figure S5

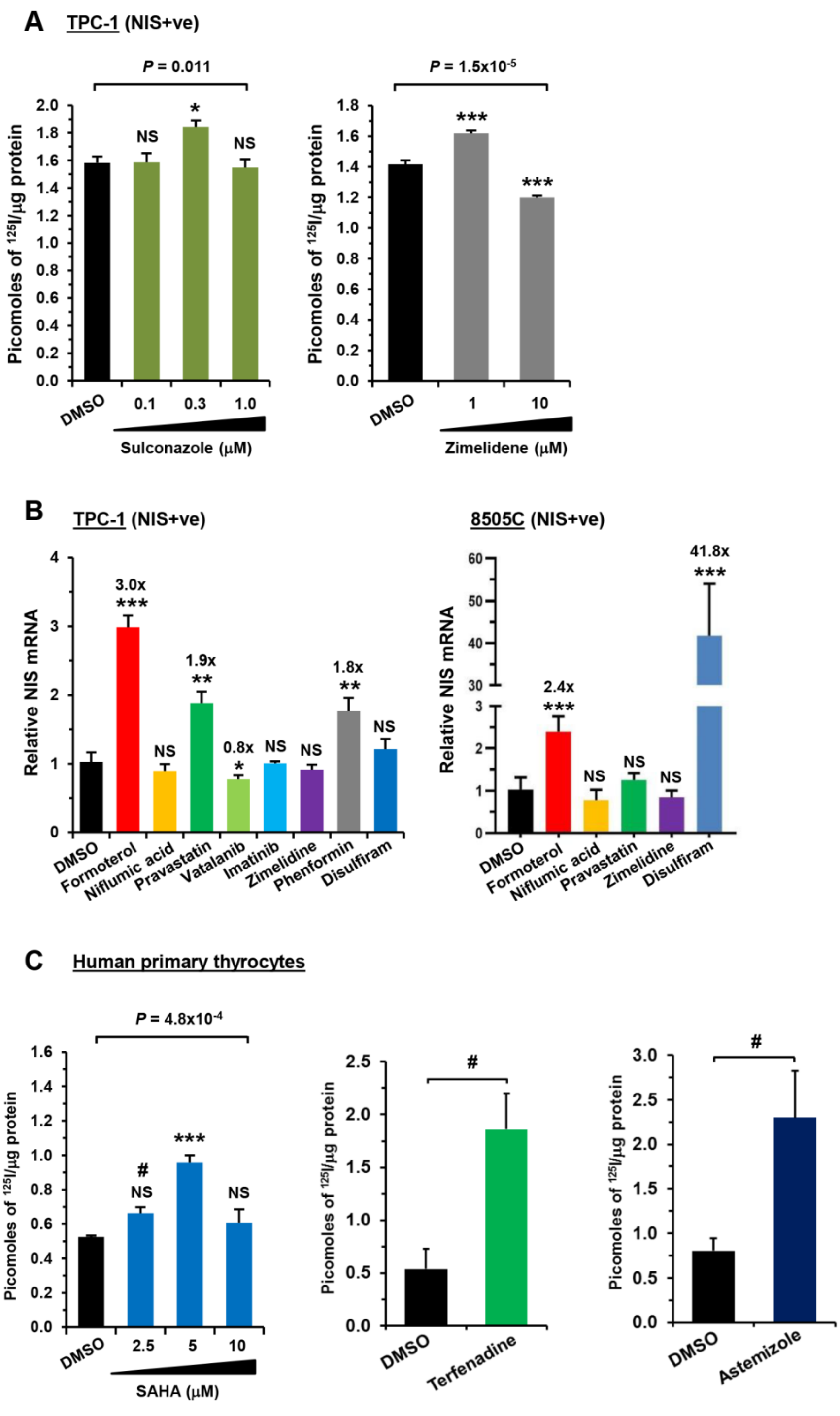


**Fig. S5. Identification of drugs enhancing NIS mRNA expression and RAI uptake.**

(*A*) RAI uptake of TPC-1 (NIS+ve) cells treated with sulconazole or zimelidene at indicated doses for 24 hours. NS, not significant; *, $P$<0.05; ***, $P$<0.001; one-way ANOVA followed by Dunnett's post hoc test. (*B*) Relative fold-change in NIS mRNA expression in TPC-1 (NIS+ve) (left) or 8505C (NIS+ve) cells (right) treated with indicated drug for 24 hours. Doses: formoterol (10 μM), niflumic acid (15 μM), pravastatin (40 μM), vatalanib (1.0 μM), imatinib (1.0 μM), zimelidine (1.0 μM), phenformin (250 μM) and disulfiram (0.5 μM). NS, not significant; *, $P$<0.05; **, $P$<0.01; ***, $P$<0.001. (*C*) RAI uptake of primary human thyrocytes treated with SAHA at indicated doses (left, $n$=3), 1.0 μM terfenadine (middle, $n$=3) or 0.25 μM astemizole (right, $n$=11) for 24 hours; one-way ANOVA followed by Dunnett's post hoc test or unpaired t-test. NS, not significant; one-way ANOVA: ***, $P$<0.001; Unpaired t-tests: #, $P$<0.05.

#### Figure S6

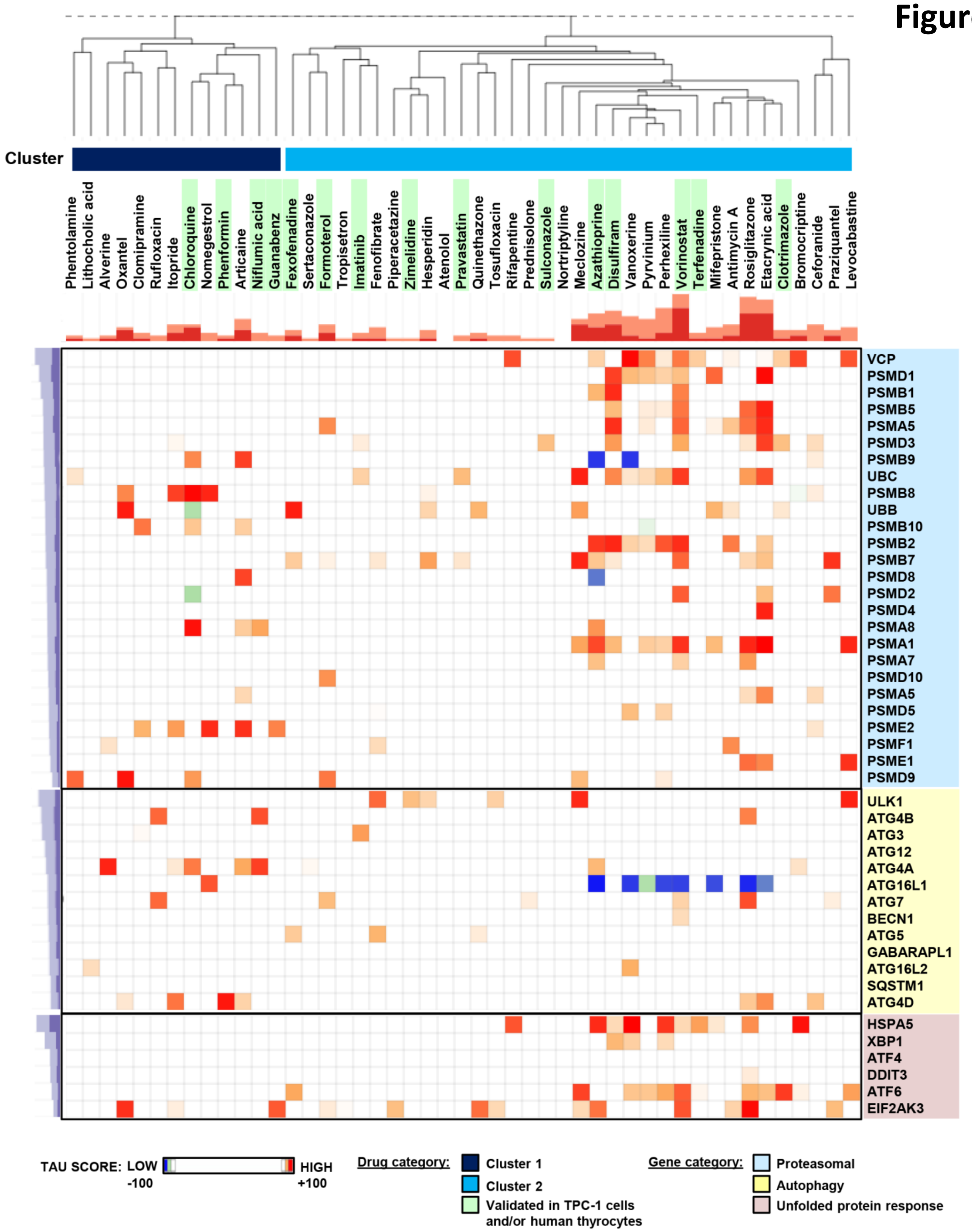


**Fig. S6. Connectivity Map (CMAP) analysis reveals cluster associated with proteasomal genes.** CMAP (L1000 assay) hierarchical cluster analysis comparing the transcriptional signatures of 47 drug treatments with those of 45 genetic perturbations involving loss of function of proteostasis genes (i.e. 26 proteasomal, 13 autophagy and 6 unfolded-response protein genes). Two main clusters were apparent from the analysis (cluster 1, dark blue; cluster 2, light blue). 15 of these drug treatments (highlighted in green) have been validated to enhance radioiodide uptake in thyroid cancer cells (Figs. 2-5; Fig. S5; Table S2). The transcriptional signatures of several drug treatments in our top 50 candidate drugs were unavailable in the CMAP dataset to include in the analysis (e.g. astemizole, carebastine, ebastine, fexofenadine, vatalanib). A tau score of +90 or higher, and of -90 or lower, are considered to be strong scores.

| Drug | 2D Structure | 3D Structure | Evidence in VCP inhibitor studies |
|---|---|---|---|
| **Ebastine** | 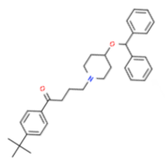 | 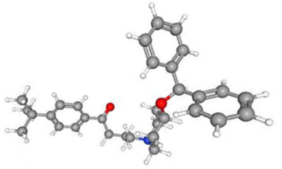 | <u>(1) Segura-Cabrera A *et al.*, (2017); Ref (11)</u><br>Structure-based virtual screening (-12.57 kcal/mol)<br>Cell-based/ biochemical assays<br><u>(2) Fletcher A *et al.*, (2020); Ref (8)</u><br>Radioiodine (RAI) uptake/ VCP siRNA/ cell surface biotinylation (CSB) assays<br><u>(3) This study</u>: Drug combination/ RAI uptake assays |
| **Astemizole** | 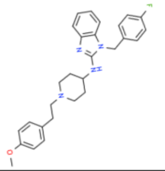 | 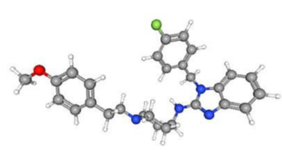 | <u>(1) Segura-Cabrera A *et al.*, (2017); Ref (11)</u><br>CMAP (build 02): MCF7 (Correlation *P*-value = 8.4E-39)<br>Structure-based virtual screening (-11.57 kcal/mol)<br>Cell-based/ biochemical assays<br><u>(2) Fletcher A *et al.*, (2020); Ref (8)</u><br>RAI uptake/ VCP siRNA/ CSB assays<br><u>(3) This study</u>: Drug combination/ RAI uptake assays |
| **Terfenadine** | 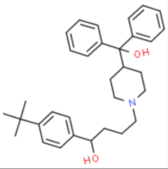 | 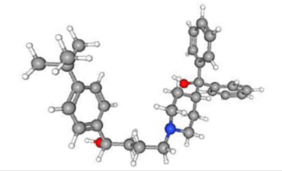 | <u>(1) Segura-Cabrera A *et al.*, (2017); Ref (11)</u><br>CMAP (build 02): MCF7 (Correlation *P*-value = 3.0E-31)<br><u>(2) This study</u><br>CMAP (L1000)/ RAI uptake assays |
| **Clotrimazole** | 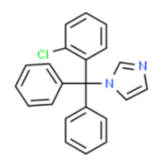 | 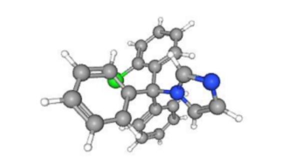 | <u>(1) Segura-Cabrera A *et al.*, (2017); Ref (11)</u><br>CMAP (build 02): MCF7 (Correlation *P*-value = 4.0E-46)<br>Structure-based virtual screening (-8.19 kcal/mol)<br>Cell-based/ biochemical assays<br><u>(2) Fletcher A *et al.*, (2020); Ref (8)</u><br>RAI uptake/ VCP siRNA/ CSB assays<br><u>(3) This study</u>:<br>CMAP (L1000)/ Drug combination/ RAI uptake assays |
| **Carebastine**<br>(active carboxylic acid metabolite of Ebastine) | 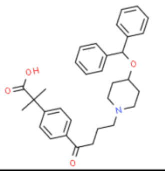 | 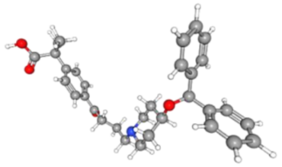 | <u>(1) This study</u>:<br>Drug combination/ RAI uptake/ VCP siRNA assays |
| **Fexofenadine**<br>(active metabolite of Terfenadine) | 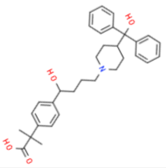 | 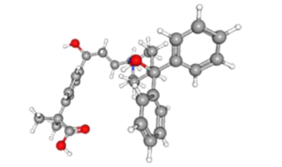 | <u>(1) This study</u>:<br>RAI uptake assays |
| **Disulfiram** | 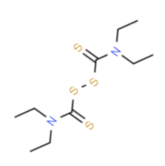 | 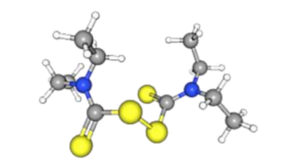 | <u>(1) Segura-Cabrera A *et al.*, (2017); Ref (11)</u><br>CMAP (build 02): MCF7 (Correlation *P*-value = 4.5E-46)<br>Structure-based virtual screening (-6.15 kcal/mol)<br><u>(2) Skrott Z *et al.*, (2017); Ref (12)</u><br>Disulfiram metabolite (DTC) binds NPL4 to disable the VCP-NPL4-UFD1 pathway<br><u>(3) This study</u>: Drug combination/ RAI uptake assays |
| **NMS-873** | 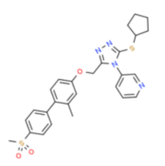 | 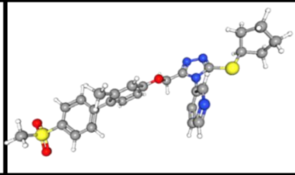 | <u>(1) Well-established VCP inhibitor</u><br>Used as a control drug for VCP inhibitor studies<br><u>(2) This study</u>:<br>Drug combination/ RAI uptake assays |

**Fig. S7. Association of validated drugs with VCP loss of function.**
Details are shown for 8 drugs identified in the study that enhance RAI uptake and are associated with VCP loss of function. A 2D and 3D image of each drug are provided, as well as brief notes on recent evidence implicating their role in abrogating VCP activity. NMS-873 is a well-established VCP inhibitor and has been included for comparison. CMAP (L1000) is the current dataset and relies on perturbational data generated using the L1000 assay. The previous Affymetrix-based CMAP dataset is referred to as (build 02). Source: ChemSpider; PubChem.

#### Figure S8

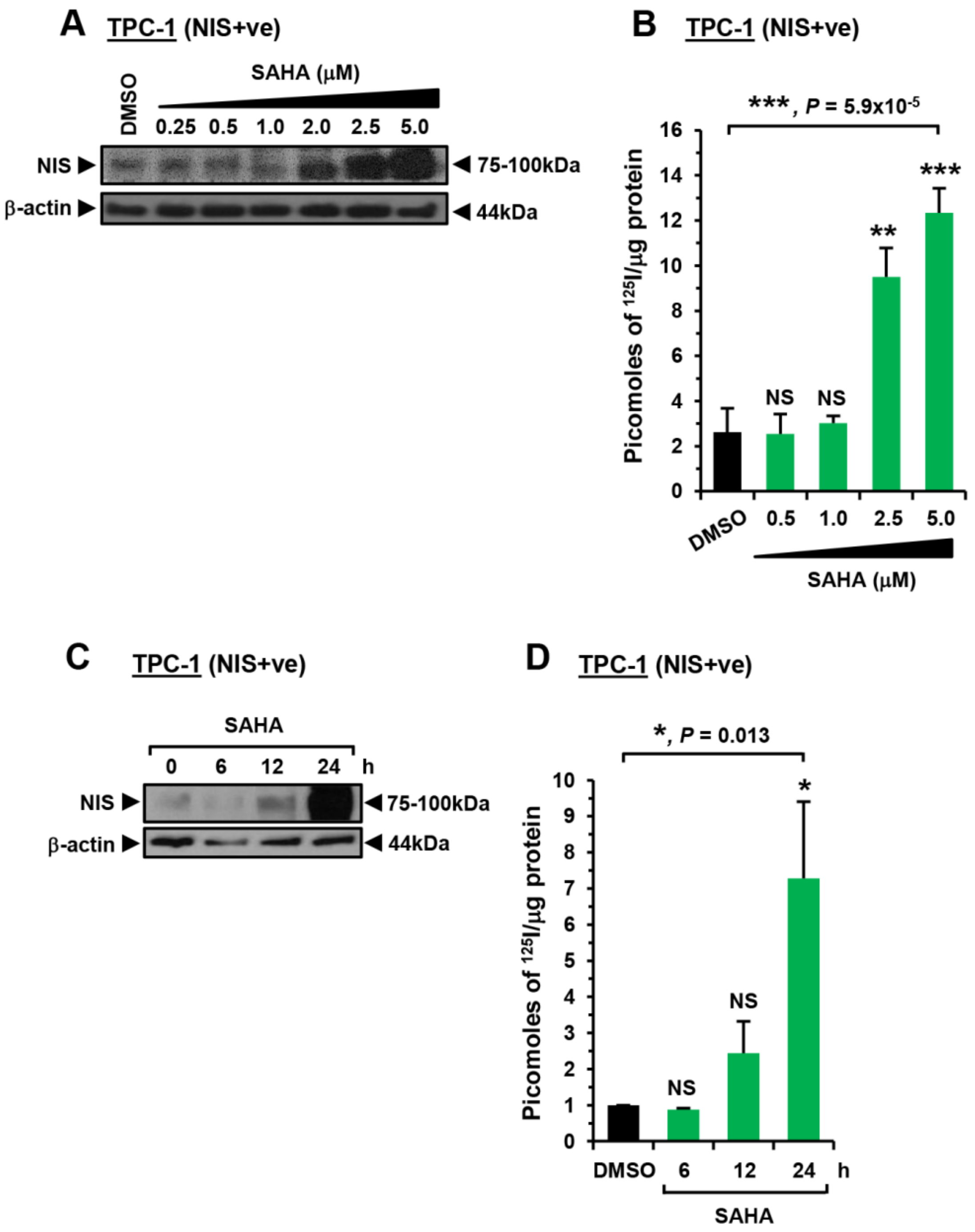


**Fig. S8. SAHA induces NIS protein and RAI uptake in a dose-dependent manner.**
(*A*) Western blot analysis of NIS expression in TPC-1 (NIS+ve) cells treated with SAHA at indicated doses for 24 hours. (*B*) RAI uptake of TPC-1 (NIS+ve) cells treated with SAHA at indicated doses for 24 hours. NS, not significant; **, $P<0.01$; ***, $P<0.001$; one-way ANOVA followed by Dunnett's post hoc test. (*C*) Time course of relative NIS protein levels in TPC-1 (+ve) cells treated with 2.5 µM SAHA. (*D*) Time course of RAI uptake in TPC-1 (NIS+ve) cells treated with 2.5 µM SAHA. NS, not significant; *, $P<0.05$; one-way ANOVA followed by Dunnett's post hoc test.

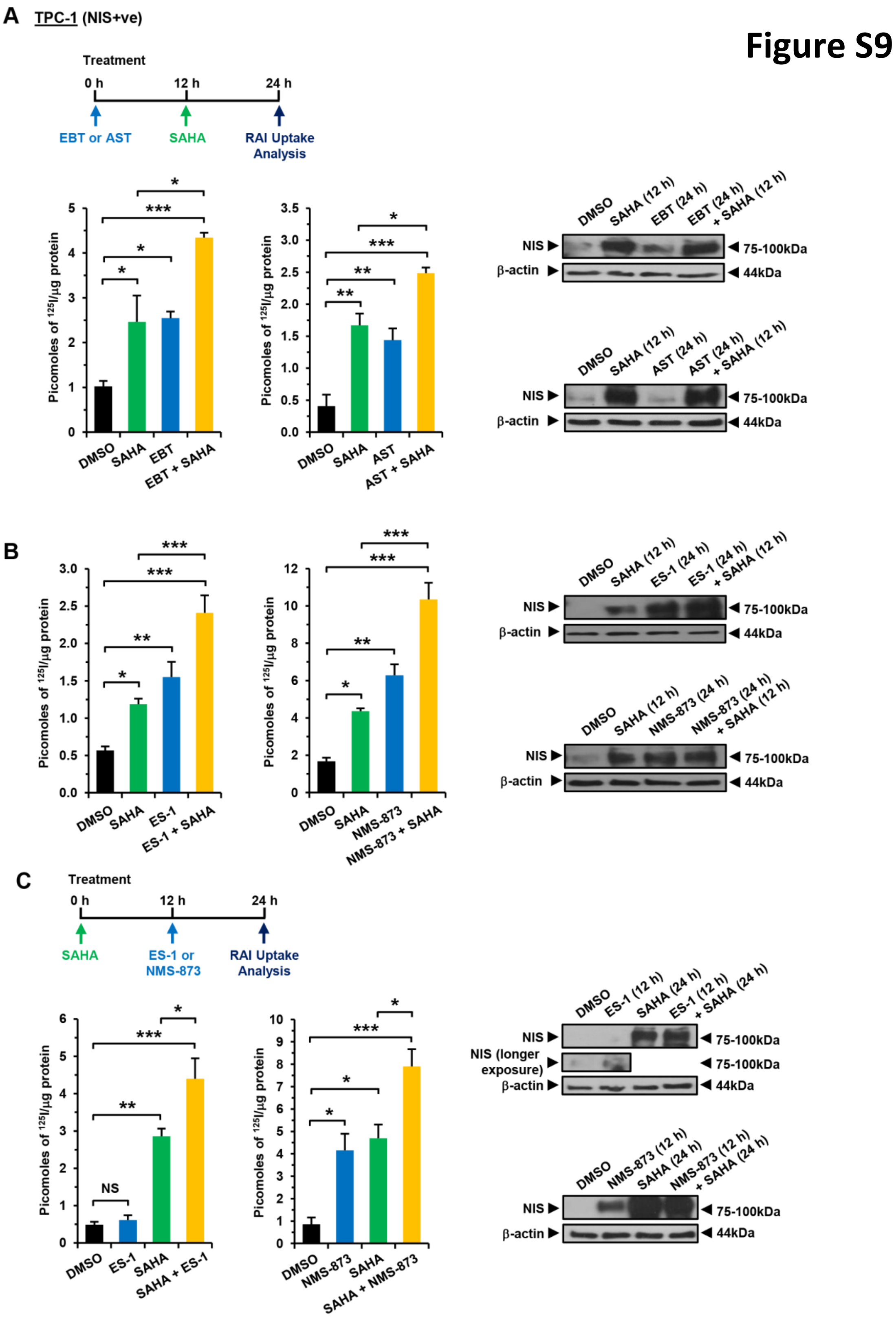


**Fig. S9. Increased RAI uptake by combining VCP inhibitors (VCPi) with SAHA.**
(*A*) RAI uptake (left) and Western blot analyses (right) in TPC-1 (NIS+ve) cells treated with VCP inhibitors, i.e. 0.5 μM ebastine (EBT) and 0.25 μM astemizole (AST), either alone or in combination with 2.5 μM SAHA following the indicated time-course (schematic at top). (*B*) Same as (*A*) but cells treated with 2.5 μM eeyarestatin-1 (ES-1) or 5 μM NMS-873. (*C*) Same as (*B*) but cells treated with ES-1 or NMS-873 following a modified time-course as shown in schematic (i.e. SAHA added 12 hours prior to ES-1 or NMS-873). NS, not significant; *, $P<0.05$; **, $P<0.01$; ***, $P<0.001$; one-way ANOVA followed by Tukey's post hoc test.

**Figure S10**

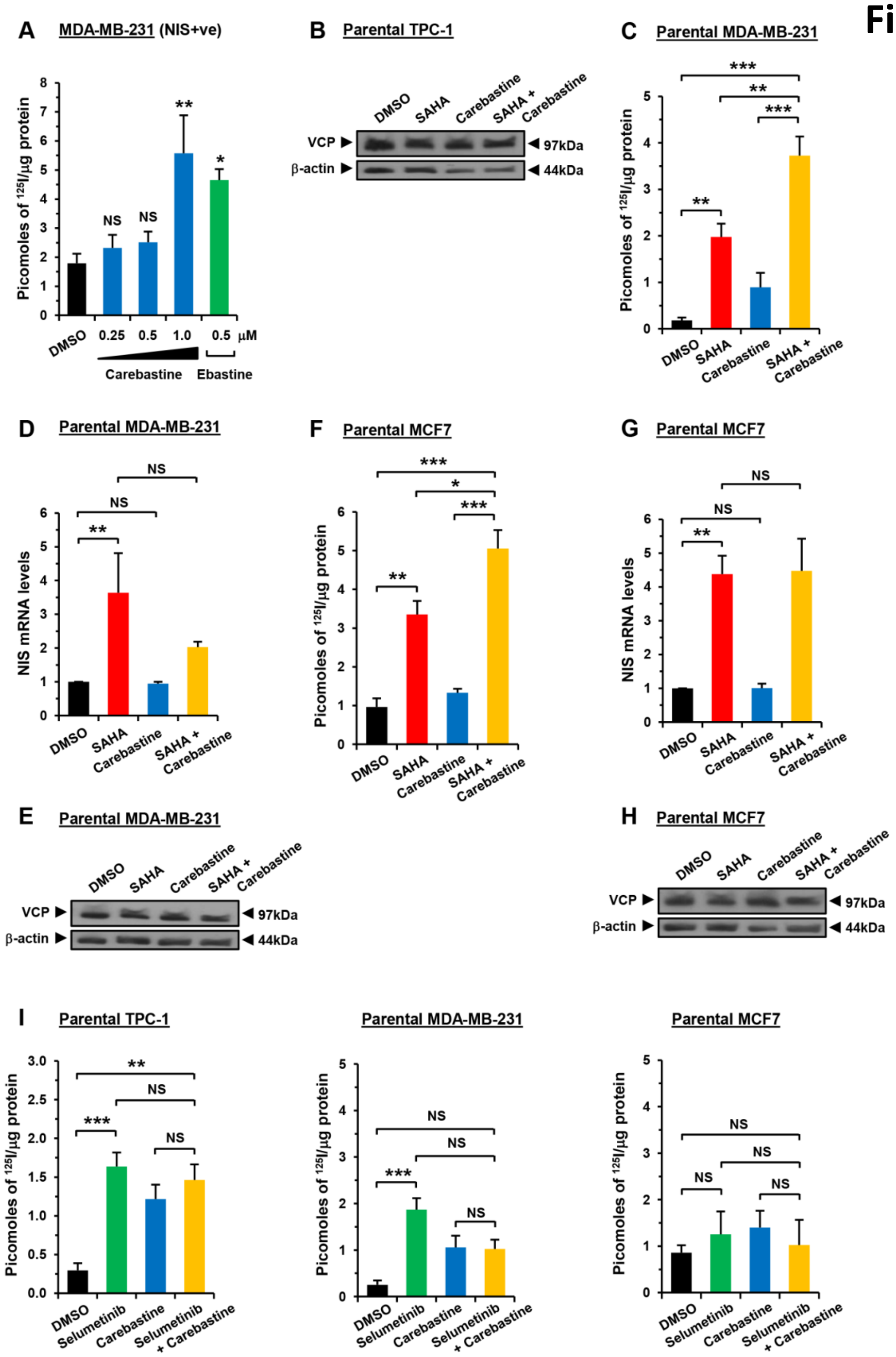


**Fig. S10. Increased RAI uptake with carebastine and SAHA in breast cancer cells.** (*A*) RAI uptake of MDA-MB-231 (NIS+ve) cells treated with carebastine at indicated doses for 24 hours. Controls: DMSO and 0.5 µM ebastine. NS, not significant; *, $P$<0.05; **, $P$<0.01; one-way ANOVA followed by Dunnett's post hoc test. (*B*) Western blot analysis of VCP expression in TPC-1 (NIS+ve) cells treated with 2.5 µM SAHA and 0.5 µM ebastine either alone or in combination. (*C*) RAI uptake in parental MDA-MB-231 cells treated with 2.5 µM SAHA and 1 µM carebastine either alone or in combination. **, $P$<0.01; ***, $P$<0.001; one-way ANOVA followed by Tukey's post hoc test. (*D*) Same as (*C*) but relative fold-change in NIS mRNA expression determined in drug-treated parental MDA-MB-231 cells. NS, not significant; **, $P$<0.01. (*E*) Same as (*B*) but with parental MDA-MB-231 cells. (*F-H*) Same as (*C-E*) but with parental MCF7 cells. (*I*) RAI uptake of parental TPC-1 (left), MDA-MB-231 (middle) or MCF7 cells (right) treated with 0.2 µM selumetinib and 1 µM carebastine either alone or in combination. **, $P$<0.01; ***, $P$<0.001; one-way ANOVA followed by Tukey's post hoc test.

### Figure S11

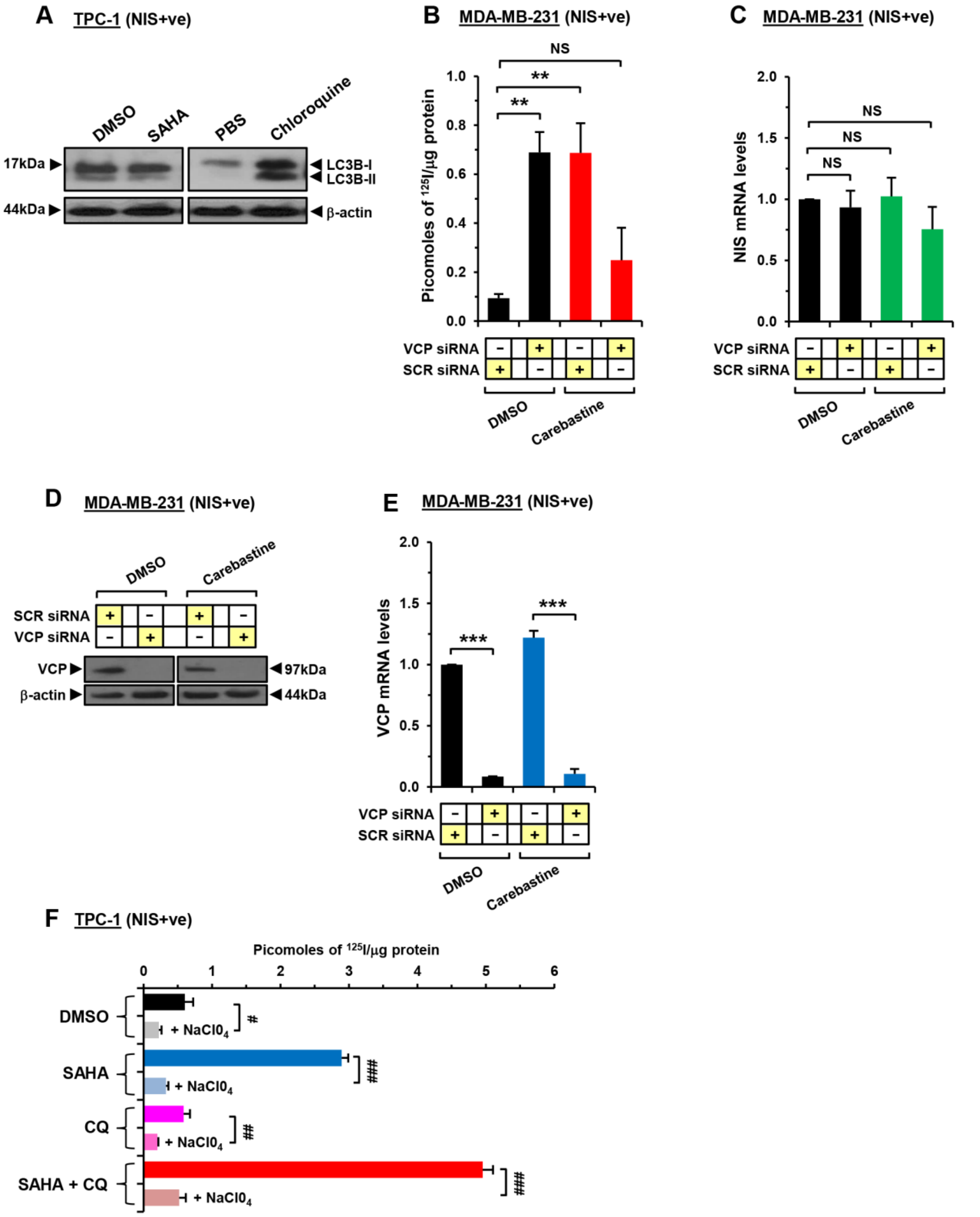


**Fig. S11. Carebastine does not induce RAI uptake in VCP-ablated MDA-MB-231 (NIS+ve) cells.** (*A*) Western blot analysis of LC3B-I and LC3B-II expression in TPC-1 (NIS+ve) cells treated with 5 μM SAHA (left) or 50 μM chloroquine (right). (*B*) RAI uptake of MDA-MB-231 (NIS+ve) cells following VCP-siRNA depletion and treatment with 1 μM carebastine. NS, not significant; **, $P<0.01$; one-way ANOVA followed by Dunnett's post hoc test. (*C*) Same as (*B*) but relative fold-change in NIS mRNA expression determined in MDA-MB-231 (NIS+ve) cells. NS, not significant. (*D*) Western blot analysis of VCP expression in MDA-MB-231 (NIS+ve) cells following VCP-siRNA depletion and treatment with 1 μM carebastine. (*E*) Same as (*C*) but with relative fold change in VCP mRNA expression. ***, $P<0.001$. (*F*) RAI uptake in TPC-1 (NIS+ve) cells treated with chloroquine (CQ) alone or in combination with SAHA. Cells treated with sodium perchlorate (+ $NaClO_4$) as indicated prior to RAI uptake assay; unpaired t-tests: $^{\#}$, $P<0.05$; $^{\#\#}$, $P<0.01$; $^{\#\#\#}$, $P<0.001$.

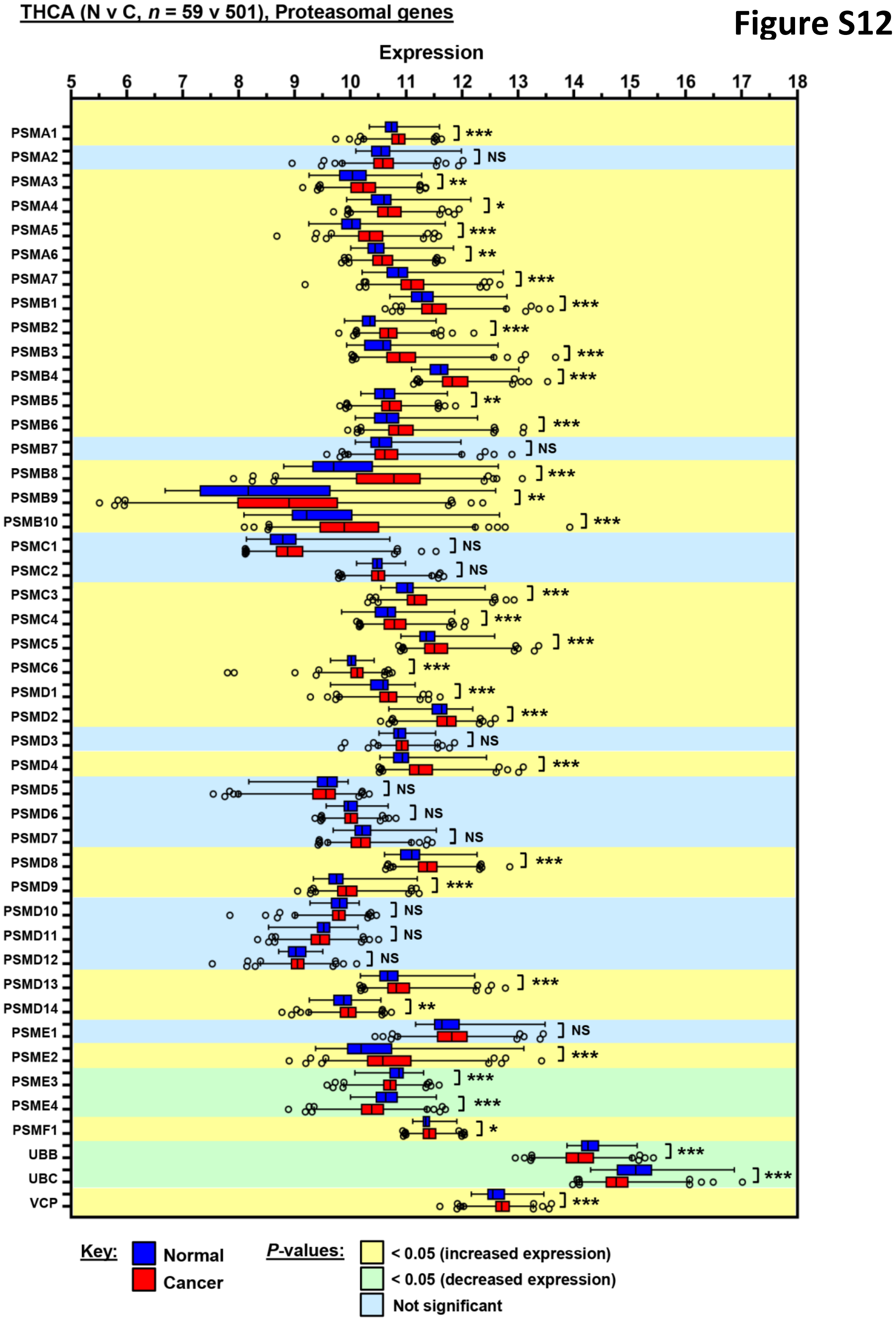


**Fig. S12. Elevated expression of proteasomal genes in PTC.**
Expression ($log_2$) of 45 proteasomal genes in normal thyroid (*n*=59, dark blue) versus PTC (*n*=501, red) in the THCA TCGA dataset. Relative gene expression in PTC versus normal thyroid is highlighted, i.e. increased (yellow), reduced (green) or not significantly different (light blue). NS, not significant; *, *P*<0.05; **, *P*<0.01; ***, *P*<0.001; *P*-values adjusted using the Benjamini-Hochberg FDR correction procedure.

**Figure S13**

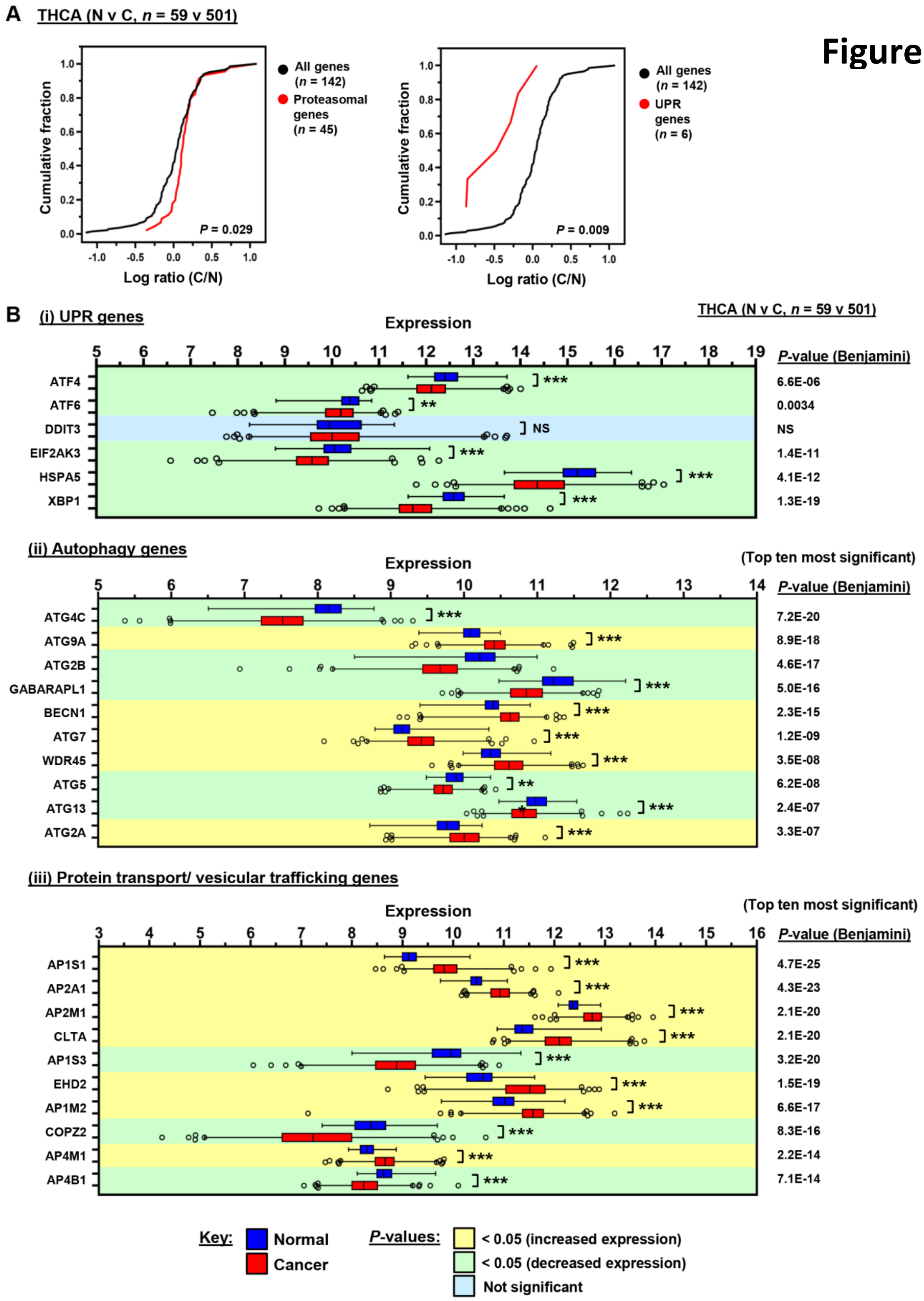


**Fig. S13. Altered expression of proteostasis genes in PTC.**
(*A*) Cumulative frequency distribution plot comparing the $\log_2$ ratio of expression (C/N) for 142 proteostasis-related genes versus 45 proteasomal (left) or 6 UPR (right) genes in THCA TCGA (N v C, *n*=59 v 501); *P*-values determined by Kruskal-Wallis test followed by Dunn's post hoc test. (*B*) Expression ($\log_2$) of (i) 6 UPR, (ii) 10 autophagy or (iii) 10 protein transport/ vesicular trafficking (PT/VT) genes in normal thyroid (*n*=59, dark blue) versus PTC (*n*=501, red) in the THCA TCGA dataset. Relative gene expression in PTC versus normal thyroid is highlighted, i.e. increased (yellow), reduced (green) or not significantly different (light blue). Top ten autophagy and PT/VT genes with most significant differences in expression (N v C) are shown. NS, not significant; **, *P*<0.01; ***, *P*<0.001; *P*-values adjusted using the Benjamini-Hochberg FDR correction procedure.

**Gene expression correlation matrix (THCA, *n* = 501)**

### Figure S14

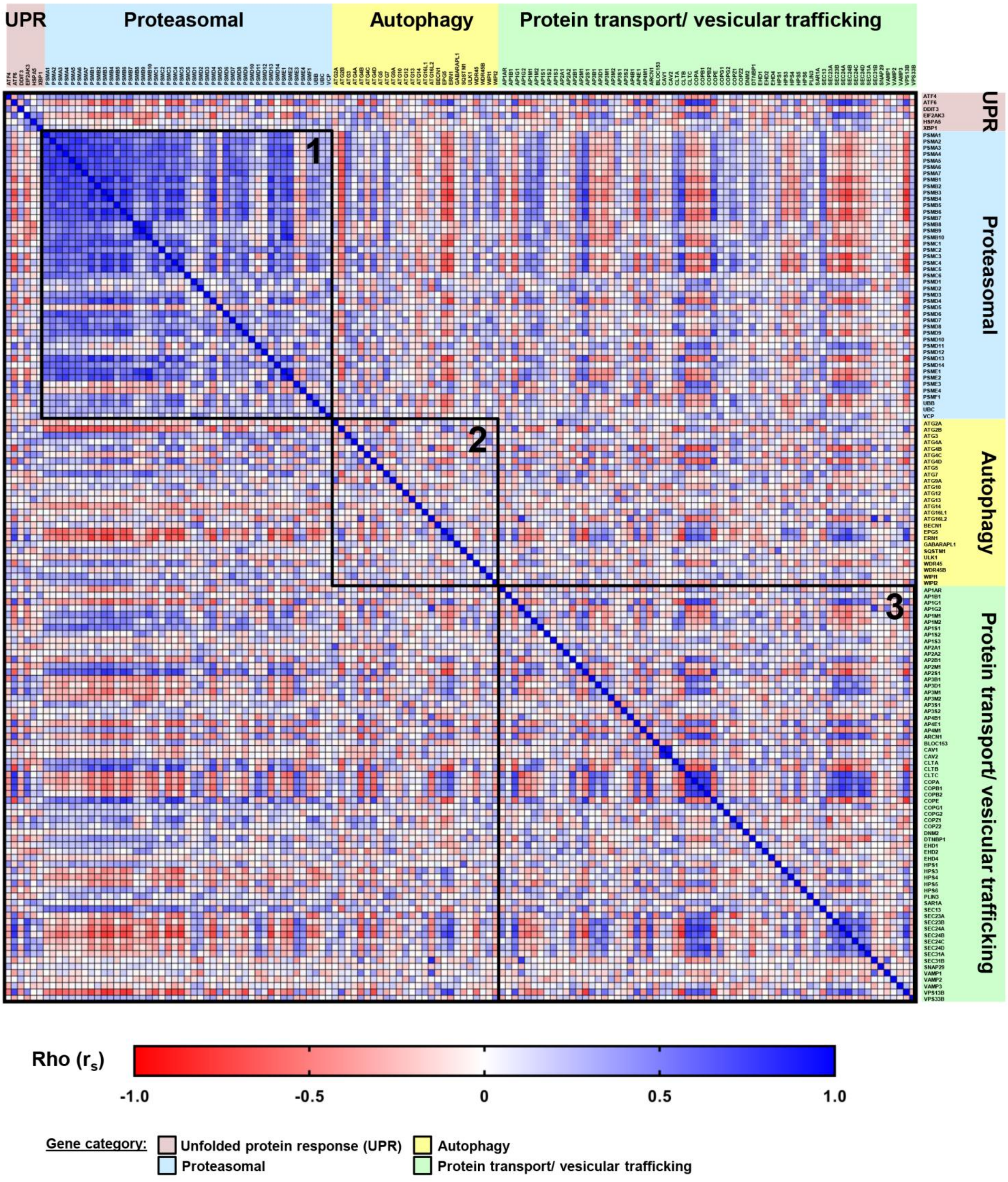


**Fig. S14. A striking correlation pattern between proteasomal genes in PTC.**
Gene expression correlation matrix of PTC in THCA TCGA (*n*=501) based on expression of 142 proteostasis-related genes divided into 4 categories: (i) unfolded protein response (UPR) genes (pink), *n*=6; (ii) proteasomal genes (blue), *n*=45; (iii) autophagy genes (yellow), *n*=26, and (iv) protein transport/ vesicular trafficking genes (green), *n*=65. Rho ($r_s$) values determined using Spearman's correlation test. Individual submatrices for proteasomal (box 1), autophagy (box 2) and protein transport/ vesicular trafficking (box 3) genes are indicated.

### Figure S15

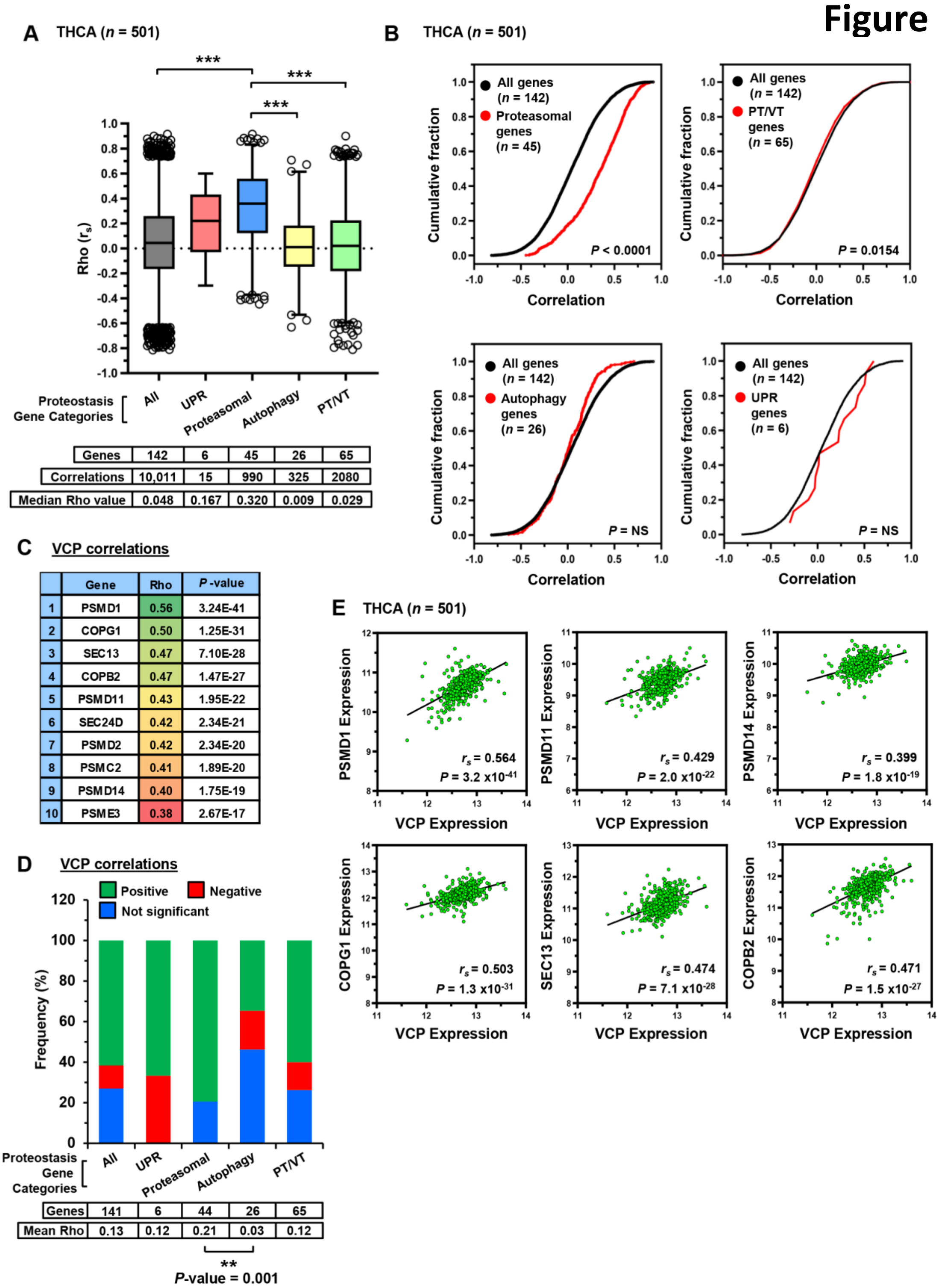


**Fig. S15. Significant and greater correlation of proteasomal genes in PTC.**
(*A*) Box and whisker plots comparing Rho values for 142 proteostasis genes subdivided into unfolded protein response (UPR), proteasomal, autophagy, and protein transport/ vesicular trafficking (PT/VT) gene categories. Number of correlations per gene category and median rho values are shown. ***, *P*<0.001; Kruskal-Wallis test followed by Dunn's post hoc test. (*B*) Cumulative frequency distribution plot comparing the correlation of 142 proteostasis genes versus different gene categories as described in (*A*); *P*-values determined by Kruskal-Wallis test followed by Dunn's post hoc test. (*C*) Top ten most significant correlations between VCP versus 141 proteostasis genes. *P*-values adjusted using the Benjamini-Hochberg FDR correction procedure. (*D*) 100% stacked bar chart showing frequency of correlations between VCP and proteostasis genes subdivided into 4 categories as described in (*A*). *P*-value determined using Fisher's exact test. (*E*) Correlation of VCP expression with indicated genes in THCA TCGA (Spearman's correlation test). *P*-values adjusted using the Benjamini-Hochberg FDR correction procedure.

### Figure S16

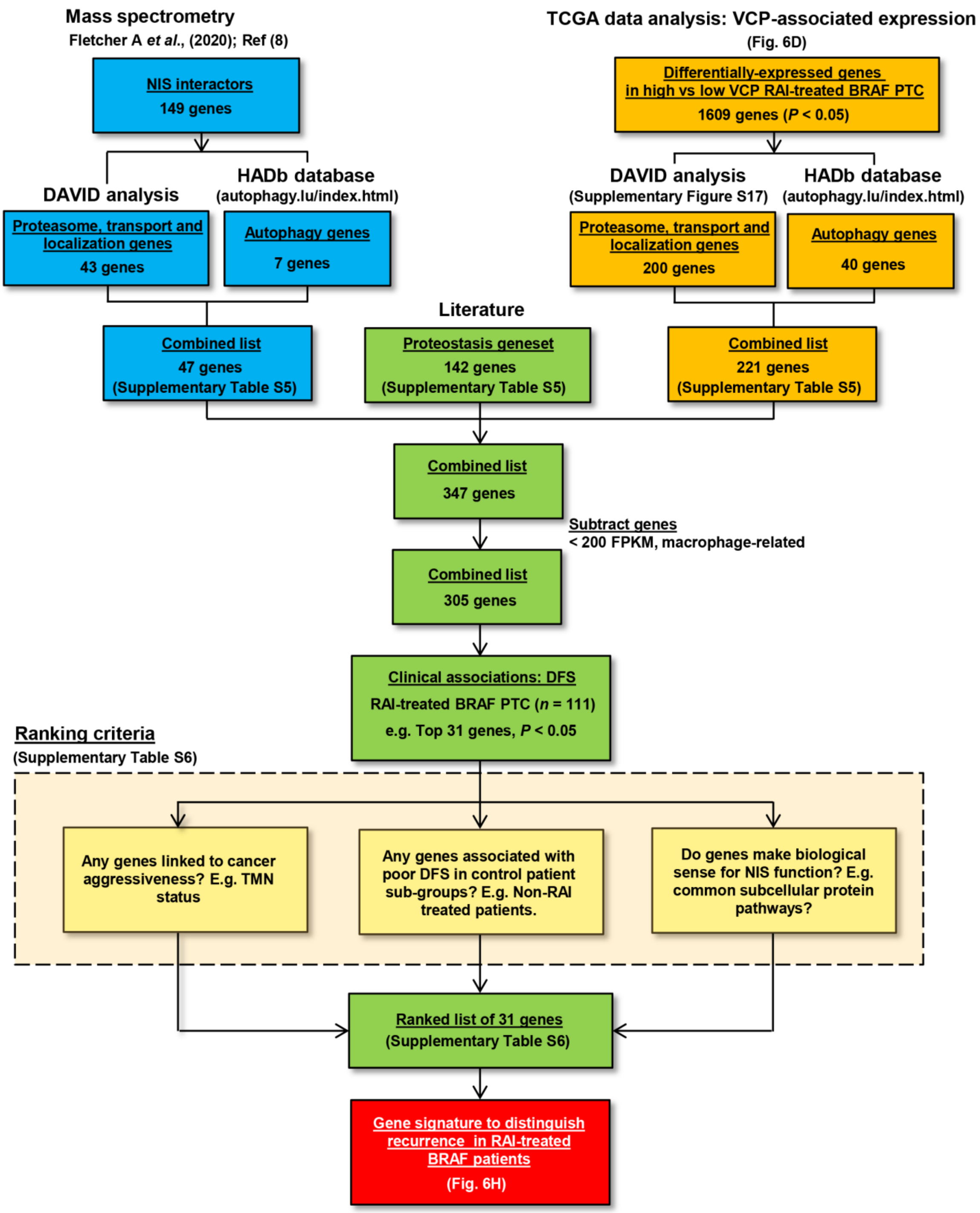


**Fig. S16. Evaluation of a focussed gene panel to identify clinically relevant drug targets in RAI-treated PTC.**
Proteostasis genes were examined to identify potential drug targets with clinical relevance linked to NIS function and disease recurrence. 347 genes were initially pooled and filtered based on DAVID functional classification (version 6.7) and the HADb database from 3 sources: (i) NIS interactors recently detected by mass spectrometry, (ii) a well-characterized proteostasis 142 geneset based on literature, and (iii) VCP-associated genes in RAI-treated BRAF THCA patients with reduced DFS in TCGA. Known macrophage-related or poorly expressed genes were removed prior to evaluation of associated clinical data in TCGA. In particular, 31 proteostasis genes significantly associated with a higher rate of recurrence in RAI-treated patients were considered top hits. Genes were ranked based on different criteria (e.g. clinical staging, patient control groups and established biological activities) and further used to generate signatures to distinguish recurrence in RAI-treated BRAF patients.

**Figure S17**

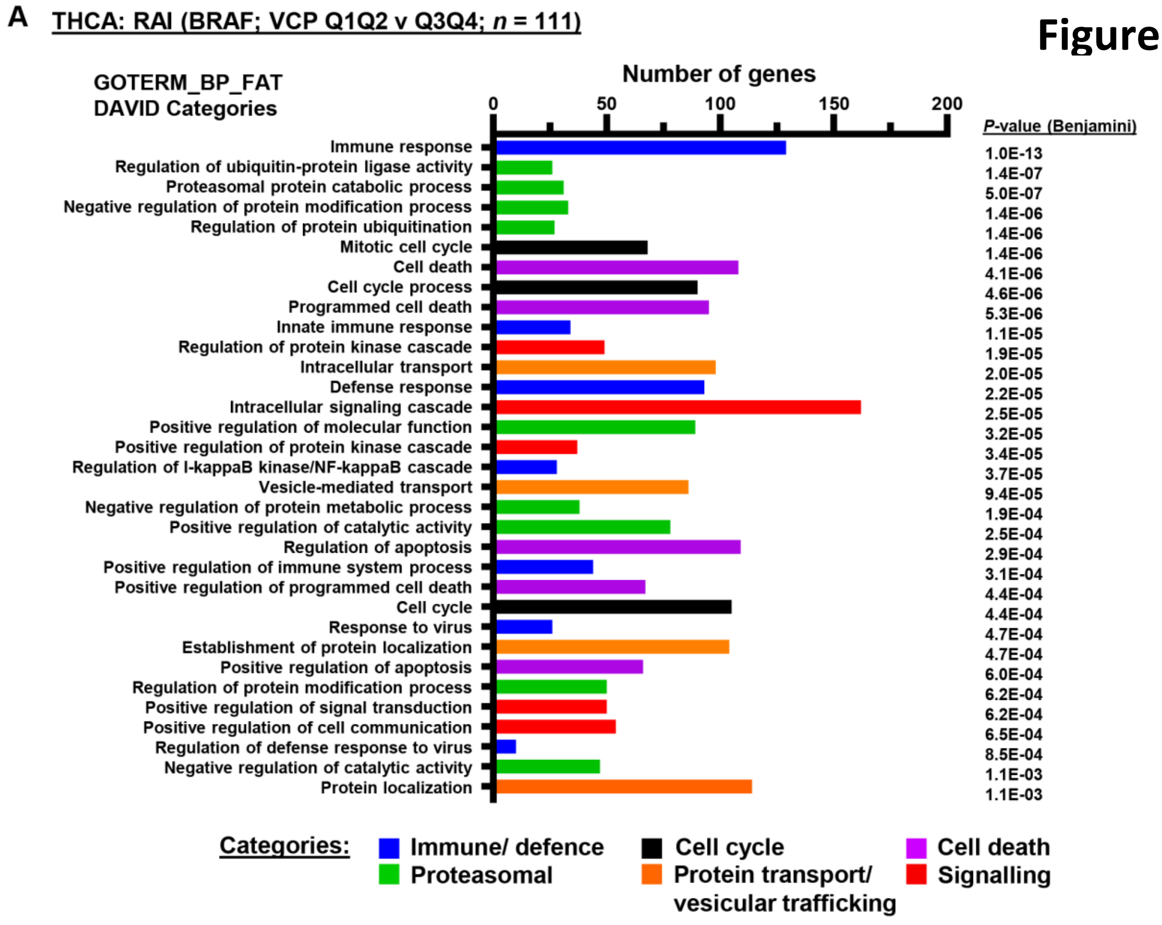


**B**

| GOTERM_BP_FAT (1609 Genes; *P* ≤ 0.05) | *P*-Value | Genes |
|---|---|---|
| Proteasomal:<br>Proteasomal protein catabolic process | 5.0E-07 | *DERL1, DERL3, FBX06, SMURF1, CDC20, NPLOC4, PSMC2, PSMD1, PSMD11, PSMD14, PSMD2, PSMD3, PSMD6, PSMD7, PSMD8, PSMD9, PSME1, PSME2, PSME3, PSMA2, PSMA4, PSMA5, PSMA6, PSMA7, PSMB10, PSMB2, PSMB5, PSMB7, PSMC4, UBC, VCP* |
| **GOTERM_BP_FAT (1609 Genes; *P* ≤ 0.05** | ***P*-Value** | **Genes** |
| Protein transport/ vesicular trafficking:<br>Intracellular transport | 2.0E-05 | *ABCG1, AP1B1, AP1M1, AP1S1, AP1S3, AP2A1, AP2A2, AP2B1, AP2M1, AP2S1, ARFGAP1, ARRB1, ARSB, ATG4A, BAK1, BAX, BCL2L1, CD209, CHMP7, COPB2, COPE, CSPG5, CUX1, DCLK1, DERL1, DST, EHD1, EIF5A, EIFAL1, ERP29, F2R, FKBP1B, FLNA, GIPC1, GOLGA3, GRIK2, HAP1, HGS, HOMER3, HSP90AA1, HSPA8, ICMT, KDELR2, KIF4A, KLC2, KPNA2, MYL6, MYO9B, NCBP1, NDEL1, NRBP1, NXT1, OPTN, PCNA, PML, POLA2, PPIF, PREB, PTTG1IP, RAB35, RAB6B, RAB8A, RAN, RTP4, SAR1A, SCAMP2, SCYL1, SEC13, SEC24C, SEC24D, SEC61A1, SEC61G, SIL1, SLC25A20, SMURF1, SSR4, STARD3, STX5, SYNGR1, SYTL1, SYTL2, SYTL5, TAP2, TGFB1, TGFB2, THOC5, TIMM17B, TOMM40, TP53, TRPC4AP, TSPO, TXNDC5, UXT, VCP, WIPI1, YKT6, YWHAB, YWAH, YWHAZ* |

**Fig. S17. DAVID functional classification to identify proteostasis genes in PTC.**
(*A*) DAVID analysis of differentially expressed genes in RAI-treated BRAF THCA patients with high (Q3Q4, *n*=56) versus low (Q1Q2, *n*=55) tumoral VCP expression. GOTERM_BP_FAT categories of greatest significance (i.e. *P*=1.1E-03 or lower) and number of genes per category are indicated. *P*-values adjusted using the Benjamini FDR correction procedure. (*B*) Detailed gene lists are shown for proteostasis-related GOTERM_BP_FAT categories, including Proteasomal protein catabolic process (Proteasomal; *P*=5.0E-07) and Intracellular transport (Protein transport/ vesicular trafficking; *P*=2.0E-05).

#### Figure S18

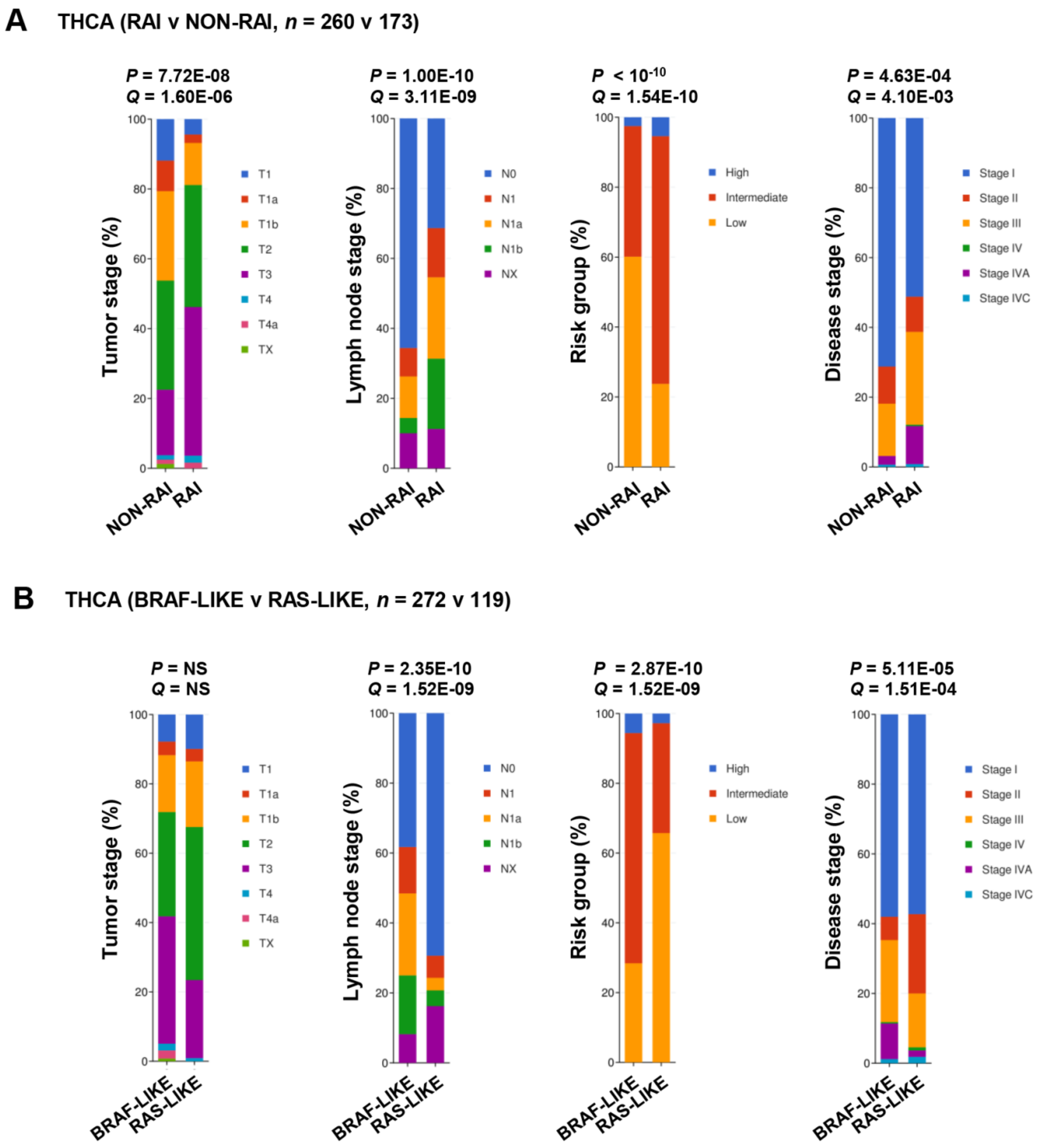


**Fig. S18. Disease classification of THCA patients stratified by RAI treatment and tumoral genetic alteration.**
(*A*) 100% stacked bar charts showing clinical characteristics of RAI (*n*=260) versus non-RAI (*n*=173) treated THCA patients in TCGA. Clinical staging attributes include: (i) tumor staging, (ii) lymph node staging, (iii) risk group and (iv) disease staging. *P*-values from cBioportal and derived using Chi-Squared test. *Q*-values derived from Benjamini-Hochberg FDR correction procedure. (*B*) Same as (*A*) but THCA patients stratified by BRAF-like (*n*=272) or RAS-like (*n*=119) tumoral genetic alteration. NS, not significant.

#### Figure S19

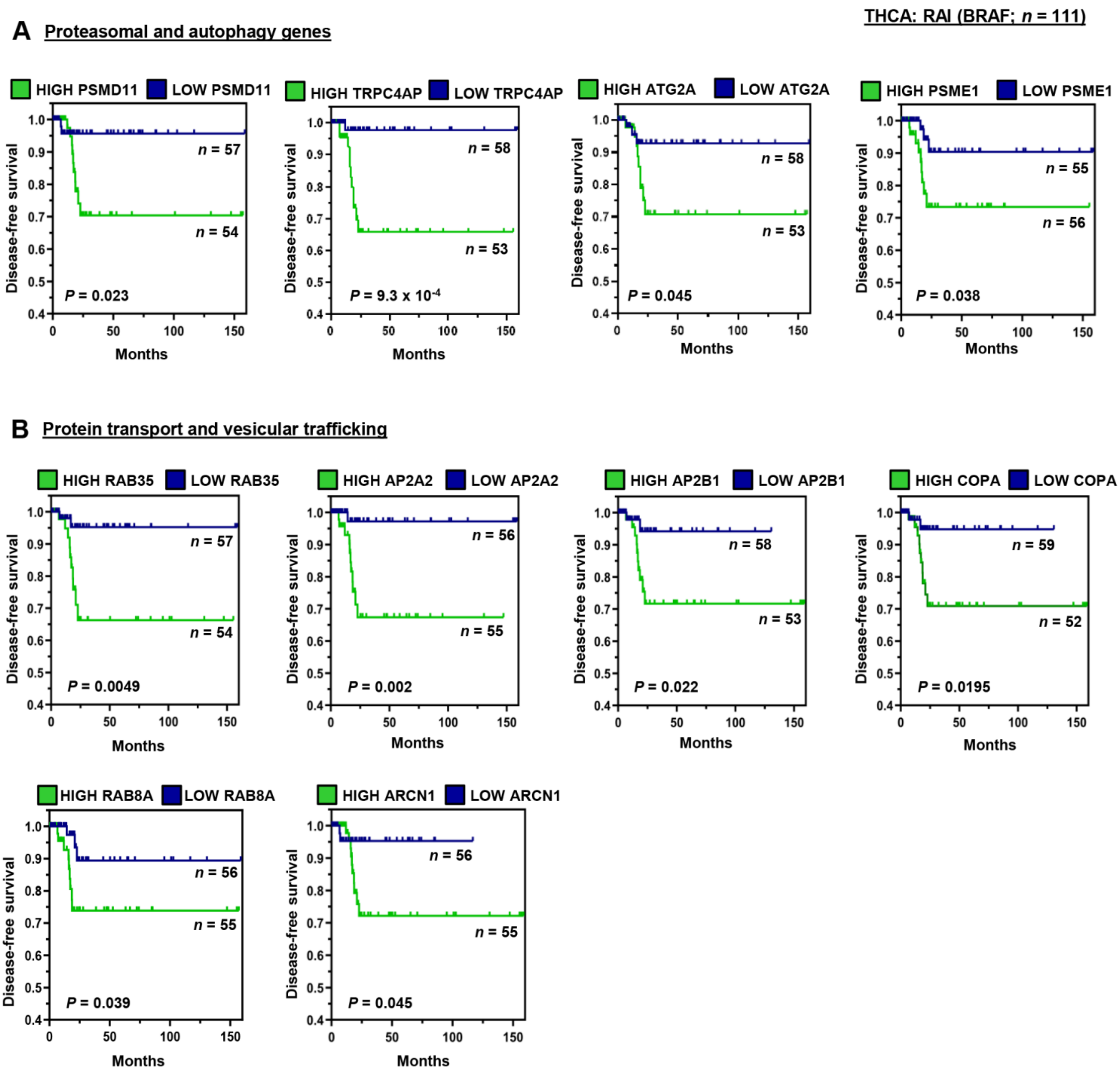


**Fig. S19. Proteostasis-related genes correlate with poorer survival in RAI-treated BRAF patients.**

(*A*) Disease-free survival (DFS) for RAI-treated BRAF patients (*n*=111) with high (Q3Q4) versus low (Q1Q2) expression of four proteasomal/ autophagy genes (i.e. *PSMD11*, *TRPC4AP*, *ATG2A* and *PSME1*). Log-rank *P*-values are shown. (*B*) Same as (*A*) but with protein transport/ vesicular trafficking genes (i.e. *RAB35*, *AP2A2*, *AP2B1*, *COPA*, *RAB8A* and *ARCN1*).

#### Figure S20

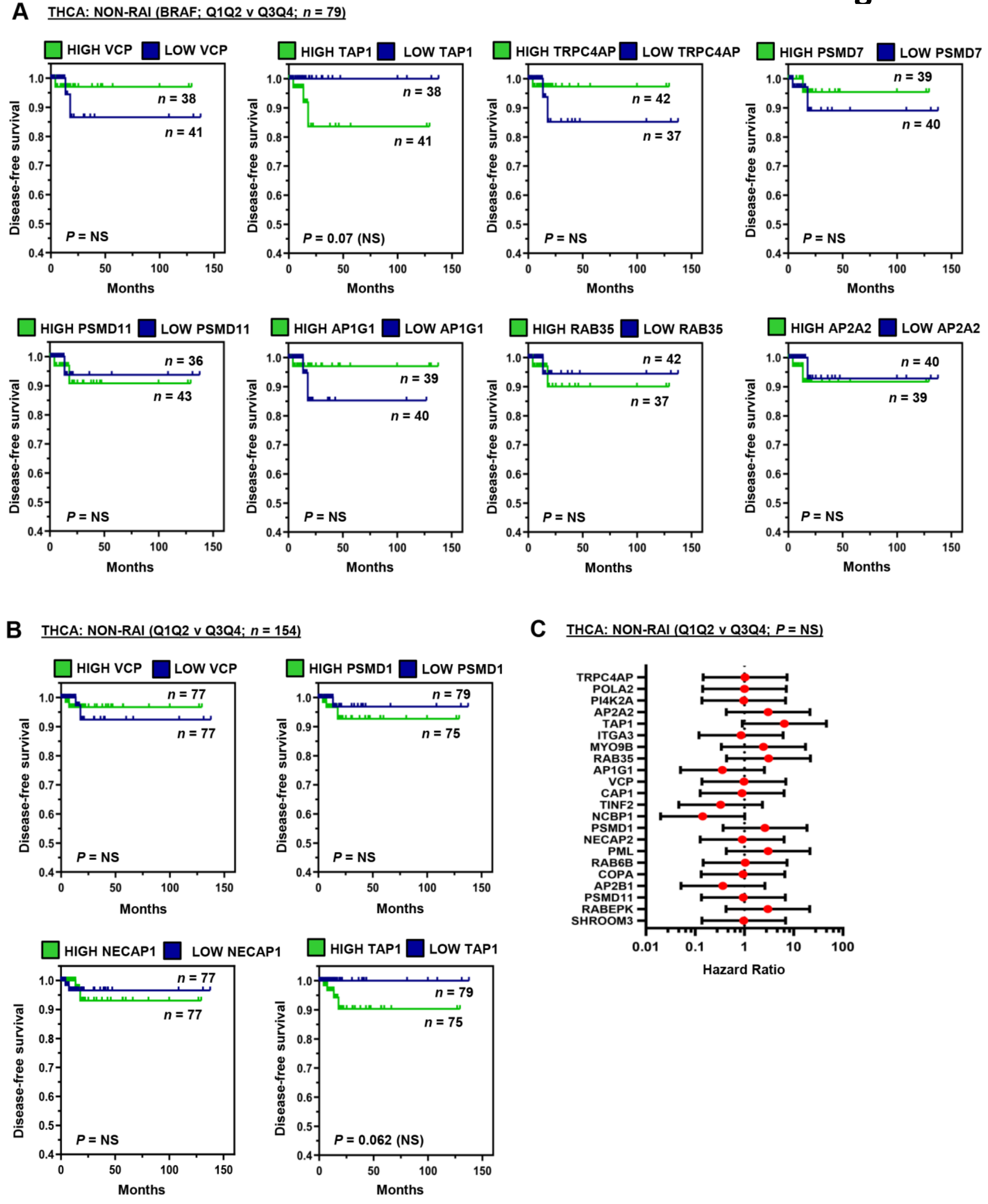


**Fig. S20. Lack of association between non-RAI treated THCA patient survival and proteostasis-related genes.**

(*A*) DFS for non-RAI treated BRAF patients (*n*=79) with high (Q3Q4) versus low (Q1Q2) expression of 8 proteostasis-related genes (i.e. *VCP*, *TAP1*, *TRPC4AP*, *PSMD7*, *PSMD11*, *AP1G1*, *RAB35* and *AP2A2*). Log-rank *P*-values are shown. (*B*) DFS for non-RAI THCA patients (*n*=154; including patients with BRAF and other tumoral genetic alterations) with high (Q3Q4) versus low (Q1Q2) expression of four proteostasis-related genes (i.e. *VCP*, *PSMD1*, *NECAP1* and *TAP1*). Log-rank *P*-values are shown. (*C*) Hazard ratios ±95% CI for non-RAI treated THCA patients stratified on median tumoral expression of 22 proteostasis-related genes as indicated. NS, not significant.

**Table S1. Materials used in study.**

[1]Stable NIS and/or YFP cell lines were generated using standard protocols with FACS single cell sorting (University of Birmingham Flow Cytometry Facility) followed by antibiotic selection. Type of antibiotic and doses used are indicated for different thyroid cancer cell lines. Typically 40-50 single cell clones were successfully expanded and relative expression of either YFP (fluorescence; flow cytometry) or NIS (mRNA levels; real-time PCR) was compared. NIS function of different clones was validated by RAI (iodine-125) uptake assays compared to parental NIS negative matched cell lines. [2]Pooled reagent of three MISSION siRNAs targeted to VCP. Designed using algorithm from Rosetta Inpharmatics. Approximate positioning where siRNA targets gene are at nucleotides 768 (Hs01_00118726), 1908 (Hs02_00343009) and 2548 (Hs01_00118728). VCP ablation used to determine the extent to which certain drugs (e.g. carebastine, SAHA) were dependent on VCP inhibition to enhance RAI uptake (Fig. 4F; Supplementary Fig. S11B). siRNA transfections were performed with Lipofectamine RNAi/MAX (Thermo Fisher Scientific) following standard protocols. [3]Primary antibodies used in Western blotting. To quantify detected bands by densitometry, blots were scanned, keeping all scanning parameters the same and analyzed using ImageJ software (e.g. Fig. 2C and E). [4]List of commercial Taqman® assays (Thermo Fisher Scientific) used in study. PPIA was used as internal control. Relative expression calculated using the $2^{-\Delta\Delta Ct}$ method. For cDNA synthesis, total RNA was extracted using the RNeasy Micro Kit (Qiagen) prior to reverse transcription using the Reverse Transcription System (Promega) in accordance with the manufacturer's guidelines.

| **Vector[1]** | **Gene** | **Source** | **Notes** |
|---|---|---|---|
| pcDNA3.1 Hygro | EYFP-H148Q/I152L/F46L | Dr Peter Haggie<br>Prof Alan Verkman<br>(UCSF) | Previously described<br>pcDNA3.1 Hygro EYFP H148Q/I152L<br>(Galietta LJ *et al*., 2001)<br>150 µg/ml hygromycin B (TPC-1/8505C) |
| pcDNA3.1 Geneticin | NIS-HA | Previously described<br>(Smith VE *et al*., 2013) | 300 µg/ml geneticin (TPC-1)<br>500 µg/ml geneticin (8505C) |
| **Target[2]** | **Type of siRNA** | **Company** | **ID** |
| VCP | MISSION siRNA (pooled) | Sigma-Aldrich | Hs01_00118726; Hs01_00118728;<br>Hs02_00343009 |
| Scrambled (Scr) | Negative control siRNA | Thermo Fisher Scientific | AM4635 |
| **Antibody[3]** | **Clone** | **Supplier** | **Dilution** |
| β-actin | Mouse monoclonal [AC-15] | Sigma-Aldrich | 1:2000 |
| HA | Mouse monoclonal [16B12] | Legend | 1:1000 |
| LC3B | Rabbit polyclonal [#2775] | Cell Signalling Technology | 1:1000 |
| NIS | Rabbit polyclonal [24324-1-AP] | ProteinTech | 1:1000 |
| VCP | Rabbit polyclonal [10736-1-AP] | ProteinTech | 1:1000 |

| **Commercial Taqman Assays[4]** | |
|---|---|
| Gene target | ID |
| *NIS* | Hs00166567_ml |
| *VCP* | HS00997642_m1 |
| *PPIA* | Hs04194521_s1 |

**Table S2. Compounds used in study.**

[1]List of individual drugs and drug libraries used in study. Chloroquine diphosphate was resuspended in PBS without calcium/magnesium (Thermo Fisher Scientific); Phenformin HCL in 100 % ethanol and all other drugs in dimethyl sulfoxide (DMSO; Sigma-Aldrich), before being diluted in RPMI-1640 medium (Life Technologies) to appropriate concentrations and added directly to cells at 1:100 dilution. [2]Drugs purchased from the indicated company. [3]Positive control drugs reported to enhance NIS activity were used to validate the YFP-iodide and radioiodine uptake assays. [4]Negative control wells consisted of cells being treated with DMSO alone in the absence of any other drugs. [5]PBS was injected into wells by the PHERAstar FS plate reader as a negative control in the YFP-iodide assay. [6]Microlab STAR Liquid Handling Robotic System (Birmingham Drug Discovery Facility) was used to pre-load 96 well microtiter plates with 1200 drugs (Prestwick Chemical Library) prior to cell seeding. [7]NaI was injected into wells by the PHERAstar FS plate reader as a positive control in the YFP-iodide assay at a final concentration of 34 mM. [8]Cells were incubated with 100 μM sodium perchlorate for 1 hour at 37ºC prior to addition of iodine-125 to validate NIS-mediated radioiodine uptake.

| Drug[1] | Company[2] | Notes/ (Abbreviation) |
|---|---|---|
| Astemizole | Sigma-Aldrich | (AST) |
| Carebastine | Sigma-Aldrich | (CBT) |
| Chloroquine diphosphate | Sigma-Aldrich | (CQ) |
| Clotrimazole | Sigma-Aldrich | (CLOT) |
| DBeQ | Sigma-Aldrich | Positive control[3] |
| Dimethyl sulfoxide | Sigma-Aldrich | Negative control[4]; (DMSO) |
| Disulfiram | Sigma-Aldrich | (DSF) |
| Dynasore | Sigma-Aldrich | Positive control[3] |
| Ebastine | Sigma-Aldrich | (EBT) |
| Eeyarestatin-1 (ES-1) | Cayman Chemicals | Positive control[3]; (ES-1) |
| Fexofenadine HCL | Sigma-Aldrich | |
| Formoterol hemifumarate | Stratech Scientific | |
| Imatinib mesylate | Cambridge Bioscience | (IMAT) |
| Niflumic acid | Alfa Aesar | (NFA) |
| NMS-873 | SelleckChem | Positive control[3] |
| Phenformin HCL | Stratech Scientific | |
| Phosphate buffered saline | SelleckChem | Negative control[5]; (PBS) |
| Pravastatin sodium salt | Cambridge Bioscience | |
| Prestwick Chemical Library (1200 drugs) | Prestwick Chemical | Birmingham Drug Discovery Facility[6] |
| SAHA/ Vorinostat | Stratech Scientific | |
| Selumetinib | SelleckChem | Positive control[3]; (SEL) |
| Sodium iodide | Sigma-Aldrich | Positive control (34mM)[7]; (NaI) |
| Sodium perchlorate | Sigma-Aldrich | Negative control[8]; ($NaClO_4$) |
| Sulconazole nitrate | Sigma-Aldrich | |
| Terfenadine | Sigma-Aldrich | |
| Vatalanib (PTK787) 2HCL | Stratech Scientific | |
| Vemurafenib | Sigma-Aldrich | Positive control[3]; (VEM) |
| Zimelidine 2HCL | Sigma-Aldrich | |

**Table S3. High proportion of drugs in the primary screen target the proteostasis network.** List of 48 drugs identified in the primary screen which increased intracellular iodide (ΔYFP > 2; Fig. 1D). Known categories for each drug are shown, as well as any reported relevant proteostasis targets or related mechanisms of action (highlighted in green). Supplementary references are provided to support categorization of drugs as targeting the proteostasis network (Supplementary References).

| Drug | Category | Relevant proteostasis target(s) or action(s), if known | Supplementary Reference |
|---|---|---|---|
| Atenolol | Beta blocker | N/A | |
| Antimycin A | Piscicide | Autophagy | (13) |
| Articaine | Anesthetic | N/A | |
| Astemizole | Antifungal | Lysosomotrophic, VCP inhibitor (ERAD) | (11, 14) |
| Bromocryptine | Dopamine agonist | Autophagy | (15) |
| Ceforanide | Antibiotic | N/A | |
| Chloroquine | Antimalarial | Autophagy, lysosomotrophic, vesicular transport | (14, 18) |
| Clofilium | Anti-arrhythmia | N/A | |
| Clomipramine | Antidepressant | Autophagy, lysosomotrophic | (14) |
| Clotrimazole | Antifungal | VCP inhibitor (ERAD) | (11) |
| Demecarium | Anticholinesterase | N/A | |
| Dequalinium | Antimicrobial | N/A | |
| Ebastine | Antihistamine | VCP inhibitor (ERAD) | (11) |
| Eucatropine | Anticholinergic | N/A | |
| Fendiline | Calcium channel blocker | N/A | |
| Fenofibrate | Lipid-lowering | Autophagy | (16) |
| Formoterol | β(2) -agonist | Proteasomal, autophagy | (17) |
| Halofantrine | Antimalarial | Autophagy, lysosomotrophic | (18) |
| Hesperidin | Flavanone glycoside | Autophagy, ER stress | (19) |
| Imatinib | Anticancer | Autophagy, lysosomotrophic, tyrosine kinase inhibitor | (20) |
| Itopride | Dopamine D2 antagonist | N/A | |
| Lithocholic acid | Detergent | Autophagy, ER stress | (21) |
| Meclozine | Antihistamine | Histamine H1 antagonist | |
| Methotrimeprazine | Phenothiazine | N/A | |
| Methyldopate | Antihypertensive | N/A | |
| Mifepristone | Antiprogestogenic | Autophagy, unfolded protein response | (22) |
| Nomegestrol | Progestogen | N/A | |
| Nortriptyline | Antidepressant | Autophagy, lysosomotrophic | (14) |
| Oxantel | Anthelminthic | N/A | |
| Oxyphenbutazone | Anti-Inflammatory | N/A | |
| Phenformin | Antidiabetic | Autophagy, mTOR | (23) |
| Phentolamine | α-adrenergic antagonist | N/A | |
| Piperacetazine | Antipsychotic | N/A | |
| Pravastatin | Lipid-lowering | HMG-CoA reductase inhibitor | |
| Prednisolone | Anti-Inflammatory | N/A | |
| Propidium | Intercalation | N/A | |
| Pyrvinium | Anthelminthic | Autophagy | (24) |
| Quinidine | Anti-arrhythmia | Lysosomotrophic | (25) |
| Rivastigmine | Anticholinesterase | N/A | |
| Rosiglitazone | Hypoglycemic | Autophagy, mTOR | (26) |
| Rufloxacin | Antibiotic | N/A | |
| Sisomicin | Antibiotic | N/A | |
| Terfenadine | Antihistamine | VCP inhibitor (ERAD) | (11) |
| Tioconazole | Antifungal | Autophagy (ATG4B inhibitor) | (27) |
| Tosufloxacin | Antibiotic | N/A | |
| Trimethoprim | Antibiotic | N/A | |
| Vanoxerine | Dopamine reuptake | N/A | |
| Vatalanib | Anticancer | Tyrosine kinase inhibitor | |

**Table S4. Comparison of drug-induced RAI uptake in TPC-1 cells and human thyrocytes at maximal responses.**

[1]Number of biological replicates performed for each drug treatment. Each biological replicate typically consisted of 3-4 technical replicates from independent experiments unless otherwise stated. [2]Identified dose or range of doses for each drug that gave the greatest increase in RAI uptake. [3]Mean fold-increase in RAI uptake ± SE for each drug relative to 0.1% DMSO as vehicle control, except for chloroquine (0.5% PBS), formoterol (0.2% DMSO), pravastatin (0.4% DMSO) and phenformin (0.25% ETOH). [4]The two-tailed t-test was used to determine *P*-values for all drug treatments, except in human primary thyrocytes treated with ebastine, clotrimazole and astemizole in which the non-parametric Mann-Whitney test was used. Significant *P*-values ($P \leq 0.05$) with RAI uptake > 1.5-fold are highlighted (red). [5]Data from Fletcher A *et al.* 2020 (Ref 8) included for comparison. Fold-increase in RAI uptake in primary thyrocytes determined from maximal responses using a range of drug doses. [6]Cells treated for 24 hours prior to assaying RAI uptake, except for chloroquine in which cells were treated for 8 hours. [7]N/A: data not available.

| Drugs | TPC-1 (NIS+ve) cells | | | | Human primary thyrocytes | | | |
|---|---|---|---|---|---|---|---|---|
| | n[1] | Dose (μM)[2] | RAI uptake[3] | *P*-value[4] | n[1] | Dose (μM)[2] | RAI uptake[3] | *P*-value[4] |
| Ebastine | 3 | 0.5 | 4.6 ± 0.4[5] | 0.002 | 24 | 0.05-0.25 | 2.5 ± 0.4[5] | 2.0E-04 |
| SAHA | 5 | 5 | 4.3 ± 0.2 | 5.5E-07 | 6 | 5 | 1.6 ± 0.1 | 6.9E-07 |
| Carebastine | 3 | 1 | 3.8 ± 0.4 | 0.003 | 3 | 0.75 | 3.1 ± 0.6 | 0.04 |
| Formoterol | 3 | 10 | 3.1 ± 0.2 | 0.001 | 3 | 1 | 2.2 ± 0.1 | 5.4E-06 |
| Clotrimazole | 4 | 0.25 | 2.4 ± 0.2[5] | 0.002 | 24 | 0.05-0.75 | 2.3 ± 0.5[5] | 0.006 |
| Astemizole | 3 | 0.25 | 2.2 ± 0.2[5] | 0.004 | 11 | 0.05-0.75 | 2.9 ± 0.7 | 0.016 |
| Disulfiram | 5 | 0.5 | 2.2 ± 0.1 | 0.001 | 6 | 0.5 | 2.5 ± 0.1 | 8.9E-06 |
| Phenformin | 3 | 250 | 1.9 ± 0.1 | 0.001 | 3 | 50 | 1.3 ± 0.1 | NS |
| Imatinib | 4 | 1 | 1.9 ± 0.2 | 0.004 | 3 | 1 | 2.6 ± 0.1 | 0.023 |
| Niflumic acid | 4 | 15 | 1.7 ± 0.1 | 4.0E-04 | 4 | 20 | 1.8 ± 0.1 | 2.4E-05 |
| Vatalanib | 4 | 1 | 1.6 ± 0.1 | 0.002 | 3 | 10 | 3.4 ± 0.3 | 0.002 |
| Pravastatin | 3 | 40 | 1.6 ± 0.1 | 0.003 | 3 | 40 | 1.9 ± 0.2 | 0.009 |
| Chloroquine[6] | 6 | 50 | 1.2 ± 0.1 | NS | 3 | 30 | 2.0 ± 0.4 | 0.012 |
| Zimelidine | 3 | 1 | 1.1 ± 0.01 | 0.003 | 3 | 1 | 1.9 ± 0.1 | 0.004 |
| Terfenadine | N/A[7] | N/A | N/A | N/A | 3 | 1 | 3.5 ± 0.6 | 0.027 |
| Fexofenadine | N/A | N/A | N/A | N/A | 3 | 2 | 4.1 ± 1.0 | 0.047 |

**Table S5. Proteostasis genes used in focused gene panel to study clinical relevance.**
List of proteostasis genes used to identify potential drug targets or pathways with clinical relevance (Fig. S16). Three genesets used: (i) 142 well-characterized proteostasis genes comprising of 6 unfolded protein response (UPR), 45 proteasomal, 26 autophagy and 65 protein transport/ vesicular trafficking genes; (ii) 47 proteostasis genes filtered by DAVID functional classification or entry in HADb database (autophagy.lu/index.html) from a panel of 149 NIS interactors (Fletcher A *et al.*, 2020), and (iii) 221 proteostasis genes filtered by DAVID functional classification or entry in HADb database from 1609 VCP-associated genes identified in RAI-treated BRAF THCA patients in TCGA (*, $P$<0.05; adjusted using the Benjamini-Hochberg FDR correction procedure; Fig. 6D).

| Gene lists | No. of genes | Category | Genes |
|---|---|---|---|
| Proteostasis geneset (142 genes in total) | 6 | Unfolded protein response (UPR) | *ATF4, ATF6, DDIT3, EIF2AK3, HSPA5, XBP1* |
| | 45 | Proteasomal/ ubiquination | *PSMA1, PSMA2, PSMA3, PSMA4, PSMA5, PSMA6, PSMA7, PSMB1, PSMB2, PSMB3, PSMB4, PSMB5, PSMB6, PSMB7, PSMB8, PSMB9, PSMB1, PSMC1, PSMC2, PSMC3, PSMC4, PSMC5, PSMC6, PSMD1, PSMD2, PSMD3, PSMD4, PSMD5, PSMD6, PSMD7, PSMD8, PSMD9, PSMD10, PSMD11, PSMD12, PSMD13, PSMD14, PSME1, PSME2, PSME3, PSME4, PSMF1, UBB, UBC, VCP* |
| | 26 | Autophagy | *ATG2A, ATG2B, ATG3, ATG4A, ATG4B, ATG4C, ATG4D, ATG5, ATG7, ATG9A, ATG10, ATG12, ATG13, ATG14, ATG16L1, ATG16L2, BECN1, EPG5, ERN1, GABARAPL1, SQSTM1, ULK1, WDR45, WDR45B, WIPI1, WIPI2* |
| | 65 | Protein transport/ vesicular trafficking | *AP1AR, AP1B1, AP1G1, AP1G2, AP1M1, AP1M2, AP1S1, AP1S2, AP1S3, AP2A1, AP2A2, AP2B1, AP2M1, AP2S1, AP3B1, AP3D1, AP3M1, AP3M2, AP3S1, AP3S2, AP4B1, AP4E1, AP4M1, ARCN1, BLOC1S3, CAV1, CAV2, CLTA, CLTB, CLTC, COPA, COPB1, COPB2, COPE, COPG, COPG2, COPZ1, COPZ2, DNM2, DTNBP1, EHD1, EHD2, EHD4, HPS1, HPS3, HPS4, HPS5, HPS6, PLIN3, SAR1A, SEC13, SEC23A, SEC23B, SEC24A, SEC24B, SEC24C, SEC24D, SEC31A, SEC31B, SNAP29, VAMP1, VAMP2, VAMP3, VPS13B, VPS33B* |
| NIS interactors (Fletcher A *et al.*, 2020) | 149 | 47 filtered by DAVID analysis and HADb database | *AHSG, AP2B1, APOC3, APOE, ARCN1, ARF4, ARF6, C3, CANX, CAPN2, CAV1, COPA, COPB1, COPG2, COPZ1, CTSB, CTSD, DMBT1, EHD1, GARS, GNB2L1, HUWE1, MVP, MYBBP1A, NCALD, PDCD6IP, PICALM, RAB14, RAB18, RAB1C, RAB5C, RAC3, RHEB, SAR1A, SEC16A, SHROOM3, SLC25A6, TLN1, UBE2D3, UBE2M, UBE2N, UBE2V1, UBE2V2, VCP, YKT6, YWHAE, YWHAZ* |
| VCP-associated genes: differential expression, TCGA analysis (*P*-value < 0.5) | 1609 | 221 filtered by DAVID analysis and HADb database | *ABCG1, ACTN4, ACTR1A, ANK1, AP1B1, AP1M1, AP1S1, AP1S3, AP2A1, AP2A2, AP2B1, AP2M1, AP2S1, APLP1, APOL1, ARF4, ARFGAP1, ARRB1, ATG16L1, ATG4A, ATG7, ATG9A, BACE2, BAK1, BAX, BCL2L1, BNIP1, CAP1, CAPN1, CAPNS1, CASP1, CCL2, CCL3, CCL8, CD14, CD209, CDC20, CDKN1A, CHMP1A, CHMP7, COPB2, COPE, CORO1A, CSPG5, CUX1, CX3CL1, CXCL10, CXCL16, DBNL, DCLK1, DERL1, DERL3, DNAJB1, DRAM1, DST, EHD1, EHD2, EHD4, EIF4G1, EIF5A, EIF5AL1, ERBB2, ERGIC3, ERP29, F2R, FAS, FBXO6, FCGR1A, FCN1, FKBP1B, FLNA, FZR1, GAPDH, GARS, GIPC1, GOLGA3, GRIK2, GSN, HAP1, HGS, HOMER3, HRAS, HSP90AA1, HSPA8, HSPB8, ICA1, ICMT, ITGA2, ITGA3, ITGB4, KDELR2, KDELR3, KIF4A, KLC2, KPNA2, LAT2, LDLR, LMAN2, LRP10, MTMR14, MVP, MYL6, MYO1C, MYO9B, NACAD, NCBP1, NDEL1, NECAP2, NLRC4, NME2, NPLOC4, NRBP1, NUP85, NUP93, NXT1, OPTN, P2RX1, P4HB, PCNA, PI4K2A, PITPNM1, PLD2, PLIN3, PML, POLA2, PPIF, PPP2R1A, PREB, PRKCB, PSMA2, PSMA4, PSMA5, PSMA6, PSMA7, PSMB10, PSMB2, PSMB5, PSMB7, PSMC2, PSMC4, PSMD1, PSMD11, PSMD14, PSMD2, PSMD3, PSMD6, PSMD7, PSMD8, PSMD9, PSME1, PSME2, PSME3, PTTG1IP, RAB11FIP5, RAB20, RAB31, RAB32, RAB35, RAB36, RAB4B, RAB6B, RAB8A, RABEPK, RABIF, RAC1, RAN, RBM12, RGS19, RIN3, RTP4, SAR1A, SCAMP2, SCG2, SCYL1, SEC13, SEC24C, SEC24D, SEC61A1, SEC61G, SESN2, SFT2D2, SH3GL1, SIL1, SLC15A3, SLC25A20, SMURF1, SNF8, SNX8, SOCS1, SSR4, STARD3, SYNGR1, TAP1, TAP2, TGFB1, TGFB2, THOC5, TIMM17B, TINF2, TMED3, TNFSF13B, TOMM40, TP53, TP73, TRIM36, TRPC4AP, TSPO, TXNDC5, UBC, UNC13B, UNC13D, UXT, VAV1, VCP, VPS25, WIPI1, YIF1A, YKT6, YWHAB, YWHAH, YWHAZ* |

**Table S6. Ranking of proteostasis-related genes associated with reduced DFS.**
[1]Source of proteostasis gene entry as indicated (blue-filled square; see Table S3). [2]Genes #1 to 20 were ranked on statistical significance of reduced survival (DFS) and/or increased risk of recurrence (hazard ratio) for RAI-treated BRAF patients with high (Q3Q4) versus low (Q1Q2) proteostasis gene expression. Genes were ranked lower (e.g. *CAP1*) if there were any significant differences in clinical staging attributes (i.e. tumor, lymph node and disease) or risk group allocation, which might infer a direct involvement in the advancement of thyroid disease. Importantly, there were no significant differences in lymph node or disease staging between patient cohorts for any of the 31 genes. [3]Statistical significance in DFS for non-RAI treated patients, either BRAF or all genetic alterations, with high (Q3Q4) versus low (Q1Q2) proteostasis gene expression. [4]Uniprot classifications for each proteostasis gene are shown containing both cellular component and functional information (blue-filled squares; uniprot.org). Some genes were ranked lower (e.g. *POLA2*) due to Uniprot classifications highlighting the nucleus as the primarily cellular location, which might infer a more indirect effect on NIS protein function. NS, not significant.

| | | Source[1] | | | | BRAF THCA (RAI, *n* = 130, Q1Q2 v Q3Q4)[2] | | | | | | BRAF THCA (NON-RAI)[3] | THCA (NON-RAI)[3] |
|---|---|---|---|---|---|---|---|---|---|---|---|---|---|
| | Gene | NIS interactors | Proteostasis geneset | VCP-associated/ DAVID analysis | VCP-associated/ HADb database | Disease-Free Survival (DFS) | Hazard Ratio | Tumour stage (T) | Lymph node stage (N) | Risk group | Disease stage | DFS | DFS |
| | | | | | | *P*-values | | | | | | | |
| 1 | TRPC4AP | | | ■ | | 9.30E-04 | 1.20E-02 | NS | NS | NS | NS | NS | NS |
| 2 | PI4K2A | | | ■ | | 1.80E-03 | 1.60E-02 | NS | NS | NS | NS | NS | NS |
| 3 | AP2A2 | | ■ | ■ | | 2.00E-03 | 1.60E-02 | NS | NS | NS | NS | NS | NS |
| 4 | ITGA3 | | | | ■ | 3.60E-03 | 1.20E-02 | NS | NS | NS | NS | NS | NS |
| 5 | RAB35 | | | ■ | | 4.90E-03 | 1.50E-02 | NS | NS | NS | NS | NS | NS |
| 6 | AP1G1 | | ■ | | | 6.20E-03 | 1.70E-02 | NS | NS | NS | NS | NS | NS |
| 7 | VCP | ■ | ■ | ■ | | 6.80E-03 | 2.80E-02 | NS | NS | NS | NS | NS | NS |
| 8 | PSMD1 | | ■ | ■ | | 9.40E-03 | 3.30E-02 | NS | NS | NS | NS | NS | NS |
| 9 | NECAP2 | | | ■ | | 9.70E-03 | 2.30E-02 | NS | NS | NS | NS | NS | NS |
| 10 | PML | | | ■ | | 0.0100 | 0.020 | NS | NS | NS | NS | NS | NS |
| 11 | RAB6B | | | ■ | | 0.0153 | 0.031 | NS | NS | NS | NS | NS | NS |
| 12 | AP2B1 | ■ | ■ | ■ | | 0.0221 | 0.039 | NS | NS | NS | NS | NS | NS |
| 13 | PSMD11 | | ■ | ■ | | 0.0225 | 0.040 | NS | NS | NS | NS | NS | NS |
| 14 | RABEPK | | | ■ | | 0.028 | 0.047 | NS | NS | NS | NS | NS | NS |
| 15 | SEC24C | | ■ | ■ | | 0.0371 | NS | NS | NS | NS | NS | NS | NS |
| 16 | PSME1 | | ■ | ■ | | 0.0383 | NS | NS | NS | NS | NS | NS | NS |
| 17 | CUX1 | | | ■ | | 0.0388 | NS | NS | NS | NS | NS | NS | NS |
| 18 | RAB8A | | | ■ | | 0.0399 | NS | NS | NS | NS | NS | NS | NS |
| 19 | ATG2A | | ■ | | ■ | 0.0448 | NS | NS | NS | NS | NS | NS | NS |
| 20 | ARCN1 | ■ | ■ | | | 0.0449 | NS | NS | NS | NS | NS | NS | NS |
| 21 | COPA | ■ | ■ | | | 0.0195 | 0.036 | NS | NS | 0.0324 | NS | NS | NS |
| 22 | PSMD7 | | ■ | ■ | | 0.0362 | NS | NS | NS | 0.0194 | NS | NS | NS |
| 23 | PREB | | | ■ | | 0.0445 | NS | NS | NS | 0.0175 | NS | NS | NS |
| 24 | POLA2 | | | ■ | | 1.10E-03 | 1.30E-02 | NS | NS | NS | NS | NS | NS |
| 25 | TAP1 | | | ■ | | 2.90E-03 | 1.10E-02 | 0.0112 | NS | 1.30E-03 | NS | NS | NS |
| 26 | MYO9B | | | ■ | | 4.30E-03 | 2.30E-02 | NS | NS | 0.017 | NS | NS | NS |
| 27 | CAP1 | | | ■ | | 6.90E-03 | 1.80E-02 | 7.60E-03 | NS | NS | NS | NS | NS |
| 28 | TINF2 | | | ■ | | 9.00E-03 | 2.20E-02 | NS | NS | NS | NS | NS | NS |
| 29 | NCBP1 | | | ■ | | 9.20E-03 | 2.20E-02 | 0.0344 | NS | NS | NS | NS | NS |
| 30 | SHROOM3 | ■ | | | | 0.0331 | 0.040 | NS | NS | NS | NS | NS | NS |
| 31 | THOC5 | | | ■ | | 0.0341 | NS | NS | NS | NS | NS | NS | NS |

| | | UniProt classification[4] | | | | | | | | | | | | |
|---|---|---|---|---|---|---|---|---|---|---|---|---|---|---|
| | Gene | Plasma membrane | Nucleus | Endoplasmic reticulum | Golgi apparatus | Lysosome | Endosome | Clathrin-dependent endocytosis | Transport vesicle | Cytoskeleton | Autophagy | Proteasomal/ ubiquination | Mitochondria | Cytosol |
| 1 | TRPC4AP | ■ | | | | | | | | | | ■ | | |
| 2 | PI4K2A | ■ | | | ■ | ■ | ■ | | ■ | | | | ■ | ■ |
| 3 | AP2A2 | ■ | | | | ■ | ■ | ■ | ■ | | | | | ■ |
| 4 | ITGA3 | ■ | | | | | | | | | | | | |
| 5 | RAB35 | ■ | | | | | ■ | ■ | | | | | | ■ |
| 6 | AP1G1 | ■ | | | ■ | ■ | ■ | ■ | ■ | | | | | ■ |
| 7 | VCP | | ■ | ■ | | ■ | | | | | | ■ | | ■ |
| 8 | PSMD1 | | ■ | | | ■ | | | | | | ■ | | ■ |
| 9 | NECAP2 | ■ | | | | | | ■ | | | | | | |
| 10 | PML | | ■ | ■ | | | ■ | | | | | | | ■ |
| 11 | RAB6B | | | | ■ | | | | ■ | | | | | ■ |
| 12 | AP2B1 | ■ | | | | ■ | ■ | ■ | | | | | | ■ |
| 13 | PSMD11 | | ■ | | | | | | | | | ■ | | ■ |
| 14 | RABEPK | | | | ■ | | ■ | | ■ | | | | | ■ |
| 15 | SEC24C | | | ■ | ■ | | | | | | | | | ■ |
| 16 | PSME1 | | ■ | | | | | | | | | ■ | | ■ |
| 17 | CUX1 | | ■ | | ■ | | | | | | | | | ■ |
| 18 | RAB8A | ■ | | | ■ | | ■ | | ■ | | | | | ■ |
| 19 | ATG2A | | ■ | | | | | | | | ■ | | | ■ |
| 20 | ARCN1 | | | ■ | ■ | | | | ■ | | | | | ■ |
| 21 | COPA | | | ■ | | | | | ■ | | | | ■ | ■ |
| 22 | PSMD7 | | ■ | | | | | | | | | ■ | | ■ |
| 23 | PREB | | ■ | ■ | ■ | | | | | | | | | |
| 24 | POLA2 | | ■ | | | | | | | | | | | ■ |
| 25 | TAP1 | | | ■ | | | | | | ■ | | | | |
| 26 | MYO9B | | | | | | | | | ■ | | | | ■ |
| 27 | CAP1 | ■ | | | | ■ | | | | ■ | | | | |
| 28 | TINF2 | | ■ | | | | | | | | | | | |
| 29 | NCBP1 | | ■ | | | | | | | | | | ■ | ■ |
| 30 | SHROOM3 | ■ | | | | | | | | ■ | | | | ■ |
| 31 | THOC5 | | ■ | | | | | | | | | | | |

#### Supplementary References


1 Mangat CS, Bharat A, Gehrke SS, Brown ED. Rank ordering plate data facilitates data visualization and normalization in high-throughput screening. J Biomol Screen. 2014;19:1314-20.

2 Lamb J, Crawford ED, Peck D, Modell JW, Blat IC, Wrobel MJ, et al. The Connectivity Map: using gene-expression signatures to connect small molecules, genes, and disease. Science. 2006;313:1929-35.

3 Subramanian A, Narayan R, Corsello SM, Peck DD, Natoli TE, Lu X, et al. A Next Generation Connectivity Map: L1000 Platform and the First 1,000,000 Profiles. Cell. 2017;171:1437-52 e17.

4 Broad Institute TCGA Genome Data Analysis Center: Analysis-ready standardized TCGA data from Broad GDAC Firehose 2016_01_28 run. Broad Institute of MIT and Harvard. Dataset 2016. DOI: 10.7908/C11G0KM9.

5 Cerami E, Gao J, Dogrusoz U, Gross BE, Sumer SO, Aksoy BA, et al. The cBio cancer genomics portal: an open platform for exploring multidimensional cancer genomics data. Cancer Discov. 2012;2:401-4.

6 Gao J, Aksoy BA, Dogrusoz U, Dresdner G, Gross B, Sumer SO, et al. Integrative analysis of complex cancer genomics and clinical profiles using the cBioPortal. Sci Signal. 2013;6:pl1.

7 Grossman RL, Heath AP, Ferretti V, Varmus HE, Lowy DR, Kibbe WA, et al. Toward a Shared Vision for Cancer Genomic Data. N Engl J Med. 2016;375:1109-12.

8 Fletcher A, Read ML, Thornton CEM, Larner DP, Poole VL, Brookes K, et al. Targeting Novel Sodium Iodide Symporter Interactors ADP-Ribosylation Factor 4 and Valosin-Containing Protein Enhances Radioiodine Uptake. Cancer Res. 2020;80:102-15.

9 Huang da W, Sherman BT, Lempicki RA. Systematic and integrative analysis of large gene lists using DAVID bioinformatics resources. Nat Protoc. 2009;4:44-57.

10 Huang da W, Sherman BT, Lempicki RA. Bioinformatics enrichment tools: paths toward the comprehensive functional analysis of large gene lists. Nucleic Acids Res. 2009;37:1-13.

11 Segura-Cabrera A, Tripathi R, Zhang X, Gui L, Chou TF, Komurov K. A structure- and chemical genomics-based approach for repositioning of drugs against VCP/p97 ATPase. Sci Rep. 2017;7:44912.

12 Skrott Z, Mistrik M, Andersen KK, Friis S, Majera D, Gursky J, et al. Alcohol-abuse drug disulfiram targets cancer via p97 segregase adaptor NPL4. Nature. 2017;552:194-9.

13 Ma X, Jin M, Cai Y, Xia H, Long K, Liu J, et al. Mitochondrial electron transport chain complex III is required for antimycin A to inhibit autophagy. Chem Biol. 2011;18:1474-81.

14 Ashoor R, Yafawi R, Jessen B, Lu S. The contribution of lysosomotropism to autophagy perturbation. PLoS One. 2013;8:e82481.

15 Tang C, Sun R, Wen G, Zhong C, Yang J, Zhu J, et al. Bromocriptine and cabergoline induce cell death in prolactinoma cells via the ERK/EGR1 and AKT/mTOR pathway respectively. Cell Death Dis. 2019;10:335.

16 Jo E, Li S, Liang Q, Zhang X, Wang H, Herbert TP, et al. Chronic activation of PPARalpha with fenofibrate reduces autophagic proteins in the liver of mice independent of FGF21. PLoS One. 2017;12:e0173676.

17 Joassard OR, Amirouche A, Gallot YS, Desgeorges MM, Castells J, Durieux AC, et al. Regulation of Akt-mTOR, ubiquitin-proteasome and autophagy-lysosome pathways in response to formoterol administration in rat skeletal muscle. Int J Biochem Cell Biol. 2013;45:2444-55.

18 Logan R, Kong A, Krise JP. Evaluating the roles of autophagy and lysosomal trafficking defects in intracellular distribution-based drug-drug interactions involving lysosomes. J Pharm Sci. 2013;102:4173-80.

19 Li X, Hu X, Wang J, Xu W, Yi C, Ma R, et al. Inhibition of autophagy via activation of PI3K/Akt/mTOR pathway contributes to the protection of hesperidin against myocardial ischemia/reperfusion injury. Int J Mol Med. 2018;42:1917-24.

20 Ertmer A, Huber V, Gilch S, Yoshimori T, Erfle V, Duyster J, et al. The anticancer drug imatinib induces cellular autophagy. Leukemia 2007;21:936-42.

21 Gafar AA, Draz HM, Goldberg AA, Bashandy MA, Bakry S, Khalifa MA, et al. Lithocholic acid induces endoplasmic reticulum stress, autophagy and mitochondrial dysfunction in human prostate cancer cells. PeerJ. 2016;4:e2445.

22 Zhang L, Hapon MB, Goyeneche AA, Srinivasan R, Gamarra-Luques CD, Callegari EA, et al. Mifepristone increases mRNA translation rate, triggers the unfolded protein response, increases autophagic flux, and kills ovarian cancer cells in combination with proteasome or lysosome inhibitors. Mol Oncol. 2016;10:1099-117.

23 Thomas HE, Zhang Y, Stefely JA, Veiga SR, Thomas G, Kozma SC, et al. Mitochondrial Complex I Activity Is Required for Maximal Autophagy. Cell Rep. 2018;24:2404-17 e8.

24 Deng L, Lei Y, Liu R, Li J, Yuan K, Li Y, et al. Pyrvinium targets autophagy addiction to promote cancer cell death. Cell Death Dis. 2013;4:e614.

25 Marois I, Cloutier A, Meunier I, Weingartl HM, Cantin AM, Richter MV. Inhibition of influenza virus replication by targeting broad host cell pathways. PLoS One. 2014;9:e110631.

26 Li H, Zhang Q, Yang X, Wang L. PPAR-gamma agonist rosiglitazone reduces autophagy and promotes functional recovery in experimental traumaticspinal cord injury. Neurosci Lett. 2017;650:89-96.

27 Fu Y, Huang Z, Hong L, Lu JH, Feng D, Yin XM, et al. Targeting ATG4 in Cancer Therapy. Cancers (Basel). 2019;11.
